# Pancreatic cancer-associated organ dysfunction promotes muscle autophagy and contributes to peripheral tissue wasting

**DOI:** 10.64898/2026.02.27.708635

**Authors:** Yetiş Gültekin, Sharanya Sivanand, Kian M. Eghbalian, Anna M. Barbeau, Keene L. Abbott, George Eng, Victoria L. Tavernier, Brian T. Do, Heaji Shin, Elif Özçelik, Sabrina Hu, Tenzin Kunchok, Millenia Waite, William M Rideout, Yiğit K. Kizlier, Daniel A. Sharygin, William A. Freed-Pastor, Tyler Jacks, Eileen White, Ömer H. Yilmaz, Jonathan A. Nowak, Brian M. Wolpin, Matthew G. Vander Heiden

## Abstract

Normal pancreas function supports both digestion and the hormonal regulation of whole-body metabolism. We find pancreatic ductal adenocarcinoma (PDAC) disrupts the normal function of the remaining pancreas, leading to altered systemic metabolism and peripheral tissue wasting that begins early in disease progression. Using mouse models of PDAC, we find small pancreas tumors lead to both endocrine and exocrine pancreatic dysfunction that results in systemic nutrient depletion and loss of both muscle and fat tissue. Providing free glucose in the diet that is absorbed despite pancreatic exocrine dysfunction causes hyperglycemia and blunts fat wasting without affecting muscle loss. Muscle mass can be restored by free dietary amino acids or pancreatic enzyme supplementation. Exocrine dysfunction causing reduced dietary protein digestion promotes muscle proteolysis and autophagy. Autophagy is a major driver of muscle wasting in PDAC, as muscle-specific deletion of the core autophagy gene *Atg7* also reduces muscle wasting. Disrupting muscle autophagy without restoring systemic nutrition slows tumor growth and improves survival of mice with PDAC. Tracing the fate of amino acids released from muscle of mice with PDAC shows redistribution to both tumor and host tissues. Notably, improving nutrition in mice with disrupted muscle autophagy promotes tumor growth. Together, the data argue that early peripheral tissue wasting associated with early pancreatic cancer is driven by altered normal pancreatic organ function that leads to reduced nutrition and enhanced muscle autophagy, releasing nutrients to support both tumor and host metabolism.

## Main

Pancreatic ductal adenocarcinoma (PDAC) is one of the most lethal cancers^1^. A hallmark of PDAC is cancer-associate tissue wasting with progressive loss of skeletal muscle and adipose tissue that is associated with poor patient outcomes including reduced treatment tolerance, diminished quality of life, and shortened survival^2,3^. The causes of tissue wasting in cancer are multifactorial, with increased energy expenditure, altered cytokine secretion, reduced appetite, and anhedonia all playing a role in some contexts^4,5,6^. However, PDAC-associated tissue wasting, particularly in early disease, remains incompletely understood.

Studies in both mouse pancreatic cancer models and patients have found that breakdown of muscle and adipose tissue begins early in disease progression^7,8,9^. There is evidence that patients experience both muscle and fat loss prior to diagnosis with tissue wasting being associated with both non-metastatic and metastatic PDAC^9^. Of note, muscle loss is more prominent in patients than adipose tissue loss, yet the presence of tissue wasting prior to diagnosis is not predicted by either tumor stage or the tumor location within the pancreas^9^. These data argue even solitary pancreatic tumors in patients can cause tissue wasting^4,5,9,10,11^. Additional metabolic changes also precede pancreatic cancer diagnosis in many patients, including new onset type 3c diabetes characterized by hyperglycemia with low insulin, as well as elevation of both host and microbiome-derived metabolites in blood^12,13,14,15,16^. At least some of these changes are reflective of increased muscle breakdown^4^, but the drivers of these systemic metabolic changes in early pancreatic cancer are not known.

Mouse models of PDAC recapitulate many of the metabolic features observed in humans including tissue wasting with early disease and evidence for increased circulating branched chain amino acids (BCAAs) associated with muscle wasting^4^. However, different from what is observed in humans, mice with PDAC exhibit more prominent adipose tissue wasting as well as hypoglycemia, which can be explained at least in part by pancreatic exocrine insufficiency^8^. While humans with PDAC can also experience exocrine insufficiency^17^, a relationship between exocrine insufficiency and muscle wasting has not been demonstrated and PDAC patients often present with hyperglycemia. Why mice develop hypoglycemia rather than hyperglycemia, and the underlying cause of early muscle wasting in PDAC, are both unknown.

Skeletal muscle wasting reflects a disruption of proteostasis, the dynamic balance between protein synthesis, degradation, and remodeling in muscle tissue. Anabolic pathways, including those regulated by mTORC1, myogenic regulatory factors, and hormone-sensitive transcription factors promote protein synthesis, ribosome biogenesis, and muscle fiber maintenance under nutrient– and amino acid-replete conditions^18,19,20,21,22,23,24,25^. Conversely, catabolic programs such as activation of the ubiquitin-proteasome system, the autophagy-lysosome pathway, and calcium-dependent proteases (calpains) mediate protein breakdown, particularly in response to starvation^23,26,27,28,29,30,31,32^. In cancer cachexia, anabolic signaling in muscle is suppressed while catabolic programs are hyperactivated, ultimately resulting in loss of muscle mass, with evidence that these same processes also contribute to early muscle wasting in PDAC^4^.

Muscle breakdown has systemic consequences and can be adaptive as part of the host response to starvation. Proteolysis releases amino acids and other metabolites into circulation, providing substrates for gluconeogenesis as well as amino acids for other tissues^33,34^. This response can also fuel cancers. For example, elevated circulating BCAAs can support tumor progression in lung cancer and promote epigenetic plasticity of PDAC tumor stroma^4,11,35^. Genetic studies found that the muscle-specific ubiquitin ligase MuRF1 can be involved in PDAC-associated muscle wasting; its inactivation reduces proteolysis, alters systemic and tumor metabolism, and delays tumor growth^36^. Likewise, disruption of host autophagy influences nutrient availability and tumor progression, suggesting that tissue breakdown can provide metabolic substrates to both tumor and host tissues^37,38^. In this study, we investigated how normal pancreatic function is affected by early PDAC, and how this might contribute to systemic metabolic changes, including muscle wasting, to affect tumor growth and host survival.

## Early pancreatic cancer is associated with systemic metabolic changes, including pancreatic endocrine dysfunction

The systemic consequences of early pancreatic cancer, including peripheral tissue wasting, are evident in mouse PDAC models. The *Kras^LSL-G12D/+^; Trp53^fl/fl^; Pdx1-Cre* (KP^−/−^C) PDAC model results in pancreatic tumor formation with reproducible and consistent kinetics, and exhibits muscle and fat loss early in disease progression that progresses to weight loss and reduced food intake (Extended Data Fig.1a-h)^8^. KP^−/−^C animals develop pancreatic lesions with abundant stroma and progress from precursor lesions with limited invasive cancer at 3-4 weeks-of age to end-stage disease by 10-12 weeks (Fig. 1a-b and Extended Data Fig.1c)^7,8,11^. Despite no differences in whole-body weight, food consumption, or liver weight between KP^−/−^C mice and littermate controls until several weeks after tumor onset (Extended Data Fig.1a-d), loss of both adipose and muscle tissues is evident by 3-5 weeks of age, with dramatic and rapid adipose tissue wasting and slower kinetics of muscle wasting (Fig. 1d-g and Extended Data Fig.1e-h). Of note, adipocyte size is smaller in the tumor-bearing mice (Fig. 1d), and histological analysis of skeletal muscle groups with fast-and or slow-twitch fibers shows smaller myofibers accompany tissue wasting (Fig. 1e-h) that progresses with advanced PDAC (Extended Data Fig.1i). A limitation of the KP^−/−^C model is that cancer develops in young mice and therefore tissue loss manifests largely as a failure to gain mass^39,40,41^. In the more widely studied *Kras^LSL-G12D/+;^ Trp53^LSL-R172H/+^; Pdx1-Cre* (KPC) model, stochastic tumor initiation can occur in young mice, but invasive PDAC typically develops in adult mice following *Trp53* loss of heterozygosity^7,39,42,43^ (Extended Data Fig.1j). By end point, KPC mice also exhibit near complete adipose tissue loss as well as muscle loss (Extended Data Fig.1k-q)^7,8,11,44^.

**Figure 1:**
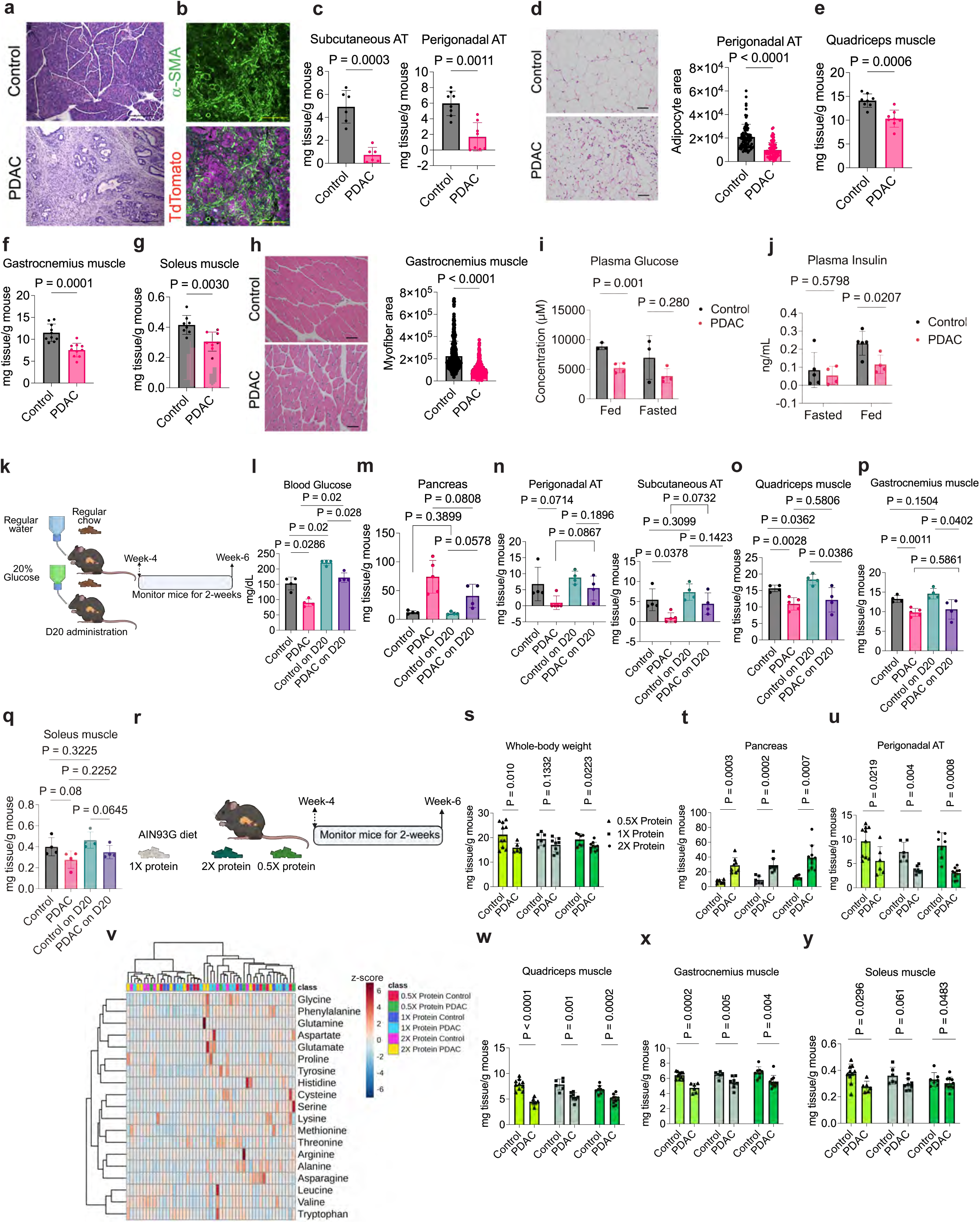
Early PDAC induces peripheral tissue wasting that is partially sensitive to dietary interventions. **a.** H&E staining of pancreas tissue from 6-week-old control and KP^−/−^C (PDAC) male mice. **b.** Immunofluorescence (IF) staining of pancreas tissue from 6-week-old KP^−/−^C; *Td-Tomato* male mice for α-smooth muscle actin (α-SMA) as a marker of stromal cells shown alone (top), and together Td-Tomato fluorescence from cancer cells (bottom). **c.** Subcutaneous and perigonadal adipose tissue (AT) weights normalized to whole-body weight of 6-week-old control and KP^−/−^C mice as indicated. (*n* = 6 control, *n* = 6 KP^−/−^C mice for subcutaneous AT; *n* = 8 control, *n* = 8 KP^−/−^C mice for perigonadal AT). **d.** H&E staining of perigonadal AT from 6-week-old control and KP^−/−^C mice. Adipocyte area was quantified for each image and is also shown. (*n* = 80 adipocytes counted from histology sections from *n* = 3 control and KP^−/−^C mice). **e-f.** Skeletal muscle groups with fast– and/or slow-twitch fibers: Quadriceps (mixed-significant fast-twitch) (e), gastrocnemius (fast-twitch) (f), and soleus (slow-twitch) skeletal muscle (g) weights normalized to whole-body weights of KP^−/−^C mice and littermate controls at 6-week-of age (*n* = 8 for quadriceps; *n* = 10 for gastrocnemius; *n* = 8 for soleus). **h.** H&E staining of gastrocnemius muscle from 6-week-old control and KP^−/−^C mice. Myofiber area was quantified for each image and is also shown. (*n* = 426 myofibers from *n* = 6 control and *n* = 541 myofibers from *n* = 6 KP^−/−^C mice). **i.** Plasma glucose levels from fed and 16h fasted 6-week-old control and KP^−/−^C mice (*n* = 3 control, *n* = 4 KP^−/−^C). **j.** Plasma insulin levels from fed and 16h fasted 6-week-old control and KP^−/−^C mice (*n* = 5 control, *n* = 4 KP^−/−^C). **k.** Experimental strategy for 20% D-glucose administration to 4-week-old mice in drinking water (D20) for 2-weeks. **l.** Plasma glucose levels after 2-weeks of water or D20 administration to control and KP^−/−^C mice as outlined in j (*n* = 4). **m-q.** Tissue weights of pancreas (m), perigonadal and subcutaneous ATs (n), quadriceps muscle (o), gastrocnemius muscle (p), and soleus muscle (q) normalized to whole-body weights of KP^−/−^C mice and littermate control mice after 2-weeks of exposure to D20 or water as outlined in k (*n* = 4). **r.** Experimental strategy for low and high protein diet administration to 4-week-old mice for 2-weeks. **s.** Whole-body weights of KP^−/−^C mice and littermate controls at after 2-weeks of diet administration as outlined in r (*n* = 10 control, *n* = 6 KP^−/−^C on 0.5X Protein diet; *n* = 6 control, *n* = 8 KP^−/−^C on 1X Protein diet; *n* = 7 control, *n* = 10 KP^−/−^C on 2X Protein diet). **t-u.** Tissue weights of pancreas (t) and perigonadal ATs (u) normalized to whole-body weights after 2-weeks of exposure to the specified diet as outlined in r. (*n* = 10 control, *n* = 6 KP^−/−^C on 0.5X Protein diet; *n* = 6 control, *n* = 8 KP^−/−^C on 1X Protein diet; *n* = 7 control, *n* = 10 KP^−/−^C on 2X Protein diet). **v**. Hierarchical clustering of plasma amino acid levels measured by GC-MS from KP^−/−^C mice and littermate controls fed the specified diets for 2-weeks as outlined in r. Columns of the heatmap were z-score normalized. (*n* = 9 control, *n* = 5 KP^−/−^C on 0.5X Protein diet; *n* = 8 control, *n* = 12 KP^−/−^C on 1X Protein diet; *n* = 7 control, *n* = 10 KP^−/−^C on 2X Protein diet). **w-y.** Tissue weights of quadriceps (w), gastrocnemius (x), and soleus muscle (y) normalized to whole-body weights of KP^−/−^C mice and littermate controls after 2-weeks of exposure to the specified diet as outlined in r. (*n* = 10 control, *n* = 6 KP^−/−^C on 0.5X Protein diet; *n* = 6 control, *n* = 8 KP^−/−^C on 1X Protein diet; *n* = 7 control, *n* = 10 KP^−/−^C on 2X Protein diet). Both male and female mice were used for all experiments except where specified. Statistical analyses were performed using one-way ANOVA with Tukey’s post hoc test for data in l-q. For other comparisons, unpaired two-sided *t*-tests were used. Data are presented as mean ± S.D., and *n* denotes the number of mice analyzed. Scale bars: 200 μm for panel a; 100 μm for panels d and g.

To explore other metabolic changes that accompany early tissue wasting in mice with PDAC, we collected plasma from fed and 16h-fasted KP^−/−^C and littermate control animals and conducted quantitative mass spectrometry to determine absolute levels of 112 metabolites^45,46^ (Extended Data Fig.2a). As expected, principal component analysis (PCA) revealed that metabolites measured in the fasted plasma samples cluster distinctly from those measured in the fed state from mice both with and without PDAC (Extended Data Fig.2b). Interestingly, hierarchical clustering was less effective at distinguishing each group (Extended Data Fig.2c). Consistent with previous reports^7,8^, circulating glucose levels were lower in the plasma of both fed and fasted PDAC-bearing mice (Fig. 1i), and BCAAs were moderately higher in the plasma of 16h-fasted PDAC mice (Extended Data Fig.2d). Serial blood sampling from KP^−/−^C mice showed blood glucose levels were lower in PDAC mice starting from 5-weeks and declined with PDAC progression (Extended Data Fig.2e). This finding is not observed in PDAC patients, who often present with new onset diabetes^47,48,49,50,51^. Consistent with the lack of hyperglycemia in KP^−/−^C mice, glycated hemoglobin levels were similar to littermate controls^52^(Extended Data Fig.2f). To assess endocrine pancreatic function in KP^−/−^C mice, we measured plasma insulin and glucagon and found similar glucagon levels as control mice in both the fed and fasted state (Extended Data Fig.2g), despite evidence of hypoglycemia (Fig. 1i and Extended Data Fig.2e). KP^−/−^C mice also have lower levels of insulin in the fed state (Fig. 1j). Histological examination of the pancreas at 6-weeks-of age from control and KP^−/−^C mice assessing the tumor marker CK19, glucagon, and insulin by immunohistochemistry showed reduced numbers of insulin-positive islets in tumor-bearing mice relative to controls despite extensive non-CK19-staining pancreatic tissue (Extended Data Fig.2h-i). These data are consistent with early loss of endocrine tissue in KP^−/−^C mice with PDAC and suggest that, similar to patients, pancreatic endocrine function is altered in mice with early PDAC even though hyperglycemia is not observed.

## Glucose supplementation selectively restores adipose tissue

The reduced insulin levels resulting from early PDAC-associated pancreatic endocrine dysfunction could contribute to increased lipolysis and adipose tissue wasting^53,54,55^. Lower circulating insulin also is consistent with activated hormone-sensitive lipase in adipose tissue of KP^−/−^C mice^8^. To further examine the relationship between hypoglycaemia and peripheral tissue loss, we explored whether increased glucose supplementation in the diet might affect the glucose metabolism phenotypes found in mice with early PDAC. Specifically, we supplemented the drinking water of KP^−/−^C mice with 20% (w/v) D-glucose (D20) starting at 4-weeks of age (Fig. 1k). This intervention increased blood glucose in mice with and without PDAC (Fig. 1l). We also assessed tissue wasting and tumor size in D20 exposed mice. D20 administration did not affect PDAC tumor size, but prevented adipose tissue wasting in KP^−/−^C mice (Fig. 1m-n). However, D20 administration had no effect on muscle loss (Fig. 1o-q). Despite reducing adipose tissue loss, long-term D20 exposure of KP^−/−^C mice did not impact survival of tumor-bearing mice and did not prevent muscle loss or weight loss at endpoint (Extended Data Fig.3a-g). Hence, increasing glucose in the diet can alter phenotypes in PDAC mouse models to better recapitulate findings observed in patients including higher blood glucose and less prominent adipose tissue loss, but does not impact tumor growth, tumor-associated muscle wasting, or survival.

Muscle is the major tissue responsible for protein storage in mammals, and the muscle proteome undergoes cycles of synthesis and breakdown in response to periods of feeding or fasting to maintain a supply of amino acids for the organism^29,56,53^. Thus, dietary amino acid deficiency leads to muscle protein breakdown that exceeds muscle protein synthesis and results in loss of muscle mass over time^54,59,60,61^. To test whether increasing protein in the diet can blunt loss of muscle tissue in PDAC mice, we evaluated the effects of isocaloric diets containing low protein (0.1g protein/g food), standard AIN-93G mouse diet protein levels (0.2g protein/g food), or high protein (0.4g protein/g food) (Fig. 1r). Short-term (2-week) exposure to these diets had no effect on body weight, PDAC tumor size, plasma amino acid levels, adipose tissue loss, or muscle loss in KP^−/−^C mice with PDAC (Fig. 1s-y). Given the adipose-sparing effects of D20, we next assessed a high-protein diet combined with D20 in KP^−/−^C PDAC mice over a two-week period. Similar to D20 alone, short-term high-protein diet combined with D20 did not alter tumor burden or muscle mass but restored adipose tissue (Extended Data Fig.3h-l). Longer-term exposure to different protein diets resulted in slightly smaller PDAC tumors and prolonged survival in KP^−/−^C mice fed a low-protein diet but had no effect on tissue wasting at endpoint (Extended Data Fig.3m-t). Notably, a high-protein diet failed to restore body weight, adipose tissue, or muscle mass in mice with end-stage PDAC and did not affect circulating amino acid levels (Extended Data Fig.3n-t). Together, these data indicate that dietary protein modulation does not prevent nor rescue PDAC-associated muscle loss.

## Pancreatic tumors cause exocrine dysfunction that promotes tissue wasting

To test whether peripheral tissue wasting in PDAC is influenced by tumor location, we implanted KP^−/−^C PDAC cells isolated from autochthonous mouse models either orthotopically into the tail of the pancreas or subcutaneously in syngeneic C57BL/6J mice (Fig. 2a and Extended Data Fig.4a). Histological analysis of PDAC tumors forming in either the flank or the pancreas showed evidence of fibrosis on trichrome staining and mucus production via alcian blue staining (Extended Data Fig.4b); however, only mice bearing orthotopic PDAC tumors exhibited early adipose and muscle tissue wasting (Fig. 2b-j and Extended Data Fig.4c-l), consistent with prior studies^5,8^. Of note, hypoglycemia was also only observed in mice with orthotopic PDAC tumors (Fig. 2d and Extended Data Fig.4e). To further test whether an adaptive immune system is required for these orthotopic PDAC associated phenotypes, we implanted murine KPC PDAC cells into the pancreas tail of Nu/J mice and still observed both muscle and adipose tissue wasting (Extended Data Fig.4m-r). These data argue that systemic metabolism changes associated with early PDAC do not require an adaptive immune system but are dependent on growth of a tumor in the pancreas.

**Figure 2:**
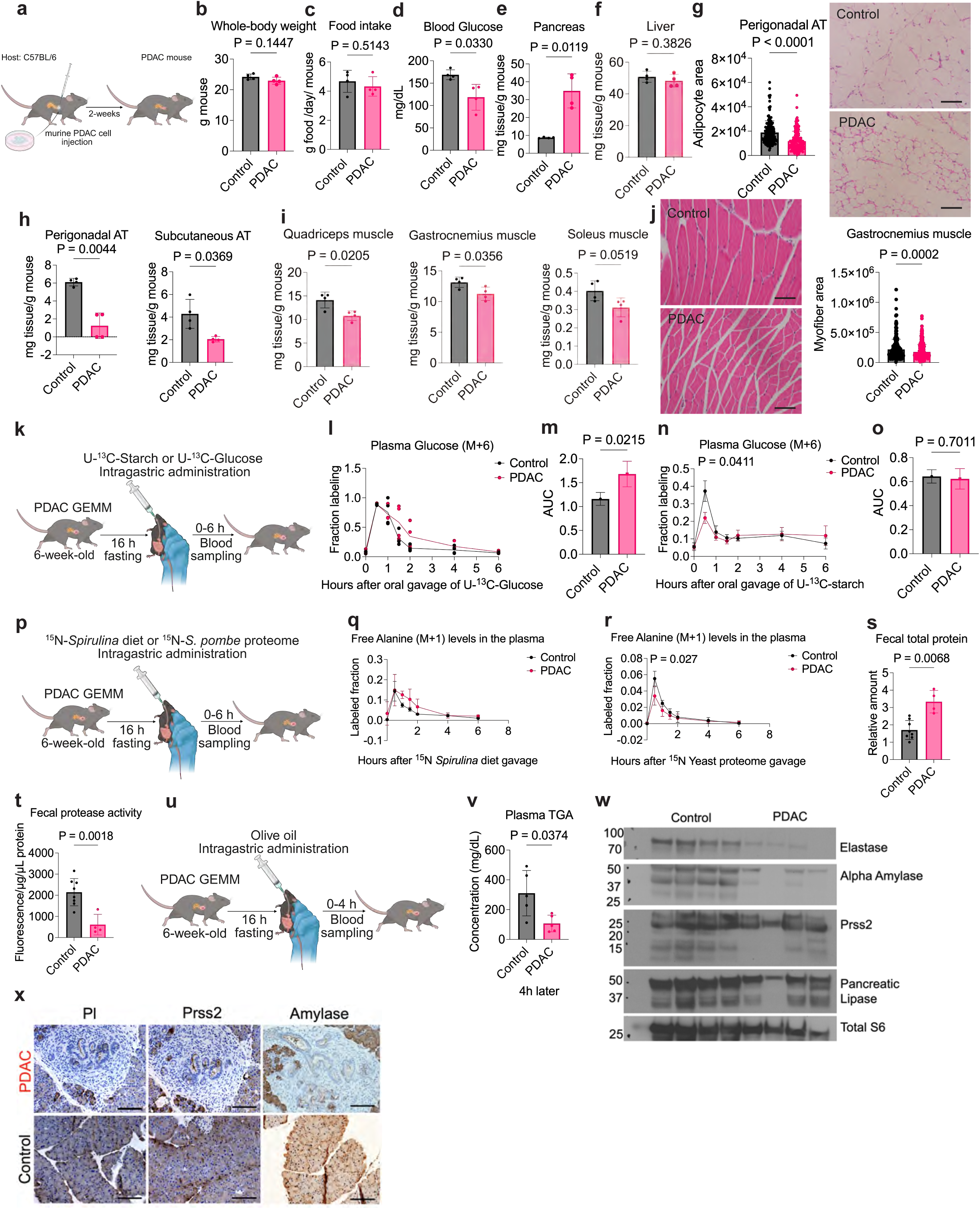
Mice with solitary PDAC tumors exhibit tissue wasting and pancreatic exocrine dysfunction. **a.** Schematic of experimental design to implant orthotopic murine KP^−/−^C PDAC cells into the tail of pancreas to form a tumor. **b-f.** Whole-body weight (b), food intake (c), blood glucose levels (d) and tissue weights normalized to whole-body weights of pancreas (e) and liver (f) 2-weeks after implantation of PBS (control) or pancreatic tumor cells (PDAC) into mice as outlined in a (*n* = 4). **g.** H&E staining of perigonadal AT histology from control and PDAC mice as outlined in a. Adipocyte area was quantified for each image and is also shown (*n* = 164 adipocytes counted from *n* = 4 control mice, and *n* = 143 adipocytes from *n* = 4 PDAC mice). **h-i.** Tissue weights of perigonadal and subcutaneous ATs (h) and quadriceps, gastrocnemius, and soleus muscle (i) normalized to whole-body weights from control and PDAC mice as outlined in a. **j.** H&E staining of gastrocnemius muscle from control and PDAC mice outlined in a. Myofiber area was also quantified for each image and is also shown. (*n* = 386 myofibers counted from *n* = 5 control and *n* = 365 myofibers from *n* = 5 PDAC mice). **k.** Experimental strategy for administration of uniformly labeled ^13^C-glucose (U-^13^C-glucose) or ^13^C-starch (U-^13^C-starch) by oral gavage to assess pancreatic exocrine function. **l.** Fraction of ^13^C-labeled glucose (M+6) measured in plasma over time after U-^13^C-glucose administration to 6-week-old control and KP^−/−^C (PDAC) mice as described in k. (*n* = 4). **m.** Area under curve (AUC) quantified for ^13^C-labeled glucose disposal for experiment shown in l. **n.** Fraction of ^13^C-labeled glucose (M+6) measured in plasma over time after U-^13^C-starch administration to 6-week-old control and KP^−/−^C (PDAC) mice as described in k. (*n* = 4). **o.** Area under curve (AUC) quantified for ^13^C-labeled glucose disposal for experiment shown in n. **p.** Experimental strategy for administration of ^15^N-*Spirulina* diet (contains labeled free amino acids) or ^15^N-labeled protein from *S. pombe* by oral gavage to assess pancreatic exocrine function. **q.** Fraction of ^15^N-labeled alanine (M+1) measured in plasma over time after ^15^N-*Spirulina* diet administration to 6-week-old control and KP^−/−^C (PDAC) mice as described in p (*n* = 4). **r.** Fraction of ^15^N-labeled alanine (M+1) measured in plasma over time after administration of ^15^N-labeled protein from *S. pombe* to 6-week-old control and KP^−/−^C (PDAC) mice as described in p (*n* = 4). **s-t.** Quantification of total protein amount (s) and protease activity in stool collected from 6-week-old control and KP^−/−^C mice 6h after ^15^N-*Spirulina* diet gavage as described in p. Protein measured in feces from KP^−/−^C mice was normalized to that measured in feces from control animals (*n* = 8 control, *n* = 4 KP^−/−^C PDAC). **u.** Experimental strategy for administration of olive oil by oral gavage to assess pancreatic exocrine function. **v.** Plasma triglyceride (TGA) concentration measured 4h after olive oil administration to 6-week-old control and KP^−/−^C mice as described in u (*n* = 5). **w.** Western blot analysis of the indicted pancreatic enzyme expression in pancreas tissue from 6-week-old control and KP^−/−^C mice (*n* = 4). Total S6 is shown as a loading control. **x.** Immunohistochemistry (IHC) staining for Pancreatic Lipase (Pl), Trypsin (Prss2), and Alpha-Amylase of pancreas tissue sections from 6-week-old control and KP^−/−^C mice (*n* = 4). Statistical analysis was performed using unpaired two-sided *t*-tests, data shown are mean ± S.D and *n* represents the number of mice analyzed. Scale bars: 100 μm for panel g and j; 200 μm for panel x.

To further test whether growth of a tumor in the pancreas is specifically required for tissue wasting, we implanted murine KPC PDAC cells subcutaneously, orthotopically, intrahepatically, or both orthotopically and intrahepatically (dual implantation) into syngeneic C57BL/6J mice (Extended Data Fig.5a). Only mice that had a PDAC tumor growing in the pancreas, with or without a tumor growing elsewhere, showed evidence of adipose and muscle wasting (Extended Data Fig.5b-g). We also assessed if the presence of any tumor in the pancreas causes this tissue wasting phenotype, or if the tumor has to be PDAC-derived. While hepatocellular carcinoma (HCC) cells exhibit a preference to form tumors in the liver and lung adenocarcinoma (LUAD) cells exhibit a preference to form tumors in the lung, these cells can form small tumors in the pancreas^62^(Extended Data Fig.5h). Tumors in the pancreas derived from either HCC or LUAD cells caused some degree of muscle and adipose tissue loss that corresponded with the size of the tumor growing in the pancreas (Extended Data Fig.5i-m). Implantation of primary rectal murine tumor organoids^63^ into the pancreas of syngeneic mice also led to tumor growth with loss of adipose and muscle tissue (Extended Data Fig.5n-s). Implanting the same primary rectal murine tumor organoids in the rectum or intrasplenically of syngeneic C57BL/6J mice led to tumor formation in the rectum and liver respectively without causing weight loss (Extended Data Fig.5t-x). After 6-weeks, mice with either rectal or liver tumors exhibited loss of adipose tissue, but muscle loss was not observed (Extended Data Fig.5y-z). We did not observe metastases outside of the intended site of implantation in any of the experiments involving HCC, LUAD, or rectal tumor organoids. In all cases, muscle loss was only observed when there was a tumor in the pancreas. Of note secreted factors have been reported to increase tissue wasting in other cancer contexts^64,61,65,66,67,68,69^ and in the case of rectal tumors, a secreted factor may play a role in adipose tissue loss^70,68,72^. However, the early tissue wasting associated with tumor growth in the pancreas appears to be a specific consequence of tumor growth in that organ.

Food intake was similar in control mice and mice with murine KP^−/−^C PDAC cells implanted in the pancreas at 2-weeks post-implantation (Fig. 2c), but a possible explanation for how tumor growth in the pancreas could cause tissue wasting is reduced nutrition secondary to pancreatic exocrine dysfunction^8^. To assess exocrine pancreatic function, we used ^13^C– or ^15^N-labeled nutrients that were incorporated into macromolecules that require pancreatic enzymes for digestion, or as free glucose or amino acids, and traced the appearance of label in the plasma at time points after oral delivery. Starch, a polymer of glucose, is the major carbohydrate source present in mouse chow. To examine whether starch digestion impairs availability of dietary glucose in PDAC mice, we administered uniformly labeled U-[^13^C]-glucose or uniformly-labeled-U-[^13^C]-starch by oral gavage to overnight-fasted control or KP^−/−^C mice (Fig. 2k). Upon administration of ^13^C-glucose, the control and PDAC mice exhibited a similar rate of appearance of ^13^C-glucose in circulation (Fig. 2l). However, the area under the curve (AUC) for glucose disposal from the plasma was higher in PDAC mice compared to control mice (Fig. 2m). These data suggest glucose clearance is slower in mice with PDAC, consistent with reduced insulin resulting from endocrine dysfunction (Fig. 1i). Consistent with impaired starch digestion, appearance of ^13^C-labeled glucose in the circulation was less in PDAC mice than control mice from oral delivery of ^13^C-labeled starch (Fig. 2n), and the AUC for glucose disposal was similar in the PDAC and control mice despite less overall glucose absorption (Fig. 2o). Furthermore, we challenged KP^−/−^C mice with a high starch diet compared to a control starch diet, and despite an increase in blood glucose observed in control mice, glucose levels remained low in the blood from KP^−/−^C mice (Extended Data Fig.6a-d). Together, these findings suggest that mice with PDAC have decreased ability to digest starch, and that this contributes to the hypoglycemia observed in mouse models of early PDAC when exposed to diets containing low levels of monosaccharides.

To assess whether pancreatic exocrine dysfunction in KP^−/−^C mice impairs digestion of dietary protein, we used an ^15^N-labeled diet derived from the algae *Spirulina*, which contains a mix of free elemental ^15^N-labeled amino acids and proteins containing ^15^N-labeled amino acids^73^, or protein with ^15^N-labeled amino acids (and no free amino acids) prepared from the yeast *S. pombe*. Following overnight fasting, we orally administered a solution of ^15^N-labeled *Spirulina* diet or ^15^N-labeled yeast protein and traced the appearance of ^15^N-labeled amino acids in blood (Fig. 2p). We detected appearance of multiple ^15^N-labeled amino acids in blood as early as 30 min post gavage from both diets (Fig. 2q-r and Extended Data Fig.6e-l). Of note, similar levels of ^15^N-labeled amino acids were observed in circulation from the ^15^N-labeled *Spirulina* diet, which contains free amino acids, while less ^15^N-labeled amino acids from ^15^N-labeled yeast protein were observed in KP^−/−^C mice compared to controls. These data are consistent with pancreatic exocrine dysfunction in PDAC mice that leads to impaired protein digestion. Also consistent with this phenotype, we found increased fecal protein content and decreased fecal protease activity in the stool of KP^−/−^C mice relative to controls (Fig. 2s-t).

To test whether pancreatic exocrine dysfunction also impairs dietary lipid metabolism in KP^−/−^C mice, we orally administered a defined dose of olive oil to overnight fasted KP^−/−^C mice (Fig. 2u). This olive oil challenge evoked higher levels of circulating triglycerides (TGAs) 4h post-gavage in control mice relative to KP^−/−^C mice (Fig. 2v). Finally, we examined the levels of several digestive enzymes including pancreatic elastase, alpha amylase, trypsin (Prss2), and pancreatic lipase (Pl) in pancreatic tissue from KP^−/−^C and control mice. We found compared to pancreatic tissue from control mice, tumor tissue from KP^−/−^C mice had lower pancreatic elastase and alpha amylase, while levels of Prss2 and PI were similar (Fig. 2w). Tissue IHC staining also showed a reduction of exocrine enzymes in tissue adjacent to pancreatic tumors in KP^−/−^C mice (Fig. 2x). These results suggest that exocrine dysfunction accompanies early PDAC, which results in reduced nutrient absorption from the diet and may contribute to tissue wasting.

To test whether systemic nutrient deficiency secondary to pancreatic exocrine dysfunctions contributes to tissue wasting in PDAC, we assessed whether pancreatic enzyme supplementation (PES) in the diet could reverse tissue wasting. For these studies, we used an orthotopic implantation model as these mice develop solitary pancreatic tumors without destruction of the remaining normal pancreas. 2-weeks after murine KP^−/−^C PDAC cell implantation into the pancreas tail, we fed mice a purified control diet or a purified diet containing 0.8% pharmaceutical grade PES and assessed the impact on exocrine function, tumor growth, and peripheral tissue wasting (Extended Data Fig.7a). We found 10-days on a PES diet restored pancreatic exocrine function in PDAC mice with decreased fecal protein content, increased fecal protease activity, decreased fecal lipid content and increased fecal lipase activity as well as increased fecal alpha amylase (Extended Data Fig.7b-f). We also assessed intestinal protease activity in saline flushed through the intestine and found that the PES diet enhanced intestinal protease activity in PDAC mice (Extended Data Fig.7g). Of note, exposure to a PES diet increased blood glucose levels and attenuated both adipose and muscle tissue wasting in PDAC mice without affecting body weight, food intake, or size of the liver or pancreas (Extended Data Fig.7h-n). We also delivered 2% PES by oral gavage (or PBS as a control) daily for 2-weeks to KP^−/−^C animals on a normal chow diet (Extended Data Fig.7p). 2-weeks of PES via oral gavage caused similar findings with reduced muscle and fat wasting (Extended Data Fig.7q-z). Importantly, the PES-containing purified diet did not influence food intake, change body weight or other organ sizes, or affect pancreatic exocrine functions when administered to non-tumor-bearing C57BL/6J mice for 4-weeks (Extended Data Fig.8). Taken together, these results confirm that pancreatic exocrine insufficiency is a contributor to peripheral tissue wasting in mice with early PDAC.

## Autophagy is increased in muscle of PDAC mice

To assess the impact of pancreatic exocrine dysfunction on systemic nutrient availability as a possible contributor to muscle wasting, we isolated plasma and interstitial fluid from the muscle of PDAC-bearing mice. We then conducted quantitative mass spectrometry to determine the absolute levels of 112 metabolites in muscle interstitial fluid (MIF) and plasma (Fig. 3a). Notably, we found that numerous metabolites, including several essential amino acids as well as glucose, were present at lower concentrations in MIF compared to plasma of PDAC-bearing mice (Fig. 3b), and hierarchical clustering showed PDAC MIF samples clustered separately from the plasma samples (Fig. 3c). We also infused [U-^13^C]-glucose into conscious unrestrained KP^−/−^C and littermate control mice to assess glucose fate in the muscle of these animals (Extended Data Fig.9a). [U-^13^C]-glucose was infused for 6hrs, as prior work showed this time was sufficient to approach steady state labeling in muscle^57,74^. We also confirmed that plasma ^13^C-glucose labeling approaches steady-state over this infusion period (Extended Data Fig.9b-c). ^13^C-labeling of lactate, pyruvate, serine, alanine, and TCA cycle metabolites was similar or slightly higher in the muscle of PDAC mice despite similar metabolite levels as littermate controls (Extended Data Fig.9d-s). These findings are consistent with a more catabolic state in muscle of PDAC mice, and fit with a recent report showing elevated glucose turnover in skeletal muscle of cachectic mice with C26-cell derived tumors^75^.

**Figure 3:**
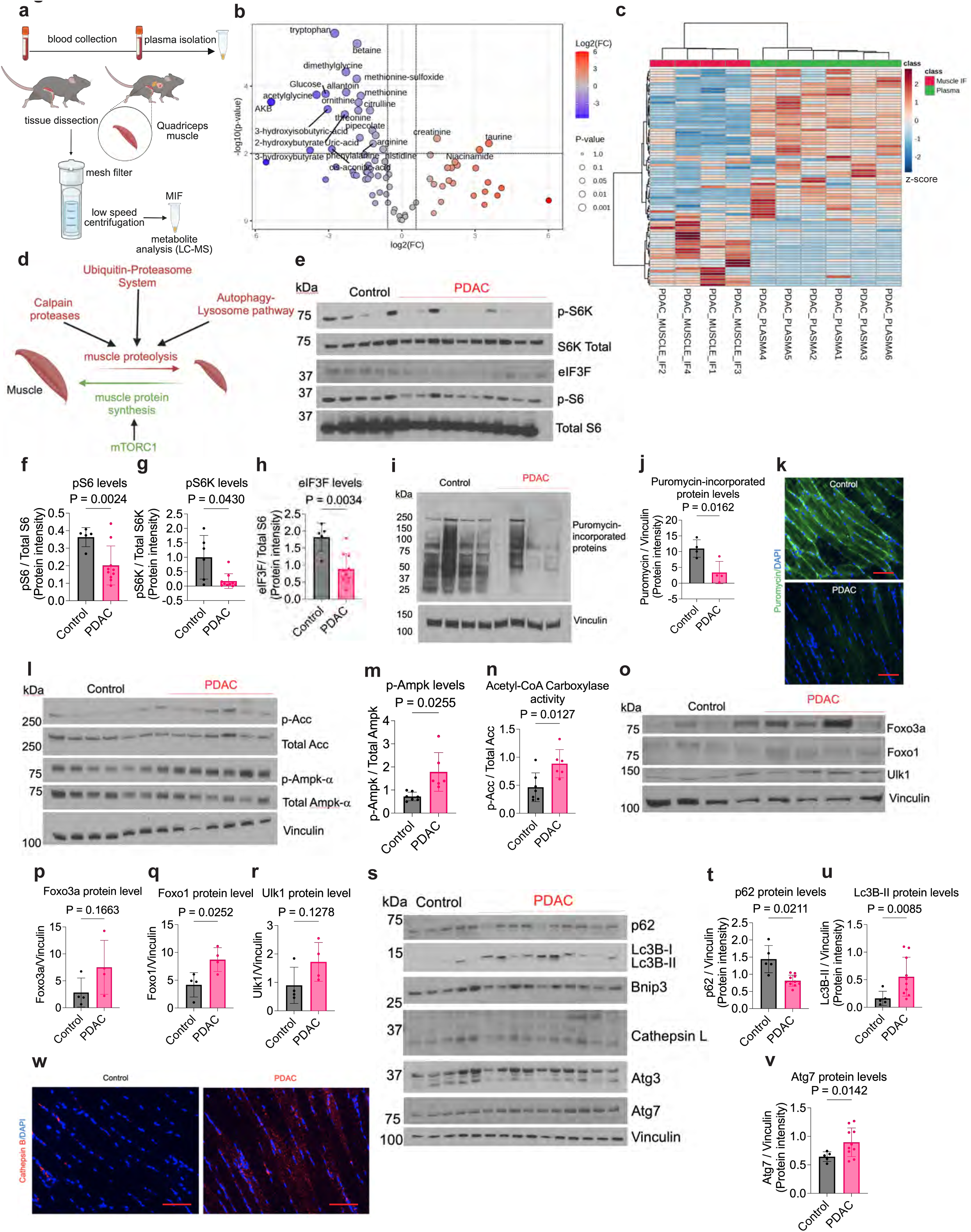
Muscle autophagy is triggered by reduced nutrition in PDAC mice. **a.** Schematic of experimental design to collect plasma and muscle interstitial fluid (MIF) from 6-week-old KP^−/−^C mice with PDAC. **b.** Volcano plot depicting the log_2_ fold change in metabolite concentration measured in MIF relative to plasma from KP^−/−^C mice as described in a (*n* = 4 MIF; *n* = 6 plasma). A fold change of 2 and a raw P-value of 0.01 assuming unequal variance were used to select significantly altered metabolites labeled. **c.** Hierarchical clustering of levels of 112 metabolites measured in MIF and plasma from PDAC-bearing mice as described in a (*n* = 4 MIF; *n* = 6 plasma). Columns of the heatmap were z-score normalized. **d.** Schematic depicting regulation of muscle protein synthesis and proteolysis. **e.** Western blot analysis of mTORC1 signaling proteins and muscle specific eIF3F in gastrocnemius muscle tissue from 6-week-old control and KP^−/−^C (PDAC) mice. Total S6 levels are included as a loading control. **f-h.** Quantifications of pS6 (f), pS6K (g), and eIF3F (h) levels from the western blot shown in e. (*n* = 5 control, *n* = 10 KP^−/−^C PDAC). **i.** Western blot analysis of puromycin-incorporated proteins to assess new protein synthesis in gastrocnemius muscle tissue from 6-week-old control and KP^−/−^C mice. Vinculin is included as a loading control (*n* = 4). **j.** Quantification of puromycin-incorporated protein levels from the western blot shown in i. **k.** IF to assess puromycin (green) staining in gastrocnemius muscle tissue from 6-week-old control and KP^−/−^C mice. **l.** Western blot analysis of Ampk signaling proteins in gastrocnemius muscle tissue from 6-week-old control and KP^−/−^C mice. Vinculin is included as a loading control (*n* = 7 control, *n* = 6 KP^−/−^C PDAC). **m-n.** Quantification of p-Ampk (m) and Acetyl-CoA carboxylase (n) from the western blot shown in l. **o.** Western blot analysis of Foxo1, Foxo3a, and Ulk1 protein in gastrocnemius muscle tissue from 6-week-old control and KP^−/−^C mice. Vinculin is included as a loading control (*n* = 4). **p-r.** Quantification of Foxo1 (p), Foxo3a (q), and Ulk1 (r) levels from the western blot shown in o. **s.** Western blot analysis of proteins in the autophagy-lysosome proteolysis pathway in gastrocnemius muscle tissue from 6-week-old control and KP^−/−^C mice. Vinculin is included as a loading control (*n* = 5 control and *n* = 10 KP^−/−^C). **t-v.** Quantification of p62 (t), Lc3B-II (u), and Atg7 (v) levels from the western blot shown in s. **w.** IF to assess Cathepsin B staining in gastrocnemius muscle tissue from 6-week-old control and KP^−/−^C mice. Both male and female mice were used for the experiments shown. Statistical analysis was performed using unpaired two-sided *t*-tests, data are mean ± S.D and *n* represents the number of mice analyzed. Scale bars: 100 μm.

The muscle proteome is regulated by coordinated control of protein synthesis and degradation in response to physiological and pathological signals^56,56^. Protein synthesis is controlled by mTORC1, which integrates nutrient, growth factor, hormones, and mechanical cues^75,77,78^ while proteolysis occurs through the ubiquitin-proteasome and autophagy-lysosome systems, as well as via activation of calcium-dependent proteases (Fig. 3d)^30,31,32^. To assess if these regulatory networks are disrupted in muscle from mice with early PDAC, we first assessed mTORC1 activity^78^, and found decreased phosphorylation of the mTORC1 substrates S6 and S6K1 in muscle from PDAC mice (Fig. 3e-g). In muscle, eIF3F links mTORC1 to the translation machinery^18,15,21^. During muscle atrophy, eIF3F is degraded and uncouples mTORC1 from translation and suppresses muscle growth^21^. Gastrocnemius muscle of KP^−/−^C mice showed diminished levels of eIF3F relative to control mice (Fig. 3e,h). To test whether decreased activity of eIF3F-mTORC1 translation machinery is associated with less protein synthesis in the muscle of KP^−/−^C mice, we measured puromycin incorporation into muscle protein^81^. Muscle from control mice showed higher puromycin incorporation into protein than was observed in muscle from KP^−/−^C mice (Fig. 3i-k). These data suggest reduced mTORC1 activity and protein synthesis in the muscle of PDAC-bearing mice.

To assess possible causes of decreased muscle mTORC1 activation and protein synthesis in mice with early PDAC, we considered that reduced nutrition resulting from exocrine dysfunction might result in increased AMP-activated protein kinase (AMPK) signaling, as under starvation AMPK inhibits mTORC1^30,80,83^. Of note, levels of active (phosphorylated) AMPK were higher in the muscle of KP^−/−^C mice, as were levels of phosphorylated acetyl-coA carboxylase (Acc), a direct AMPK target^82,83^(Fig. 3l-n). We also observed increased protein levels of Ulk1, Foxo1, and Foxo3a in the muscle of KP^−/−^C mice, which are all responsive to AMPK signaling^81^(Fig. 3o-r). Overall, these data are consistent with reduced nutrition leading to decreased protein synthesis in the muscle of mice with early PDAC.

AMPK-mediated activation of Ulk1 and Foxo1/3a, as well as loss of mTORC1 signaling, can promote the ubiquitin-proteasome system (UPS) and lysosome-autophagy pathways to promote protein breakdown as part of a response to starvation^27,26,80,81,84^. To assess whether UPS activity is increased in KP^−/−^C mouse muscle tissue, we used a luminescence reporter assay for proteosome activity as well as immunoblotting for various ubiquitin-E3 ligases associated with muscle protein catabolism. Of note, proteosome activity was similar in muscle isolated from control and KP^−/−^C mice at both earlier and later time points during disease progression (Extended Data Fig.10a-b). Levels of the UPS-related proteins including Ubcj1, Rpt3, and Alpha-7 were also similar in the gastrocnemius muscle of KP^−/−^C mice compared to control mice (Extended Data Fig.10c-f). However, levels of poly-ubiquitinylated proteins were higher in gastrocnemius muscle of KP^−/−^C mice compared to controls (Extended Data Fig.10g-h). To explore whether Calpain protease activity is increased in muscles of mice with early PDAC, we assessed both Calpain activity and levels and found no difference in gastrocnemius muscle from KP^−/−^C and littermate control mice (Extended Data Fig.10i-k). However, evidence for increased autophagy was evident in muscle of KP^−/−^C mice (Fig. 3s). This includes decreased levels of the autophagy adaptor protein p62, higher levels of Atg7, and increased conversion of Lc3B-I to Lc3B-II (Fig. 3s-v). Elevated levels of the lysosomal protease Cathepsin B were also observed in muscle from KP^−/−^C mice (Fig. 3w). These data are consistent with increased autophagy potentially contributing to increased muscle loss in mice with early PDAC.

To examine how PDAC affects lysosomal metabolism in muscle, we generated *Ckm-Cre; Rosa26-LSL–TMEM192-3xHA* (muscle LysoTag) mice^85,86^, confirmed increased muscle LysoTag-labeled lysosomes in quadriceps after fasting, and validated efficient lysosome enrichment by Lyso-IP (Extended Data Fig.11a-c). Next, to assess how lysosomal metabolites change in muscle of PDAC mice, we implanted murine PDAC cells or PBS as a control into the pancreas of muscle LysoTag mice, and 4-weeks post-implantation isolated quadriceps muscle for Lyso-IP and metabolite analysis (Extended Data Fig.11d). Similar to KP^−/−^C mice with PDAC (Fig. 3w), muscle from mice with implanted KPC PDAC tumors also contained more lysosomes (Extended Data Fig.11e). Levels of tryptophan, glutamine, methionine, and asparagine were higher in muscle lysosomes of PDAC mice relative to controls (Extended Data Fig.11f). These data are consistent with reduced essential amino acids measured in MIF, as reduced amino acid availability is expected to lead to increased autophagy and increased amino acid levels in lysosomes.

Dietary restriction of an essential amino acid can cause muscle wasting^61,87^ and has been reported to cause adipose tissue wasting even without changes in caloric intake^59,61^. To investigate how selective essential amino acid deficiencies influence tissue wasting, we examined C57BL/6J mice fed a nutritionally complete elemental amino acid diet, or a diet lacking either the essential amino acid tryptophan, methionine, or lysine, or lacking the essential branched-chain amino acids (valine, leucine, and isoleucine), or lacking both serine and glycine (non-essential amino acids) for four weeks (Extended Data Fig.12a). Restriction of any of the essential amino acids was sufficient to induce whole-body weight loss driven largely by adipose tissue wasting, and under some conditions, decreased pancreas and liver mass (Extended Data Fig.12b-f). However, only dietary tryptophan restriction led to a significant reduction in skeletal muscle mass in this experiment (Extended Data Fig.12g-i). These data confirm that reduced intake of essential amino acids can promote tissue wasting.

To test whether increasing dietary free amino acid availability could mitigate muscle wasting in PDAC, we exposed mice to purified diets formulated with elemental amino acids in which intact proteins were replaced with a matched composition of free amino acids, thereby bypassing any effects of pancreatic exocrine dysfunction on protease-dependent digestion of dietary protein. PDAC mice were maintained on these diets long term and analyzed at endpoint (Extended Data Fig.13a). A diet enriched in free amino acids (2X FAA) increased whole-body weight in KP^−/−^C mice with PDAC relative to KP^−/−^C mice with PDAC fed a diet with lower FAA content (Extended Data Fig.13b). Nonetheless, the 2X FAA diet did not affect tumor size and liver mass or mitigate adipose wasting (Extended Data Fig.13c-e). The 2X FAA diet also limited loss of skeletal muscle mass and extended survival in mice with KP^−/−^C PDAC (Extended Data Fig.13f,g). These findings further support that exocrine dysfunction leading to reduced availability of dietary amino acids contributes to PDAC-associated muscle wasting.

## Preventing muscle autophagy protects against tissue wasting and prolongs survival in PDAC-bearing mice

To test whether autophagy contributes to early muscle wasting in PDAC, we used a mouse model that enables independent genetic manipulation of tumor cells and skeletal muscle. Specifically, a Flp-recombinase-driven PDAC model was crossed with *Atg7^fl/fl^*mice who also have a *Ckm-Cre* allele to selectively delete the essential autophagy gene *Atg7* in skeletal muscle^88,87^(Fig. 4a). Similar to the KP^−/−^C model, the Flp recombinase-activated PDAC model (KP^−/−^F) involves *Pdx1-P2A-FlpO, Kras^Frt-stop-Frt-G12D^* (*Kras^FSF-G12D^*), and *Tp53^Frt/Frt^* alleles that lead to expression of mutant *Kras* and loss of *p53* in the pancreas to cause KP^−/−^F PDAC^89,90^. We confirmed that *Atg7* is deleted only in muscle of *Flp*-driven KP^−/−^F mice with *Ckm-Cre* and *Atg7^fl/fl^* alleles (Fig. 4b). Loss of *Atg7* prevented autophagy in muscle and reduced muscle wasting in PDAC mice (Fig. 4c-f). Interestingly, muscle specific *Atg7* deletion also rescued body weight loss, increased blood glucose and insulin, and reduced adipose tissue wasting in KP^−/−^F PDAC-bearing mice, while leading to slightly improved food intake (Fig. 4g-j). These effects may be due in part to loss of muscle autophagy reducing KP^−/−^F PDAC growth (Fig. 4n-p). Muscle specific *Atg7* depletion also improved survival of KP^−/−^F PDAC mice (Fig. 4q).

**Figure 4:**
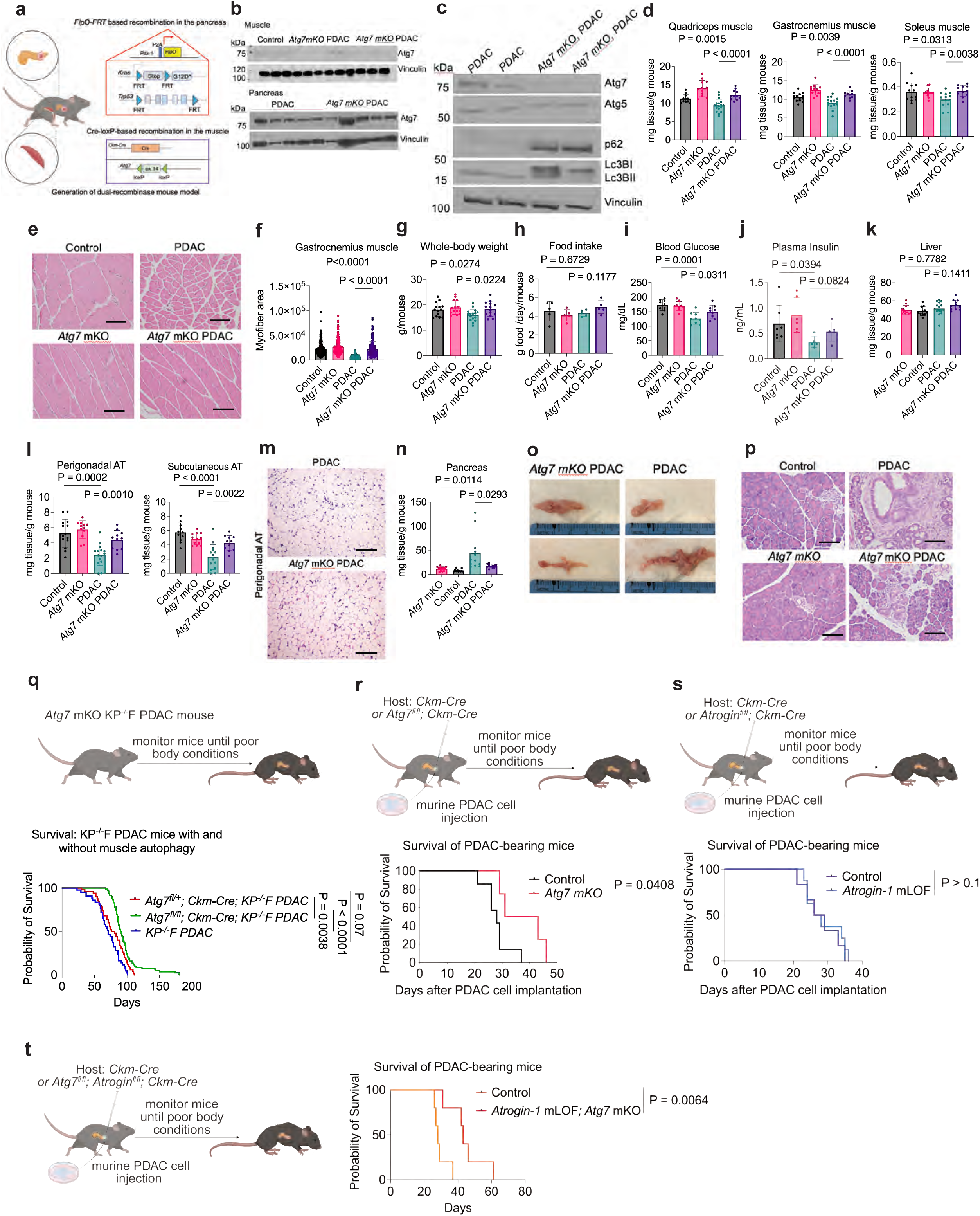
Increased muscle autophagy supports PDAC tumor growth. **a.** Schematic showing experimental approach to disrupt autophagy in the muscle of mice with genetically engineered KP^−/−^F PDAC tumors. **b.** Western blot assessing Atg7 protein in muscle tissue from 6-week-old control (*Pdx-1-P2A-FlpO; Trp53^Frt/Frt^; Ckm-Cre*) or *Atg7* muscle knock-out mice (*Ckm-Cre*; *Atg7^fl/fl^*: referred to as *Atg7* mKO), without or with (*Pdx-1-P2A-FlpO; Kras^FSF-G12D/+^; Trp53^Frt/Frt^*) KP^−/−^F PDAC as indicated (*n* = 3). Also shown is western blot for Atg7 in pancreas tissue from control and *Atg7* mKO mice with KP^−/−^F PDAC (*n* = 5). **c.** Western blot for Atg7, Atg5, p62, Lc3B-I, and Lc3B-II in gastrocnemius muscle from 6-week-old control and *Atg7* mKO mice with KP^−/−^F PDAC (*n* = 2). Vinculin is blotted as a loading control. **d.** Gastrocnemius, quadriceps, and soleus skeletal muscle weights normalized to whole-body weights from 6-week-old control or *Atg7* mKO mice, without or with KP^−/−^F PDAC as indicated (*n* = 14 control, *n* = 12 *Atg7* mKO, *n* = 16 KP^−/−^F PDAC, and *n* = 11 *Atg7* mKO KP^−/−^F PDAC). **e.** H&E staining of gastrocnemius muscle from 6-week-old control or *Atg7* mKO mice, without or with KP^−/−^F PDAC as indicated. **f.** Myofiber area quantification from histology shown in e. (*n* = 230 control, *n* = 166 *Atg7* mKO, *n* = 477 KP^−/−^F PDAC, and *n* = 186 *Atg7* mKO KP^−/−^F PDAC from *n* = 4 mice for each group). **g-l.** Whole-body weight (*n* = 14 control, *n* = 14 *Atg7* mKO, *n* = 16 KP^−/−^F PDAC, and *n* = 14 *Atg7* mKO KP^−/−^F PDAC, 6-week-old) (g), food intake (*n* = 5) (h), blood glucose (*n* = 9) (i), plasma insulin (*n* = 7 control, *n* = 6 *Atg7* mKO, *n* = 5 PDAC, and *n* = 5 *Atg7* mKO KP^−/−^F PDAC) (j), liver weight (*n* = 11 control, *n* = 12 *Atg7* mKO, *n* = 15 KP^−/−^F PDAC, and *n* = 11 *Atg7* mKO KP^−/−^F PDAC) (k), and perigonadal and subcutaneous ATs weights (*n* = 14 control, *n* = 12 *Atg7* mKO, *n* = 16 KP^−/−^F PDAC, and *n* = 11 *Atg7* mKO KP^−/−^F PDAC) (l) normalized to whole-body weights of 6-week-old control or *Atg7* mKO mice, without or with KP^−/−^F PDAC as indicated. **m.** H&E staining of perigonadal AT from control and *Atg7* mKO mice with KP^−/−^F PDAC. **n.** Pancreas tissue weights normalized to whole-body weights from 6-week-old control or *Atg7* mKO mice, without or with KP^−/−^F PDAC as indicated (*n* = 11 control, *n* = 12 *Atg7* mKO, *n* = 11 KP^−/−^F PDAC, and *n* = 12 *Atg7* mKO KP^−/−^F PDAC). **o.** Gross images of pancreas from 6-week-old control and *Atg7* mKO mice with KP^−/−^F PDAC as indicated (*n* = 2). **p.** H&E staining of pancreas tissue from 6-week-old control or *Atg7* mKO mice, without or with KP^−/−^F PDAC as indicated. **q.** Survival of control mice with KP^−/−^F PDAC (*Pdx-1-P2A-FlpO; Kras^FSF-G12D/+^; Trp53^Frt/Frt^*, referred to here as KP^−/−^F PDAC), or the same mice with a *Ckm-Cre* and one (*Atg7^fl/+^*) or two (*Atg7^fl/fl^*) *Atg7^fl^* alleles as indicated (*n* = 43 PDAC, *n* = 50 *Atg7^fl/+^*, and *n* = 82 *Atg7^fl/fl^*). **r.** Survival of control or *Atg7* mKO mice following orthotopic implantation of murine KPC PDAC cells into pancreas (*n* = 6). **s.** Survival analysis of control mice and mice with muscle-specific loss of function of *Atrogin-1* (*Atrogin-1^fl/fl^; Ckm-Cre: Atrogin-1* mLOF) following orthotopic pancreatic implantation of KPC PDAC cells (*n* = 6). **t.** Survival analysis of control mice and mice with combined muscle-specific *Atrogin-1* loss of function and *Atg7* deletion (*Atrogin-1* mLOF; *Atg7* mKO) following orthotopic KPC PDAC implantation (*n* = 6). Both male and female mice were used in experiments. Statistical analyses were performed using one-way ANOVA with Tukey’s post hoc test for data in panels c-j and m-n. P-values on survival experiments were calculated with Gehan-Breslow-Wilcoxon test. Data are presented as mean ± S.D., and *n* denotes the number of mice analyzed. Scale bars: 100 μm.

To test the impact of disrupting autophagy in the muscle on PDAC tumor growth in another mouse model, we implanted murine KPC PDAC cells into the flank or the pancreas of *Atg7* muscle knock-out (*Atg7* mKO) mice or control animals (Extended Data Fig.14). At 4-weeks after KPC PDAC cell implantation in the pancreas, control PDAC mice had larger tumors and more muscle and adipose tissue wasting than *Atg7* mKO mice (Extended Data Fig.14a-f). *Atg7* mKO mice with orthotopically implanted KPC PDAC tumors also had improved survival relative to control mice (Fig. 4r). Of note, flank tumors derived from KPC PDAC tumor cells exhibited similar growth in control and *Atg7* mKO mice, and significant differences in the size of organs was not observed between these groups of mice (Extended Data Fig.14g-m). Taken together, these data support a model where tumor growth in the pancreas leads to pancreatic exocrine dysfunction-associated increases in muscle autophagy that contribute to tissue wasting.

While autophagy appears to be a major driver of muscle wasting in the PDAC models tested, increased proteasome-mediated protein degradation has been reported in skeletal muscle of tumor-bearing hosts^36,87^. Various ubiquitin-E3 ligases are known to be involved in muscle protein degradation such as E3 ligases F-Box Protein 32 (Fbxo32/Atrogin-1) and muscle RING finger protein 1 (MuRF1/Trim63)^32^. *Atrogin*-*1* has been linked to cancer-associated muscle wasting^27,91^. To test a role for Atrogin-1 in PDAC-associated muscle wasting, we crossed *Atrogin-1^fl/fl^* mice to animals with a *Ckm-Cre* allele to enable muscle specific loss of functional *Atrogin-1* (*Atrogin*-*1* mLOF) (Extended Data Fig.15a-b). To determine whether Atrogin-1 is required for PDAC-associated muscle wasting, we orthotopically implanted murine KPC PDAC cells into the pancreas of control or *Atrogin-1* mLOF mice (Extended Data Fig.15c) and found muscle specific loss of Atrogin-1 function only slightly reduced PDAC tumor size while reducing both adipose tissue and muscle wasting (Extended Data Fig.15d-g). This mild effect on tumor and organ size led to no difference in survival between control and *Atrogin*-*1* mLOF PDAC-bearing mice (Fig. 4s). We also crossed *Atrogin*-*1* mLOF mice to *Atg7* mKO mice to disrupt both autophagy and Atrogin-1-mediated proteolysis in muscle (*Atg7* mKO; *Atrogin*-*1* mLOF). When KPC PDAC cells are implanted into the pancreas of these mice, they survive longer than PDAC-bearing mice with intact muscle Atrogin-1 and Atg7 (Fig. 4t). Taken together, these results confirm that genetic disruption of muscle breakdown pathways, most prominently autophagy, leads to reduced PDAC tumor growth and tissue wasting, as well as improved survival.

## ^15^N-Spirulina labeling demonstrates redistribution of muscle-derived free amino acids into both PDAC tumors and host organs

The finding that genetically blocking muscle wasting impedes PDAC tumor growth and improves survival suggests muscle breakdown may provide amino acids to support the tumor. To begin to test this, we conducted quantitative metabolomic analysis of plasma samples from mice with KP^−/−^F PDAC with or without *Atg7* deletion in muscle (Extended Data Fig.16a); however, hierarchical clustering revealed that metabolites measured in the plasma samples did not cluster distinctly (Extended Data Fig.16b). We next assessed whether nutrients released from host organ stores contribute to tissue biomass in mice with PDAC. For these experiments, we fed mice an ^15^N-labeled *Spirulina* diet and found this resulted in 70-80% enrichment of ^15^N-labeled free amino acids in plasma, as well as in plasma proteins after two weeks of diet exposure (Fig. 5a-c). We also assessed labeling of free amino acids and amino acids in protein from multiple mouse tissues after two weeks of diet exposure. Up to 80% of the free amino acids in many tissues were labeled, but different protein labeling was observed across organs, with more of the proteome ^15^N-labeled in kidney, colon, small intestine, liver, pancreas, spleen, and adipose tissues and less of the proteome ^15^N-labeled in brain and muscle tissues (Fig. 5d-e). However, when mice were exposed to ^15^N-labeled *Spirulina* diet for two weeks and then switched to regular non-labeled chow diet for four weeks there was a different distribution of ^15^N-labeled amino acids in both free pools and tissue protein, with much lower labeling of circulating amino acids and higher enrichment of ^15^N-label in the proteome of brain and muscle (Fig. 5f-j). To assess how tissue labeling changes in mice with PDAC, with or without genetic impairment of muscle breakdown, we fed control and *Atg7* mKO mice the ^15^N-labeled *Spirulina* diet for two weeks, then shifted animals to unlabeled diet and orthotopically implanted murine KPC PDAC cells into the pancreas. We allowed tumors to grow in mice for four weeks on regular chow diet and then assessed ^15^N-labeled amino acids in plasma and tissues (Fig. 5k). Interestingly, similar labeling of free amino acids as well as amino acids in the proteome of plasma, PDAC tumors, and other organs was observed in mice with or without disruption of muscle autophagy (Fig. 5l-q and Extended Data Fig.17). These data suggest that muscle does not uniquely supply amino acids to the tumor or to any other tissue in the mouse. It also argues that wasting of multiple tissues is likely involved in the redistribution of nutrients across the animal as a result of early PDAC-associated exocrine dysfunction leading to reduced nutrition.

**Figure 5:**
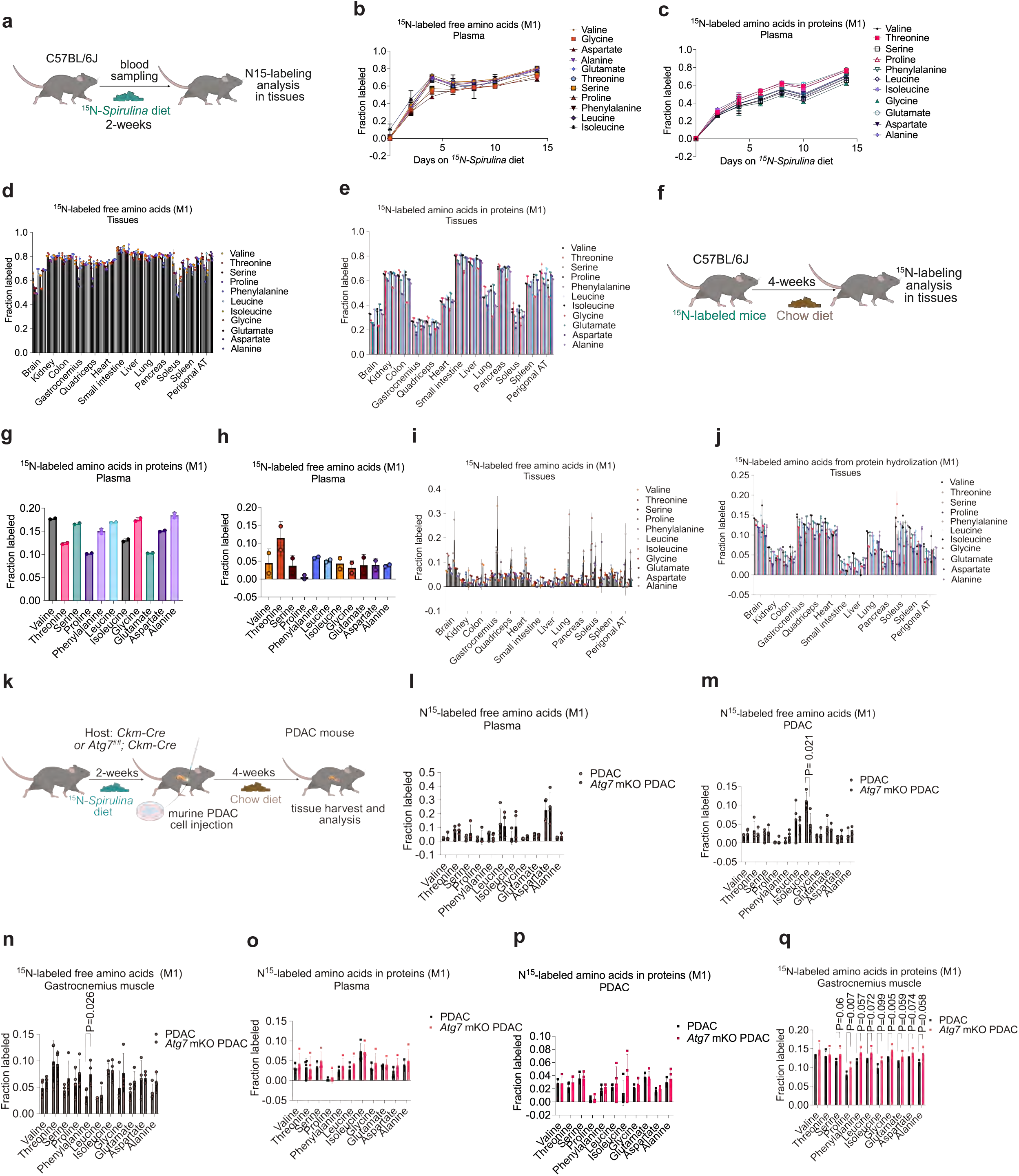
Muscle-derived nutrients support both tumor and host tissues. **a.** Schematic showing experimental design for tracing the fate of ^15^N-labeled dietary amino acids in tissues. **b-c.** % ^15^N-labeling of free amino acids in blood (b) and incorporated into plasma proteins (c) over time in 8-week-old mice fed an ^15^N-labeled *Spirulina* diet as in a (*n* = 4). **d-e.** % ^15^N-labeling of free amino acids in blood (d) and incorporated into protein in the specified tissues (e) after exposing 8-week-old mice to an ^15^N-labeled *Spirulina* diet for 2-weeks (*n* = 2). **f.** Schematic showing experimental design for assessing the fate of an ^15^N-labeled proteins from tissues by exposing mice which had been fed an ^15^N-labeled *Spirulina* diet for 2-weeks with a chow diet for 4-weeks. **g-j.** % ^15^N labeling of amino acids incorporated into plasma proteins (g) and free amino acids in blood (h), free amino acids in tissues (i), and amino acids incorporated into protein in the specified tissues (j) after 2-weeks of ^15^N-labeled *Spirulina* diet and 4-weeks of a chow diet as in f (*n* = 2). **k.** Schematic for use of ^15^N-*Spirulina* diet-based tissue labeling to assess amino acid fate in mice bearing orthotopic KPC PDAC tumors, comparing control animals and mice with muscle-specific loss of autophagy (*Atg7* mKO PDAC). **l-n.** % ^15^N-labeling of free amino acids in plasma (l), PDAC tumors (m), and gastrocnemius muscle (n) in mice bearing orthotopic PDAC tumors, comparing control animals and mice with muscle-specific loss of autophagy (*Atg7* mKO PDAC) exposed to the diets described in k (*n* = 4). **m-o.** % ^15^N-labeling of amino acids incorporated into proteins in plasma (m), PDAC tumors (n), and gastrocnemius muscle (o) in mice bearing orthotopic PDAC tumors, comparing control animals and mice with muscle-specific loss of autophagy (*Atg7* mKO PDAC) exposed to the diets described in k. (*n* = 4). 8-week-old male mice were used for these experiments. Statistical analysis was performed using unpaired two-sided *t*-tests, data are mean ± S.D and *n* represents the number of mice analyzed.

## Systemic amino acid availability impacts PDAC growth and survival

To gain further insight into how loss of muscle autophagy impairs PDAC growth, we performed bulk RNA sequencing analysis on KP^−/−^F PDAC tumors from mice with or without loss of *Atg7* in muscle. We observed that *Atg7* mKO KP^−/−^F PDAC tumors upregulate genes related to metabolism and cell cycle programs whereas they downregulate genes related to Kras signaling, focal adhesion, and extracellular matrix (ECM) remodeling (Extended Data Fig.18a-h). Immune genes were also altered in *Atg7* mKO KP^−/−^F PDAC tumors (Extended Data Fig.18b-e), suggesting that changes in immune cell composition may potentially contribute to slower tumor growth in *Atg7* mKO mice. Of note, this analysis revealed that tumors in *Atg7* mKO mice upregulate genes related to amino acid metabolism, glucose metabolism, and cholesterol synthesis (Fig. 6a-c and Extended Data Fig.18a), consistent with muscle wasting serving as a nutritional buffer in mice with PDAC-associated exocrine dysfunction leading to reduced systemic nutrient availability. Given the important role of muscle as an amino acid storage tissue for mammals, we hypothesized that dietary amino acid supplementation might restore tumor growth in *Atg7* mKO mice with KP^−/−^F PDAC. To test this possibility, we took advantage of diets formulated with intact protein, or with free amino acids to overcome impaired dietary amino acid absorption due to pancreatic exocrine dysfunction (Fig. 6d and Extended Data Fig.19a). While diets with different amounts of intact protein had no impact on either KP^−/−^F PDAC growth or tissue wasting (Extended Data Fig.19b-f), diets with increased free amino acids promoted KP^−/−^F PDAC tumor growth, without affecting body weight or adipose, liver, or muscle tissue mass (Fig. 6e-j). There was also a trend toward increased cell proliferation in KP^−/−^F PDAC tumors based on FAA content of the diet, while cell death was unchanged (Fig. 6k). Lastly, we questioned how different protein diets affected survival of *Atg7* mKO KP^−/−^F PDAC mice. Modifying intact dietary protein had no effect on survival of *Atg7* mKO KP^−/−^F PDAC mice (Fig. 6l), while reducing FAA in the diet conferred a small survival benefit and increasing FAA in the diet slightly decreased survival of *Atg7* mKO KP^−/−^F PDAC mice (Fig. 6m). Taken together, these data argue systemic amino acid availability, which is compromised from the diet via pancreatic exocrine dysfunction in early PDAC, is sustained by increased autophagic muscle breakdown, which impacts PDAC tumor growth and mouse survival.

**Figure 6:**
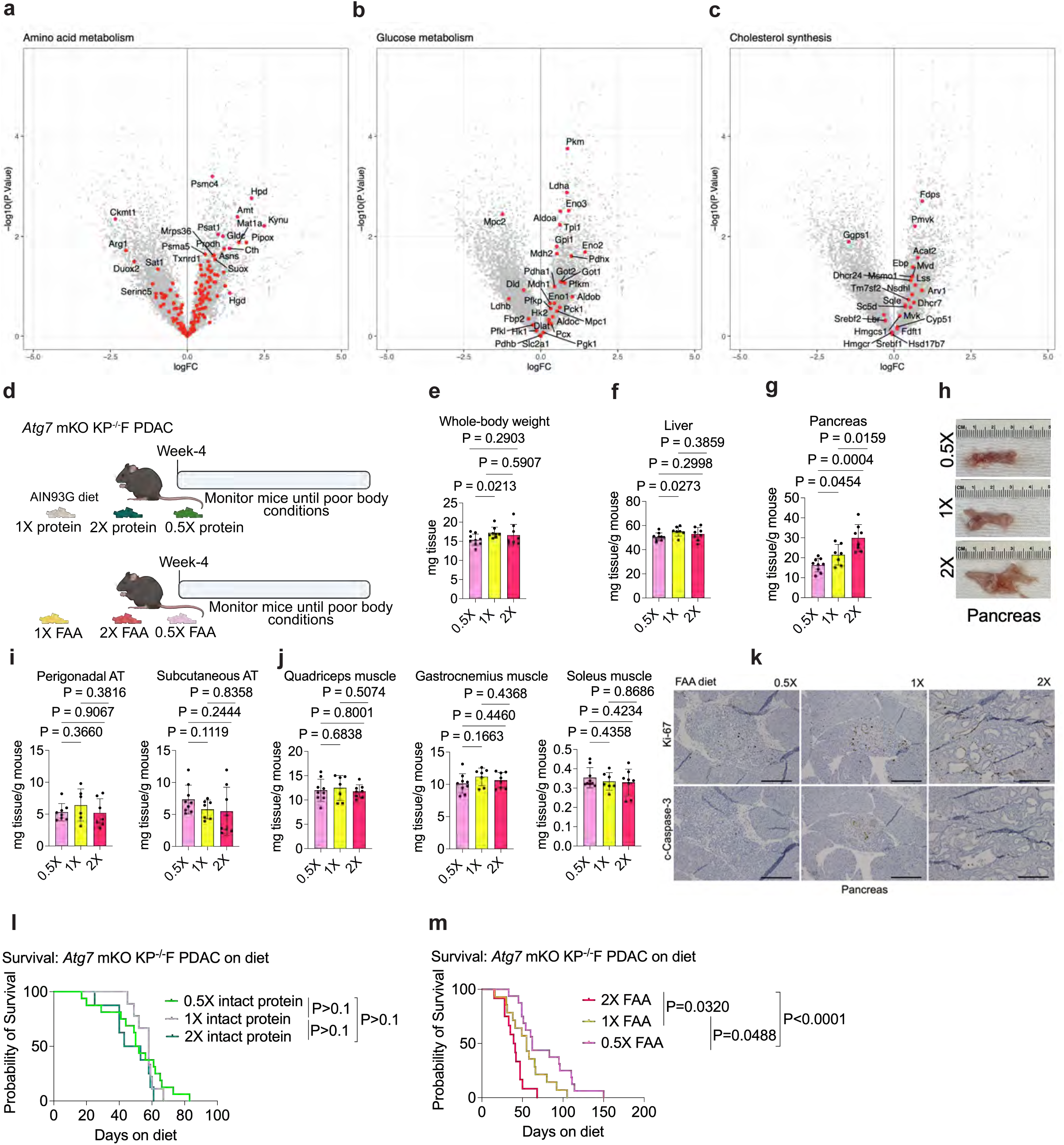
Amino acids promote PDAC tumor growth when muscle autophagy is impaired. **a-c.** Volcano plot of log2 fold change (logFC) in gene expression in PDAC tumors isolated from 6-week-old mice with KP^−/−^F PDAC (*Pdx-1-P2A-FlpO; Kras^FSF-G12D/+^; Trp53^Frt/Frt^*) without or with *Atg7* (*Pdx-1-P2A-FlpO; Kras^FSF-G12D/+^; Trp53^Frt/Frt^; Ckm-Cre; Atg7^fl/fl^,* (*Atg7* mKO KP^−/−^F PDAC)) loss in muscle for genes involved in amino acid (a), glucose (b), and cholesterol metabolism (c). (*n* = 5). **d.** Schematic showing experimental design for use of diets with different amounts of protein or different amounts of free amino acids instead of protein test effects on mice with KP^−/−^F PDAC and *Atg7* mKO KP^−/−^F PDAC. **e-g.** Whole-body weights (e), liver weights (f), and pancreatic tumor weights (g) normalized to total body weight measured at endpoint after 4-week-old mice were fed diets with different free amino acid (FAA) amounts as per the experiment outlined in d (*n* = 8 *Atg7* mKO KP^−/−^F PDAC on 0.5X protein, *n* = 7 *Atg7* mKO KP^−/−^F PDAC on 1X FAA, and *n* = 8 *Atg7* mKO KP^−/−^F PDAC on 2X FAA). **h.** Gross image of dissected pancreas following 2-weeks of the indicated FAA diet administration as outlined in d. **i-j.** Perigonadal and subcutaneous ATs (i), quadriceps, gastrocnemius, and soleus muscle (j) weights normalized to total body weight measured at endpoint after 4-week-old mice were fed diets with different FAA amounts as per the experiment outlined in d (*n* = 8 *Atg7* mKO KP^−/−^F PDAC on 0.5X FAA, *n* = 7 *Atg7* mKO KP^−/−^F PDAC on 1X FAA, and *n* = 8 *Atg7* mKO KP^−/−^F PDAC on 2X FAA). **k.** Ki-67 and cleaved Caspase-3 IHC to assess proliferation and apoptosis, respectively, in *Atg7* mKO KP^−/−^F PDAC tumor tissue after 2-weeks of the indicated FAA diet administration as outlined in d (*n* = 3). **l.** Survival of *Atg7* mKO KP^−/−^F PDAC mice fed diets with different amounts of intact protein as described in d (*n* = 16 *Atg7* mKO KP^−/−^F PDAC on 0.5X protein, *n* = 9 *Atg7* mKO KP^−/−^F PDAC on 1X protein, and *n* = 8 *Atg7* mKO KP^−/−^F PDAC on 2X protein diet). **m**. Survival of *Atg7* mKO KP^−/−^F PDAC mice fed diets with different amounts of FAAs as described in d (*n* = 16 *Atg7* mKO KP^−/−^F PDAC on 0.5X FAA, *n* = 14 *Atg7* mKO KP^−/−^F PDAC on 1X FAA, and *n* = 12 *Atg7* mKO KP^−/−^F PDAC on 2X FAA). Both male and female mice were used in these experiments. P-values on survival experiments in l-m were calculated with Gehan-Breslow-Wilcoxon test. Statistical analyses were performed using one-way ANOVA with Tukey’s post hoc test for data in other panels, data are mean ± S.D and *n* represents the number of mice analyzed. Scale bars: 200 μm.

## Discussion

Loss of muscle and adipose tissue mass contributes to morbidity and mortality in PDAC^10^, yet the mechanisms linking tumor progression to systemic tissue wasting remain incompletely understood. Here, we show that PDAC disrupts both pancreatic endocrine and exocrine function, leading to altered systemic metabolism and compromised nutrition that drives progressive loss of muscle and fat. Although restoring nutrition can attenuate tissue wasting, it may also promote tumor growth or unmask additional pathology, such as hyperglycemia^48,49,50,51^.

We find that skeletal muscle wasting has a greater impact on survival than adipose tissue loss. Consistently, postdiagnostic loss of skeletal muscle, but not adipose tissue, is associated with shorter survival in patients with advanced pancreatic cancer^10^. Mechanistically, we find muscle wasting in PDAC can be driven in part by reduced nutrition-induced activation of AMPK signaling, suppression of muscle mTORC1 activity, and consequent induction of autophagy. Activation of muscle proteolysis through Atrogin-1 and other Ub-E3 ligases^28,36^ also likely contributes to muscle wasting, but autophagy appears to be a dominant driver of muscle loss in the mouse PDAC models tested. Importantly, disrupting autophagy in the muscle can slow muscle wasting without restoring nutrition to the tumor, and thus may have a therapeutic role in PDAC if selective autophagy inhibition were possible in that organ.

The causes of cancer-associated cachexia, a syndrome that involves the profound loss of both adipose and lean body mass, is known to be multifactorial, with contributions from both increased energy expenditure and reduced food intake^2,3,33,92^. While it is unlikely that the early tissue wasting we observe due to pancreatic exocrine insufficiency in mice with PDAC fully explains all manifestations of cachexia associated with late-stage disease, disruption of normal pancreatic function by pancreatic tumors leading to changes in systemic metabolism with reduced nutrition appears is an important contributor to muscle and adipose tissue wasting, at least in the models tested. That dietary enzyme supplementation can improve nutrition and blunt the loss of muscle and fat tissue mass underscores a role for systemic nutrient availability in modulating peripheral tissue wasting.

In animals with PDAC, muscle autophagy emerges as a dynamic regulator of nutrient partitioning between host tissues and the tumor. Suppression of autophagy in muscle-specific *Atg7* knockout mice slowed tumor growth and protected host tissues without inducing global metabolic deficits. However, dietary supplementation with elemental amino acids in this setting increased tumor growth and decreased host survival, revealing that skeletal muscle normally competes with the tumor for amino acids. When this protective reservoir is bypassed, tumors gain unrestricted access to circulating nutrients, which appears to support tumor growth. Thus, tissue loss in PDAC can reflect a metabolic competition of different organs for metabolites in the setting of decreased nutrition.

The data support a model in which PDAC induces a starvation-like state by impairing dietary protein digestion, thereby activating skeletal muscle autophagy early in disease and establishing a nutritional buffer that shapes systemic metabolism and cancer progression. Consistent with this model, both systemic and muscle-specific *Atg7* loss slow implanted tumor growth and preserve tissue mass, but may involve distinct mechanisms^37,38^. In whole-body *Atg7*-deficient mice, tumor growth is restricted primarily by systemic arginine depletion, independent of effects on appetite^37^. Separately, autophagy suppresses Ccl2-driven anorexia as a general physiological function, thereby preserving feeding behavior and preventing lethal cachexia^93^. In contrast, muscle-specific autophagy impairment affects PDAC tumor growth by limiting a nutritional buffer. Thus, muscle autophagy provides direct nutrient support to tumors independent of other effects of autophagy loss on systemic metabolism or appetite caused by whole-body autophagy loss.

An unexpected finding from this study relates to changes in circulating glucose levels in PDAC-bearing mice, possibly informing why mouse PDAC models do not exhibit the diabetes phenotype found in some patients with pancreatic cancer^48,49,50,51,52,94^. Standard laboratory mouse diets contain predominantly complex carbohydrates, mainly starches that require enzymatic digestion to yield free glucose, whereas human diets are often much higher in simple sugars that rely less on pancreatic enzymes for absorption from the gut. In the setting of pancreatic exocrine dysfunction, this dietary distinction may substantially influence glucose availability. In patients, abundant dietary glucose may lead to diabetes due to pancreatic endocrine dysfunction, while also limiting fat wasting. This could explain why muscle wasting is more prominent in PDAC patients^8,9,10,51^. Notably, we observed that providing increased free glucose in the diet blunted fat tissue wasting in mice without affecting muscle wasting or PDAC growth. It also increased blood glucose, perhaps explaining some of the differences in systemic metabolism observed in mouse PDAC models and patients.

Our findings have translational implications for the management of PDAC-associated tissue wasting. First, although pancreatic enzyme replacement therapy (PERT) is routinely prescribed to treat exocrine insufficiency, well-designed epidemiological and interventional studies remain limited. Clinical data indicate that exocrine pancreatic insufficiency is prevalent in PDAC, particularly in patients with pancreatic head tumors^17^. Optimizing PERT may alleviate the reduced nutritional state induced by impaired protein digestion, thereby reducing reliance on muscle autophagy and helping preserve host tissue mass. Second, although dietary supplementation with high protein, leucine, and fish oil improves muscle function and daily activity in cancer cachexia models^95^, our results indicate that free amino acid (FAA) supplementation may provide more effective nutritional support than intact protein in the setting of pancreatic exocrine dysfunction. However, our findings also caution that increasing systemic amino acid availability can also support tumor growth, underscoring the need for considering both effects on both muscle mass and tumor growth when evaluating interventions to limit tissue wasting. Finally, the observation that early PDAC can cause disruption of normal pancreatic function might be useful in finding pancreatic cancers earlier, which could also improve patient outcomes.

## Material and methods

### Animal studies

All animal experiments performed in this study were approved by the MIT Institutional Animal Care and Use Committee (IACUC, recent protocol 2409000733; previous protocols 012200425 and 011900122). Male and female C57BL/6J mice between 1-and 4-months old were used in this study. All animals were housed at ambient temperature and humidity (18-23 °C, 40-60% humidity) with a 12h light and 12h dark cycle and co-housed with littermates with ad libitum access to water, unless otherwise stated. All experimental groups were age-matched, numbered and assigned based on treatment, and experiments were conducted in a blinded manner. Data were collected from distinct animals, where *n* represents biologically independent samples. Statistical methods were not performed to predetermine sample size.

### Mouse strains

The following mouse strains were studied in here: C57BL/6J mouse (The Jackson Laboratory, 000664), Nu/J mouse (The Jackson Laboratory, 002019), *Ckm-Cre* (The Jackson Laboratory, 006475), the conditional LysoTag mouse commonly known as Conditional LysoTag (The Jackson Laboratory, 035401), the conditional *Atrogin-1^fl/fl^*(Taconic, 9569), conditional *Atg7^fl/fl^* mouse (provided by Dr. Eileen White from Rutgers University, New Brunswick, NJ-USA). Genetically engineered mouse models for colorectal cancer included *Villin-Cre-ERT^2^; APC^fl/fl^; Kras^LSL-G12D/+;^ Trp53^fl/fl^* (provided by Dr. Ömer H. Yilmaz, Massachusetts Institute of Technology, Cambridge, MA-USA). Pancreatic cancer models included *Pdx1-P2A-FlpO* mice (generated and provided by Dr. Tyler Jacks at Massachusetts Institute of Technology, Cambridge, MA-USA), conditional *Kras^Frt-stop-Frt-G12D^*mouse (The Jackson Laboratory, 023590), conditional *Tp53^Frt/Frt^* mouse (The Jackson Laboratory, 017767), conditional *Kras^LSL-G12D/+^* mouse (The Jackson Laboratory, 008179), conditional *Trp53^LSL-R172H/+^* mouse (The Jackson Laboratory, 008652), conditional *Trp53^fl/fl^* mouse (The Jackson Laboratory, 008462), and *Pdx1-Cre* mouse (The Jackson Laboratory, 014647). All genetically engineered mice were backcrossed for greater than 6 generations and then maintained on a C57BL/6J background.

### Generation of *Pdx1-P2A-FlpO* knock-in mice

DNA mixes (1:1 mix of ‘U6-sgPdx1-eCas9-T2A-BlastR’ + ‘*Pdx1-P2A-FlpO* TV’; [sgPdx1 5’-GACAGCAGTCTGAGGGTGAG-3’]) were ethanol precipitated prior to DNA (1 µg) transfection of approximately 3 x 10^5^ KP17 mESCs in a gelatin-coated 24-well plate using Lipofectamine 2000 according to manufacturer instructions. mESCs were selected with Blasticidin (6 µg/mL) for 2 days, starting approximately 36 hours post-transfection, prior to low-density re-plating on MEF feeder lines. Large mESC colonies were manually picked using a stereomicroscope, expanded and evaluated for correct integration using PCR spanning both the 5’ and 3’ homology arms. Correct clones by PCR evaluation were evaluated using Southern blot analysis. Briefly, genomic DNA was digested overnight with Kpn1, digestions were electrophoresed on 0.7% agarose gels and blotted to Amersham Hybond XL nylon membranes (GE Healthcare). Samples were probed with 32P-labeled 5’ external, 3’ external, and internal probes applied in Church buffer^96^. Correctly targeted clones were injected into albino *C57BL/6J* blastocysts. Chimerism was assessed by coat color. Germline transmission was confirmed by PCR, after which *Pdx1-P2A-FlpO* mice were backcrossed onto a C57BL/6J background and crossed sequentially with *Atg7^fl/fl^*; *Ckm-Cre* and *Kras^FSF-G12D/+;^ Trp53^Frt/Frt^*mice, as described in the text.

### Whole-body and tissue weights

For assessment of tissue weights, mice were weighed immediately prior to euthanasia. The quadriceps, gastrocnemius, and soleus muscles from the left hindlimb, perigonadal adipose tissue (left side), subcutaneous adipose tissue (left inguinal), liver, and pancreas were then dissected and weighed. Tissue weights from each mouse were normalized to the corresponding whole-body weight.

### Tumor cell isolation and cell culture

Cells were isolated from primary mouse pancreatic tumors from KP^−/−^C and KPC as described previously^62,97^. RIL-175 mouse HCC cells were isolated by the Duda Laboratory from hepatic tumors arising in C57BL/6 mice as previously described^62^. LUAD cells were obtained from Kras*^LSL-^*^G12D/+^; *Trp53^fl/fl^; Ad-Cre* lung cancer mice as previously described^62^. Cells were cultured in DMEM (Corning, 10-013-CV) supplemented with 10% heat-inactivated FBS. Penicillin-streptomycin was added only at the time of cell isolation from mice. Cells were regularly tested for mycoplasma using the MycoAlert Plus kit (Lonza). All cell lines previously were verified using an antibody to detect the p. Gly12Asp *Kras* mutation (Cell Signaling, 14429), was also verified by Sanger Sequencing and, if applicable, by fluorescent TdTomato expression^62^.

## Tumor cell implantation

### Subcutaneous injection of mouse PDAC cells

C57BL/6J mice were injected with 1 x 10^5^ mouse PDAC cells as previously described^11,62,97,99^. Cells were injected subcutaneously into one flank in 100 µL of PBS per injection.

### Tumor transplantation with mouse PDAC cells

For all organ injection methods, animals were anesthetized using isoflurane, the appropriate surgical area was depilated and disinfected with alternating Betadine/isopropyl alcohol. All mice received pre-operative analgesia with extended-release Buprenorphine and were followed post-operatively for any signs of discomfort or distress. Mice aged approximately 6-8 weeks (for C57B16/J) or 8-10 weeks (for Nu/J) were injected with 1 x 10^5^ mouse PDAC cells SS3371 derived from primary tumors arising in C57BL/6J KPC mice or AL1376 PDAC cells, isolated from C57BL/6J KP^−/−^C mice. Cells were delivered into the pancreas as previously published^11,62,97^.

### Intrahepatic injection of mouse PDAC cells

To introduce cells into the liver, after anaesthetization, a small incision was introduced to exteriorize one lobe of the liver, and 1 x 10^5^ mouse PDAC cells were directly injected into the liver as described previously^62^. Mice were monitored for recovery and appropriate postoperative care was provided.

Mice were euthanized 4-weeks after tumor cell injection or at signs of distress. All mice within the same experimental group were euthanized at the same time point.

### Transplantation of colon cancer organoids via multiple routes

1 x 10^5^ of *Villin-Cre-ERT^2^; APC^f//fl^; Kras^LSL-G12D/+;^ Trp53^fl/fl^* colon cancer organoids were implanted using three distinct approaches^63,90^. Syngeneic C57BL/6J mice (age-12-weeks) were used for all transplantation procedures.

Mice were euthanized 6-weeks after tumor cell injection or at signs of distress. All mice within the same experimental group were euthanized at the same time point.

### Splenic Injection with hepatic engraftment of mouse colon cancer organoids

For hepatic tumor modeling via splenic injection, a small (∼1 cm) skin incision was made in the left subcostal area, and the spleen was visualized through the peritoneum. A small incision (∼1 cm) was made through the peritoneum overlying the spleen, and the spleen was exteriorized. A 30-gauge needle was inserted into the splenic parenchyma and 100 μL (containing 1 x 10^5^ *Villin-Cre-ERT^2^; APC^fl/fl^; Kras^LSL-G12D/+;^ Trp53^fl/fl^* colon cancer organoids in PBS) was slowly injected into the spleen to allow portal vein dissemination and subsequent hepatic engraftment. Following injection, hemostasis was confirmed and a splenectomy was performed by ligating the splenic vessels with 5-0 vicryl sutures and removing the spleen. The peritoneal and skin layers were sutured independently using 5-0 vicryl sutures.

Mice were euthanized 6-weeks after tumor cell injection or at signs of distress. All mice within the same experimental group were euthanized at the same time point.

### Direct pancreatic injection of mouse colon cancer organoids

For pancreatic tumor modeling, a small (∼1 cm) skin incision was made in the left subcostal area, and the spleen was visualized through the peritoneum. A small incision (∼1 cm) was made through the peritoneum overlying the spleen, and the spleen and pancreas were exteriorized. A 30-gauge needle was inserted into the pancreatic parenchyma parallel to the main pancreatic artery and 50 μL (containing 1 x 10^5^ *Villin-Cre-ERT^2^; APC^fl/fl^; Kras^LSL-G12D/+;^ Trp53^fl/fl^* colon cancer organoids in PBS) was injected into the pancreatic parenchyma. Successful injection was visualized by formation of a fluid-filled region within the pancreatic parenchyma without leakage. The pancreas and spleen were gently internalized, and the peritoneal and skin layers were sutured independently using 5-0 vicryl sutures.

Mice were euthanized 6-weeks after tumor cell injection or at signs of distress. All mice within the same experimental group were euthanized at the same time point.

### Rectal submucosal injection of mouse colon cancer organoids

For rectal/colon tumor modeling, *Villin-Cre-ERT^2^; APC^fl/fl^; Kras^LSL-G12D/+;^ Trp53^fl/fl^* mice were positioned in dorsal recumbency, and the perianal area was depilated and disinfected. Forceps were gently inserted into the rectum to visualize the rectal mucosa. Using a 30-gauge needle attached to a Hamilton syringe, 50 µL of 4-hydroxytamoxifen (100 µM in PBS) was injected into the submucosal layer of the rectal wall. Successful submucosal injection was confirmed by the appearance of a localized, fluid-filled bleb beneath the intact mucosa. The needle was slowly withdrawn to minimize backflow, and the anoscope was carefully removed. No skin incision was required for this procedure. For all injection procedures, organoid cell suspensions, when used, were prepared fresh immediately prior to transplantation and kept on ice throughout the procedure.

Mice were euthanized 6-weeks after tumor cell injection or at signs of distress. All mice within the same experimental group were euthanized at the same time point.

## Diet studies

### Protein and free amino acid (FAA) diets

Protein diets containing 0.5X protein (TD.210660; Inotiv), 1X protein (TD.06706; Inotiv), or 2X protein (TD.210586; Inotiv) were generated from a modified AIN-93G purified diet using vitamin-free test (VFT) casein as the protein source and were purchased from Inotiv. The 0.5X protein diet was formulated by reducing the casein content of TD.06706 by 50%, whereas the 2X protein diet was formulated by doubling the amounts of casein and cysteine relative to TD.06706. Free amino acid (FAA) diets containing 0.5X FAA (TD.210661; Inotiv), 1X FAA (TD.170213; Inotiv), or 2X FAA (TD.210587; Inotiv) were generated from a modified AIN-93G diet (TD.06706; Inotiv). The 1X FAA diet consisted of individual free amino acids replacing intact protein, with an amino acid composition matched to VFT casein. The 0.5X FAA diet contained half the amount of each amino acid relative to the 1X FAA diet, whereas the 2X FAA diet contained double the amount of each amino acid. To vary protein or free amino acid content across diets, carbohydrate levels were adjusted; accordingly, fat content was maintained constant across all diet formulations.

Four-week-old PDAC-bearing KP^−/−^C mice or *Atg7* mKO KP^−/−^F mice or non-tumor-bearing control mice were placed on the indicated protein or FAA diets. For short-term feeding experiments, mice were maintained on the diets for 2-weeks. At the end of the feeding period, blood was collected by cardiac puncture prior to tissue harvest. Mice were euthanized, and tissues were collected in 10% neutral buffered formalin for histological analysis.

For long-term feeding experiments, mice were maintained on the indicated diets until humane endpoints were reached. Blood was collected by cardiac puncture prior to tissue harvest. Mice were euthanized, and tissues were collected in 10% neutral buffered formalin for histological analysis.

### High-starch diet

AIN-93G powdered purified diet (TD.94045; Inotiv) was mixed with either waxy cornstarch or resistant starch to generate high-starch diets. Briefly, 50 g of AIN-93G diet was combined with 50 g waxy corn starch lacking resistant starch (AMIOCA; Ingredion, Bridgewater, NJ) or with resistant starch derived from high-amylose corn starch (Hi-Maize 260; Ingredion, Bridgewater, NJ). The powder mixtures were formed into a dough using sterile tap water, portioned into 5 g pellets, and dried overnight at room temperature. Dried pellets were stored at 4 °C, and one 5 g pellet per mouse was provided once every 24 h.

Based on starch composition, the AIN-93G high-starch diet (AIN-93G HSD) contained approximately 75% digestible starch. Because the resistant starch preparation consisted of approximately 40% digestible starch and 60% resistant starch, the AIN-93G control starch diet (AIN-93G CSD) contained approximately 45% digestible starch.

Six-week-old PDAC-bearing KP^−/−^C mice or non-tumor-bearing control mice were placed on either the AIN-93G HSD or AIN-93G CSD for 24 h following an overnight fast. Mice were randomly assigned to diet groups. Blood glucose levels were measured in the morning by tail vein puncture using a Contour glucose meter (Ascensia Diabetes Care).

### [U-^13^C]-starch oral gavage

Uniformly ^13^C-labeled starch ([U-^13^C]-starch from algae, 99 atom% ^13^C; Sigma-Aldrich, 605336) was dissolved at 80 mg mL⁻^1^ in blood bank saline and heated to 90 °C for 15 min to gelatinize the starch. Preparations were made fresh on the day of use and maintained under constant stirring until administration. Mice were food-restricted overnight prior to gavage. Each mouse received [U-^13^C]-starch at a dose of 0.01 mL g⁻^1^ body weight by oral gavage. Blood samples were collected immediately in EDTA-containing tubes (Sarstedt, DE) before gavage and at 30, 60, and 90 min, and 2, 4, and 6 h post-gavage. Plasma was isolated, and [U-^13^C]-glucose enrichment was quantified from 5 μL of plasma by GC-MS as described above in this study.

### [U-^13^C]-glucose oral gavage

Uniformly ^13^C-labeled glucose ([U-^13^C]-glucose; Cambridge Isotope Laboratories) was dissolved in sterile blood bank saline (Azer Scientific) and administered at a dose of 2 g kg⁻^1^ whole-body weight, as previously described⁹⁵. Six-week-old KP^−/−^C or non-tumor bearing mice were food-restricted overnight prior to gavage. Each mouse received 0.01 mL g⁻^1^ body weight of the [U-^13^C]-glucose solution by oral gavage. Blood samples were collected immediately in EDTA-containing tubes (Sarstedt, DE) before gavage and at 30, 60, and 90 min, and 2, 4, and 6 h post-gavage. Plasma was isolated, and [U-^13^C]-glucose enrichment was quantified from 5 μL of plasma by GC-MS as described in this study.

### ^15^N-labeling of *Schizosaccharomyces pombe* and proteome extraction

*S. pombe* was cultured at 30 °C and 220 rpm in Edinburgh minimal medium (EMM2) as previously described^100^, using ^15^NH_4_Cl (Sigma-Aldrich, 299251) as the sole nitrogen source. At an OD_600_ of 0.8-1.0, cultures were centrifuged at 3,000 x g for 5 min at 4 °C. Pellets were washed with 100 mL ice-cold water and centrifuged again under the same conditions. Pellets were resuspended in lysis buffer prepared as described previously^100^, supplemented with protease inhibitor (Roche, 11836170001) according to the manufacturer’s instructions. Cells underwent eight freeze-thaw cycles, followed by sonication in a 4 °C water bath for eight cycles of 5 min each, with 5 min pauses on ice between cycles. Lysates were centrifuged at 11,900 x g for 30 min at 4 °C, and the supernatants were collected for further processing.

Overnight dialysis was performed at 4 °C in PBS using 3.5 kDa molecular weight cutoff dialysis flasks (Thermo Fisher Scientific, 87761). To remove free amino acids, the proteome solution was concentrated using 3 kDa cutoff filter tubes (Sigma-Aldrich, UFC900324) with multiple rounds of centrifugation at 3,000 x g for 1 h at 4 °C. Protein concentration in the PBS solution was determined using the Pierce BCA Protein Assay Kit (Thermo Fisher Scientific, 23225).

### Oral gavage of ^15^N-labeled proteome of *Saccharomyces pombe*

Six-week-old PDAC-bearing KP^−/−^C mice or non-tumor-bearing control mice were fasted overnight. The following morning, mice received ^15^N-labeled yeast proteome by oral gavage at a dose of 1 g protein per kg whole-body weight using a plastic feeding tube (MDAFN2425S; Pet Surgical).

Blood samples were collected into EDTA-coated tubes (Sarstedt) immediately prior to gavage and at 30, 60, and 90 min, and 2, 4, and 6 h post-gavage. Plasma was isolated by centrifugation, and ^15^N-labeled free amino acid enrichments were quantified from 5 µL of plasma by GC-MS as described in this study.

### ^15^N-Spirulina gavage

^15^N-*Spirulina* diet powder (5 g; MF-SPIRULINA-N; Cambridge Isotope Laboratories) was resuspended in 20 mL PBS. To improve solubility, the suspension was vortexed at 4 °C for 15 min, sonicated for 5 min, and centrifuged at maximum speed for 20 min at 4 °C. The supernatant was transferred to a clean tube and used for gavage.

Six-week-old PDAC-bearing KP^−/−^C mice or non-tumor-bearing control mice were fasted overnight. The following morning, mice received the ^15^N-*Spirulina* solution by oral gavage at a dose of 10 µL per g body weight using a plastic feeding tube (MDAFN2425S; Pet Surgical).

Blood samples were collected into EDTA-coated tubes (Sarstedt) immediately prior to gavage and at 30, 60, and 90 min, and 2, 4, and 6 h post-gavage. Plasma was isolated by centrifugation as described in this study, and ^15^N-labeled free amino acid enrichment was quantified from 5 µL of plasma by GC-MS as described in this study.

### Dietary amino acid restriction

To assess the impact of systemic amino acid availability on whole-body physiology and nutrient deprivation-associated tissue wasting, male and female C57BL/6J mice (12 weeks old) were placed on Teklad amino acid-modified purified diets (Inotiv) for 4 weeks. Diets included a nitrogen-matched control diet (TD.01084), tryptophan-depleted (-Trp; TD.130674), methionine-depleted (-Met; TD.140119), lysine-depleted (-Lys; TD.210440), branched-chain amino acid-depleted (-BCAA; TD.210441), or serine and glycine-deficient (-Ser and Gly; TD.160752) diets.

Mice were randomly assigned to diet groups. All plasma and tissue collections were performed at the same time of day to control for circadian variation. Prior to tissue harvest, blood was collected by facial vein puncture into EDTA-coated tubes (Sarstedt, 41.1395.105) and processed to obtain plasma by centrifugation at 800 x g for 10 min at 4 °C. Plasma samples were flash-frozen and stored at –80 °C until analysis. Tissues were rinsed in cold saline, blotted dry, and fixed in 10% neutral buffered formalin for histological analysis.

### Pancreatic enzyme supplemented (PES) diet

AIN-93G powdered purified diet (TD.94045; Inotiv) was mixed with a pharmaceutical-grade pancreatic enzyme preparation (PancreVed powder; VEDCO). Briefly, 100 g of AIN-93G diet was combined with 0.8 g PancreVed powder and 0.5 g cherry flavor powder (755834; Carmi Flavors). The powder mixture was formed into a dough using sterile tap water, portioned into 5 g pellets, and dried overnight at room temperature. Dried pellets were stored at 4 °C, and one 5 g pellet per mouse was provided once every 24 h.

### PES diet studies in PDAC and control mice

Male and female C57BL/6J mice (12 weeks old) were orthotopically implanted with 1 x 10^5^ mouse PDAC cells suspended in 100 µL PBS into the tail of the pancreas. Following surgery, mice were maintained on standard ad libitum diet for a 3-day post-operative recovery period. Mice were then blindly randomized into two groups and fed either the pancreatic enzyme-supplemented AIN-93G diet or control AIN-93G diet containing 0.5% cherry flavor for 2-weeks. At the end of the feeding period, mice were euthanized and tissues were collected in 10% neutral buffered formalin for histological analysis.

Male and female non-tumor bearing C57BL/6J mice were fed either the pancreatic enzyme-supplemented AIN-93G diet or control AIN-93G diet containing 0.5% cherry flavor for 4-weeks. At the end of the feeding period, mice were euthanized and tissues were collected in 10% neutral buffered formalin for histological analysis.

### Oral gavage of pancreatic enzymes

4-week-old male and female KP^−/−^C mice were randomly assigned to two groups: PBS vehicle control or pancreatic enzyme solution (PES). Control mice received PBS by oral gavage at a dose of 10 µL/g body weight per day for 2 weeks. The PES group received a 2% pancreatic enzyme solution (w/v; g PancreVed/mL PBS), a pharmaceutical enzyme mixture containing porcine pancreatic amylase, lipase, and proteases (PancreVed powder; VEDCO), administered by oral gavage at 10 µL/g body weight per day for 2 weeks.

### 20% (w/v) glucose (D20) drinking water

As previously reported^101^, A 20% (w/v) glucose solution (D20) was prepared by dissolving D-(+)-glucose (Sigma-Aldrich, G8270) in drinking water and sterilized through a 0.22 µm filter (Corning, CLS430517). Mice were provided ad libitum access to D20 drinking water for two weeks. Whole-body weights and blood glucose levels were monitored throughout the treatment period. At the endpoint, mice were euthanized by cervical dislocation, and tissues were collected and weighed.

For combination experiments, mice received ad libitum access to D20 drinking water together with either 1X Protein or 2X Protein diets for two weeks.

For survival experiments, mice were provided ad libitum access to D20 drinking water and monitored until humane endpoints were reached. Mice were euthanized at the humane endpoint, and tissues were collected for weight analysis.

### ^15^N-Spirulina diet and ^15^N tissue labeling

Male C57BL/6J mice (12 weeks old) were assigned to a ^15^N-*Spirulina* diet (MF-SPIRULINA-N; Cambridge Isotope Laboratories) for 2 weeks (*n* = 4). Body weight and food intake were recorded weekly. Plasma was collected in the morning at baseline and every other day thereafter via the lateral tail vein. At the end of the 2-week feeding period, two mice were euthanized by cervical dislocation. Serum was collected by cardiac puncture prior to tissue harvest. All tissues were snap-frozen in liquid nitrogen, and serum and tissues were stored at –80 °C until analysis.

The remaining two mice were maintained on an *ad libitum* diet for an additional 4 weeks. Body weight and food intake were recorded weekly, and plasma was collected every other day via the lateral tail vein in the morning. At the end of the feeding period, mice were euthanized by cervical dislocation. Serum was collected by cardiac puncture prior to tissue harvest. All tissues were snap-frozen in liquid nitrogen and stored at –80 °C until analysis.

### ^15^N tissue labeling of PDAC-bearing mice

Male *Ckm-Cre* or *Atg7* muscle-specific knockout (*Atg7* mKO) C57BL/6J mice (12 weeks old) were assigned to a ^15^N-*Spirulina* diet (MF-SPIRULINA-N; Cambridge Isotope Laboratories) for 2-weeks. Whole-body weight and food intake were recorded weekly, and plasma was collected at baseline and every other day via the lateral tail vein in the morning. At the end of the 2-week feeding period, 1 x 10^5^ mouse PDAC cells suspended in 100 µL PBS were orthotopically implanted into the tail of the pancreas. Mice were then maintained on an ad libitum diet for an additional 4-weeks. During this period, body weight and food intake were recorded weekly, and plasma was collected every other day in the morning. At the end of the feeding period, mice were euthanized by cervical dislocation. Serum was collected by cardiac puncture prior to tissue harvest, including the tumor-bearing pancreatic tail and non-tumor-bearing pancreatic head. All tissues were snap-frozen in liquid nitrogen, and serum and tissues were stored at –80 °C until analysis.

### Isolation of muscle tissue interstitial fluid and plasma

Interstitial fluid was collected from mouse tissues using an adapted protocol based on prior published work^45,46^. Six-week-old male KP^−/−^C mice maintained on an ad libitum diet were used for muscle interstitial fluid isolation. All mice were euthanized at the same time of day to control for circadian effects on metabolism and were euthanized by cervical dislocation to minimize metabolic artifacts associated with CO_2_ exposure. Immediately following euthanasia, muscle tissues were maintained on a chilled metal block on ice throughout harvest. Tissues were briefly rinsed in ice-cold saline (Azer Scientific, 16005), blotted dry on filter paper (VWR, 28298-020), and placed into 50 mL conical tubes lined with a 20 µm nylon mesh filter (Spectrum Labs, 148134). EDTA (0.25 µL of 0.5 M, pH 8.0) was added to the bottom of each tube to prevent coagulation. Samples were centrifuged at 450 x g for 10 min at 4 °C, and the flow-through was collected and centrifuged again at 450 x g for 10 min at 4 °C. The resulting interstitial fluid was flash-frozen and stored at –80 °C until analysis. Prior to tissue harvest, plasma was collected from live mice by facial vein puncture into EDTA-coated tubes (Sarstedt, 41.1395.105) and centrifuged at 800 x g for 10 min at 4 °C. The plasma supernatant was flash-frozen and stored at –80 °C until analysis.

## Mass spectrometry metabolite measurements

### Quantification of metabolite levels in biological fluids

Metabolite quantification in murine fluid samples was performed as described previously^45,46^. In brief, 5 μL of sample or external chemical standard pool (ranging from ∼5 mM to ∼1 μM) was mixed with 45 μL of acetonitrile: methanol: formic acid (75:25:0.1) extraction mix including isotopically labeled internal standards. All solvents used in the extraction mix were HPLC grade. Samples were vortexed for 15 min at 4 °C and insoluble material was sedimented by centrifugation at 16,000 x g for 10 min at 4 °C. 20 µL of the soluble polar metabolite extract was taken for Liquid-chromatography-Mass Spectrophotometry (LC-MS) analysis. Following analysis by LC-MS, metabolite identification was performed with XCalibur 2.2 software (Thermo Fisher Scientific) using a 5-ppm mass accuracy and a 0.5 min retention time window. For metabolite identification, external standard pools were used for assignment of metabolites to peaks at given m/z and retention time. Absolute metabolite concentrations were determined as published.

### LC-MS analysis

As previously described^45,46^, metabolite profiling was conducted on a QExactive benchtop orbitrap mass spectrometer equipped with an Ion Max source and a HESI II probe, which was coupled to a Dionex UltiMate 3000 HPLC system (Thermo Fisher Scientific). External mass calibration was performed using the standard calibration mixture every 7 days. An additional custom mass calibration was performed weekly alongside standard mass calibrations to calibrate the lower end of the spectrum (m/z 70-1050 positive mode and m/z 60-900 negative mode) using the standard calibration mixtures spiked with glycine (positive mode) and aspartate (negative mode). 2 μL of each sample was injected onto a SeQuant® ZIC®-pHILIC 150 x 2.1 mm analytical column equipped with a 2.1 x 20 mm guard column (both 5 mm particle size; EMD Millipore). Buffer A was 20 mM ammonium carbonate, 0.1% ammonium hydroxide; Buffer B was acetonitrile. The column oven and autosampler tray were held at 25 °C and 4 °C, respectively. The chromatographic gradient was run at a flow rate of 0.150 mL min^−1^ as follows: 0-20 min: linear gradient from 80-20% B; 20-20.5 min: linear gradient form 20-80% B; 20.5-28 min: hold at 80% B. The mass spectrometer was operated in full-scan, polarity-switching mode, with the spray voltage set to 3.0 kV, the heated capillary held at 275 °C, and the HESI probe held at 350 °C. The sheath gas flow was set to 40 units, the auxiliary gas flow was set to 15 units, and the sweep gas flow was set to 1 unit. MS data acquisition was performed in a range of m/z = 70-1000, with the resolution set at 70,000, the AGC target at 1 x 10^6^, and the maximum injection time at 20 msec.

### Stable isotope-labeled nutrient infusion experiments in mice

[U-^13^C]-glucose (Cambridge Isotope Laboratories) infusion into control or tumor-bearing mice was performed as previously described. Five-week-old mice underwent surgical implantation of a jugular vein catheter 3-4 days prior to tracer infusion. Mice were food-restricted for 6 h before infusion and remained conscious and freely moving throughout the experiment. [U-^13^C]-glucose was infused at a constant rate of 0.4 mg min⁻^1^ for 6 h, and blood samples were collected hourly to assess plasma ^13^C-glucose enrichment. At the end of the infusion, mice were euthanized and plasma and tissues were harvested within 5 min and flash-frozen in liquid nitrogen for GC-MS analysis. All isotope-labeling experiments were performed at the same time of day to minimize circadian variability.

### Plasma metabolite extraction for GC-MS-based analysis

Blood collected from animals was immediately placed in EDTA tubes (Sarstedt, DE) and centrifuged to separate plasma. For absolute quantification of polar metabolites in biofluids, plasma or TIF was mixed 1:1 with a solution of isotopically labelled amino acids, pyruvate, lactate and β-hydroxybutyrate at known concentrations (Cambridge Isotope Laboratories, MSK-A2-1.2, CNLM-3819-H, CLM-1822-H, CLM-2440, CLM-10768 and CLM-3853) in glass vials.

Metabolites were extracted in 1.5 mL of dichloromethane: methanol (containing 25 mg L^−1^ of butylated hydroxytoluene (Millipore Sigma, B1378): 0.88% KCl (w/v) (8:4:3 v/v/v), vortexed for 15 min and centrifuged at maximum speed for 10 min. The extraction buffer contained 0.75 µg ml^−1^ of norvaline. Polar metabolites (aqueous fraction) were transferred to Eppendorf tubes, dried under nitrogen gas and stored at –80 °C until further analysis.

### Glucose analysis in the plasma by Gas chromatography-mass spectrometry (GC-MS)

As described previously^98^, dried and frozen metabolite extracts were derivatized with 50 µl of 2% (w/v) hydroxylamine hydrochloride in pyridine (Millipore Sigma) for 60 min at 90 °C, followed by derivatization with 100 µL of propionic anhydride (Millipore Sigma) for 30 min at 60 °C. Derivatized samples were then dried under nitrogen gas and resuspended in 100 µL of ethyl acetate (Millipore Sigma) in glass GC-MS vials. Samples were analyzed by GC-MS as described above, except helium was used as the carrier gas at a constant flow rate of 1.1 mL min^−1^, and 1 µL of sample was injected in spitless mode at 250 °C. Following the injection, the GC oven was held at 80 °C for 1 min, ramped to 280 °C at 20 °C per min and held at 280 °C for 4 min.

### GC-MS analysis of polar metabolites

Polar metabolites were analyzed by gas chromatography-mass spectrometry (GC-MS) as described previously^74^. Dried and frozen metabolite extracts were derivatized with 16 µL of MOX reagent (Thermo Fisher Scientific, TS-45950) for 60 min at 37 °C, followed by derivatization with 20 µL of *N*-tert-butyldimethylsilyl-*N*-methyltrifluoroacetamide with 1% tert-butyldimethylchlorosilane (Millipore Sigma, 375934) for 60 min at 60 °C. Derivatized samples were analyzed by GC-MS, using a DB-35MS column (Agilent Technologies, 122-3832) installed in an Agilent 7890B gas chromatograph coupled to an Agilent 5997B mass spectrometer. Helium was used as the carrier gas at a constant flow rate of 1.2 ml min^−1^. A microliter of sample was injected in split mode (1:10) at 270 °C. Following the injection, the GC oven was held at 100 °C for 1 min, increased to 105 °C at 2.5 °C per min, held at 105 °C for 2 min, increased to 250 °C at 3.5 °C per min and then ramped to 320 °C at 20 °C per min. The MS system operated under electron impact ionization at 70 eV, and the MS source and quadrupole were held at 230 °C and 150 °C, respectively. The detector was used in scanning mode with an ion range of 100-650 *m*/*z*. Total ion counts were determined by integrating appropriate ion fragments for each metabolite^102^ using EL-MAVEN software (Elucidata). Mass isotopologue distributions were corrected for natural abundance using IsoCorrectoR^103^. Metabolite data were normalized to the internal standard norvaline and tissue weights. Absolute concentrations of metabolites were calculated based on the known concentrations of isotopically labelled internal standards.

### Muscle Lyso-IP

To optimize muscle Lyso-IP, muscle LysoTag (*Ckm-Cre*; *TMEM192-3xHA)* mice were fasted or fed overnight. Following euthanasia, quadriceps muscles were immediately dissected on an ice-cold plastic dish. Approximately 200 mg of muscle tissue per sample was collected to reduce variability and minimize processing time. Samples were processed immediately without freezing, following the original procedure described previously^85^.

Muscle biopsies were gently homogenized in 1 mL ice-cold PBS containing protease (Roche, 11836170001) and phosphatase inhibitor cocktails (Millipore Sigma, 4906845001) using a 2 mL Dounce homogenizer with 60 strokes per sample. Twenty-five microliters of homogenate were saved on ice as the input fraction. The remaining homogenate was centrifuged at 1,000 x g for 2 min at 4 °C, and the supernatant was incubated with anti-HA beads (Thermo Fisher Scientific, 66101) prewashed three times with cold PBS. Beads were incubated on a gentle rotator for 10 min at 4 °C, followed by three washes in cold PBS. Washes were performed by pipetting the beads three times per wash and using a magnetic rack to separate the beads from the supernatant. During the final wash, the bead suspension was transferred to a new low-protein-binding tube to improve lysosomal purity.

Input and lysosomal fractions were resuspended in ice-cold muscle lysis buffer containing protease and phosphatase inhibitors and incubated for 10 min at 4 °C. Lysosomal fractions were immobilized on a magnetic rack and transferred to new tubes twice. For immunoblotting, 0.08% of the input fraction and 15% of the lysosomal fraction (relative to whole tissue) were loaded per lane.

### Lysosomal metabolite extraction from mouse muscle

Mouse PDAC cells (1 x 10^5^ in 100 µL PBS) were implanted into the tail of the pancreas of muscle LysoTag (*Ckm-Cre; TMEM192-3xHA*) mice. Four weeks post-implantation, quadriceps muscles were immediately dissected from PDAC-bearing or control non-tumor-bearing mice on an ice-cold plastic dish. Muscle lysosomes were isolated using Lyso-IP as described above. For polar metabolite extraction, lysosomal fractions were resuspended in 500 µL ice-cold 80% methanol in LC-MS-grade water containing 500 nM isotope-labeled amino acids as internal standards (Cambridge Isotope Laboratories). Samples were vortexed for 10 min at 4 °C and centrifuged at 15,000 × g for 15 min at 4 °C. The supernatant containing polar metabolites was collected and dried under a stream of nitrogen. Dried samples can be stored at –80 °C. Immediately prior to analysis, samples were resuspended in 100 µL LC-MS-grade water.

### LC-MS-based muscle lysosome metabolite profiling

LC-MS analyses were performed on a Q Exactive benchtop Orbitrap mass spectrometer equipped with an Ion Max source and HESI II probe, coupled to a Dionex UltiMate 3000 ultra-high performance liquid chromatography system (Thermo Fisher Scientific, San Jose, CA). External mass calibration was performed using a standard calibration mixture every seven days. Acetonitrile was LC-MS HyperGrade (EMD Millipore), and all other solvents were LC/MS Optima grade (Thermo Fisher Scientific). The metabolite extraction solution (stored at –20 °C) consisted of 80:20 (v/v) methanol: water supplemented with 17 isotope-labeled amino acids at 500 nM each as internal standards (Cambridge Isotope Laboratories, MSK-A2-1.2).

For chromatographic separation, 2.5 µL of lysosomal IP sample was injected onto a SeQuant ZIC-pHILIC polymeric column (2.1 x 150 mm) with a guard column (2.1 x 20 mm); both columns had 5 µm particle size (EMD Millipore). The flow rate was 0.150 mL min^−1^, the column compartment was maintained at 25 °C, and the autosampler at 4 °C. Mobile phase A consisted of 20 mM ammonium carbonate with 0.1% ammonium hydroxide, and mobile phase B was 100% acetonitrile. The gradient program was as follows: 0-20 min, linear gradient from 80% to 20% B; 20-20.5 min, linear gradient from 20% to 80% B; 20.5-28 min, hold at 80% B. The mass spectrometer was operated in full-scan, polarity-switching mode. Spray voltage was set to 3.0 kV, the heated capillary to 275 °C, and the HESI probe to 350 °C. Sheath gas flow was 40 units, auxiliary gas 15 units, and sweep gas 1 unit. MS data were acquired over an m/z range of 70-1000 with a resolution of 70,000, AGC target of 1 x 10^6^, and maximum injection time of 20 ms.

### LC-MS metabolomics data processing and analysis

LC-MS-based metabolite identification and quantification were performed using XCalibur v4.0 software (Thermo Fisher Scientific) by matching accurate masses within a 5-ppm tolerance and retention times of authentic standards within a 0.5 min window. For relative quantification, the raw peak area of each metabolite was divided by the peak area of the corresponding isotope-labeled internal standard. Amino acids were normalized using their respective isotope-labeled standards, while other metabolites were normalized using the internal standard with the closest retention time. For comparison of PDAC versus control samples, amino acid abundances were expressed relative to the mean peak area of each amino acid in control samples.

### Measurement of hemoglobin A1C (HbA1c) levels

Whole-blood samples were collected in EDTA-containing tubes (Sarstedt, DE) from 6-week-old KP^−/−^C PDAC-bearing or control non-tumor bearing mice. 5 μL of plasma samples was assayed to measure HbA1c levels using mouse HbA1C assay kit (Crystal Chem, 80310) according to manufacture details.

### Blood glucose and plasma hormone measurements

Blood glucose levels were measured using a Contour glucose meter (Ascensia Diabetes Care). Whole-blood samples were collected in EDTA-containing tubes (Sarstedt, DE) from 6-week-old KP^−/−^C PDAC-bearing or control non-tumor bearing mice. Plasma insulin was sampled from fed or overnight fasted mice in the morning and was measured with an ultra-sensitive mouse insulin ELISA (Crystal Chem, 90080) according to manufacture details. Plasma glucagon was measured using a glucagon ELISA (R&D Systems, DGCG0) according to manufacture details.

### Proteasome activity assay

Muscle tissue samples from indicated genotypes were prepared in PIPES lysis buffer (50mM PIPES, 1mM MgCl_2_, 50mM NaCl_2_, 2mM EGTA, and 2mM fresh ATP) by homogenizing tissue using pellet pestle motor and spinning samples in tabletop centrifuge for 20 min at 4°C. Supernatant was taken and protein concentration measure with BCA assay (Pierce BCA Protein Assay Kit, Thermo Scientific, 23225). Briefly, 50 μg of protein was mixed at a 1:1 ratio with the Proteasome-Glo Chymotrypsin reagent (Promega, G8621) according to manufacture details. Bortezomib (Sigma-Aldrich, 5043140001), a proteasome inhibitor, was incubated with control samples as a readout for proteasome specific signal. Reactions were done in triplicate in a 96 well plate and read on Tecan Precious infiniteM200Pro microplate reader.

### Calpain activity assay

Enzymatic activity of Calpain was determined in duplicate in muscle homogenates using the substrates Suc-LLVY-AMC (Enzo Life Sciences, BML-P802) alone, or with the Calpain inhibitor Z-LL-CHO (Enzo Life Sciences). The fluorescence from homogenate incubated with and without the Calpain inhibitor was taken as the Calpain activity^104^. Briefly, muscle was homogenized in ice-cold lysis buffer (in mM: 20 HEPES, 10 NaCl, 1.5 MgCl, 1 DTT, 20 % glycerol and 0.1 % Triton X-100; pH 7.4) without protease inhibitors and centrifuged at 1,000 x g at 4 °C for 10 min. Supernatant protein concentrations were measured with BCA assay (Pierce BCA Protein Assay Kit, Thermo Scientific). Supernatants were then incubated in duplicate with the appropriate substrate at room temperature and fluorescence measured using on Tecan Precious infiniteM200Pro microplate reader with excitation and emission wavelengths of 380 nm and 460 nm for Calpain activity. Finally, proteolytic activity was normalized to total protein content.

### Fecal total protein and lipid quantification

For total fecal protein measurements, fecal samples (20 mg) were resuspended in 400 µL ice-cold phosphate-buffered saline (PBS), vortexed for 15 min, and sonicated for 5 min. Samples were centrifuged at 15,000 x g for 15 min at 4 °C, and protein concentration in the supernatant was determined using a Pierce BCA Protein Assay Kit (Thermo Fisher Scientific, 2322523225) according to the manufacturer’s instructions.

To measure total fecal lipid content, fecal samples (1,000 mg) were collected and homogenized in 1 mL 0.09% NaCl by vortexing at 4 °C for 10 min. Then, the samples were mixed with 1 mL of a 2:1 (v/v) chloroform: methanol solution and vortexed at 4 °C for 10 min. Finally, the samples were centrifuges at 1000 x g for 10 min at 4 °C. The lower organic phase was dried under a stream of nitrogen gas, and lipid content was determined gravimetrically by weighing the dried extracts.

### Fecal protease activity

Fecal samples (20 mg) were resuspended in 400 µL ice-cold phosphate-buffered saline (PBS), vortexed for 15 min at 4 °C, sonicated for 5 min, and centrifuged at 15,000 x g for 15 min at 4 °C. Supernatants were collected, and protein concentration was determined using the Pierce BCA Protein Assay Kit (Thermo Fisher Scientific, 23225). For protease activity measurements, 20 µL of clarified supernatant was incubated with 20 µL FITC-casein (Sigma-Aldrich, C0528) in a total reaction volume of 40 µL at 37 °C for 60 min. Reactions were terminated by the addition of 300 µL 10% (w/v) trichloroacetic acid, followed by centrifugation to remove precipitated protein. Fluorescence of the supernatant (excitation/emission: 485/535 nm) was measured using a Tecan Infinite M200 Pro microplate reader. Proteolytic activity was quantified as fluorescence intensity and normalized to total protein content.

### Fecal lipase activity

Fecal samples (20 mg) were resuspended in 400 µL ice-cold phosphate-buffered saline (PBS), vortexed for 15 min at 4 °C, and sonicated for 5 min. Samples were centrifuged at 15,000 x g for 15 min at 4 °C, and the supernatants were collected. Protein concentration was determined using the Pierce BCA Protein Assay Kit (Thermo Fisher Scientific, 23225). For lipase activity measurements, 20 µL of clarified supernatant was incubated with 40 µM EnzChek Lipase substrate (Thermo Fisher Scientific, E33955) in assay buffer containing 0.15 M NaCl, 20 mM Tris-HCl (pH 8.0), 0.0125% Zwittergent, and 1.5% fatty acid-free BSA, in a total reaction volume of 100 µL. Reactions were performed in black 96-well plates at 37 °C for 60 min. Fluorescence (excitation/emission: 485/535 nm) was measured using a Tecan Infinite M200 Pro microplate reader. Lipase activity was quantified as fluorescence intensity and normalized to total protein content.

### Fecal amylase activity

Fecal samples (20 mg) were resuspended in 400 µL ice-cold phosphate-buffered saline (PBS), vortexed for 15 min at 4 °C, and sonicated for 5 min. Samples were centrifuged at 15,000 x g for 15 min at 4 °C, and the supernatants were collected. Protein concentration was determined using the Pierce BCA Protein Assay Kit (Thermo Fisher Scientific, 23225). For amylase activity measurements, 20 µL of clarified supernatant was incubated with 10 µL EnzChek Amylase substrate (Thermo Fisher Scientific, E33651) in a total reaction volume of 100 µL in black 96-well plates at 37 °C for 60 min. Fluorescence (excitation/emission: 502/512 nm) was measured using a Tecan Infinite M200 Pro microplate reader. Amylase activity was quantified as fluorescence intensity and normalized to total protein content.

### Intestinal PBS wash-out and enzyme activity measurements

Intestinal segments (duodenum, jejunum, ileum, or colon) were carefully excised from the abdominal cavity and placed on ice. For all experiments shown, a 4 cm segment of proximal duodenum was analyzed. Luminal contents were gently flushed with 1 mL ice-cold phosphate-buffered saline (PBS) using a syringe, and the wash-through was collected for analysis of luminal enzyme activity. Samples were centrifuged at 15,000 x g for 30 min at 4 °C to remove insoluble debris, and the supernatants were retained. Protein concentration was determined using the Pierce BCA Protein Assay Kit (Thermo Fisher Scientific, 23225). For enzymatic activity measurements, 20 µL of clarified supernatant was incubated with either 10 µL EnzChek Amylase substrate (Thermo Fisher Scientific, E33651), 10 µL EnzChek Lipase substrate (Thermo Fisher Scientific, E33955), or 20 µL FITC-casein (Sigma-Aldrich, C0528), in a total reaction volume of 40 µL. Reactions were performed in black 96-well plates at 37 °C for 60 min. Fluorescence was measured using a Tecan Infinite M200 Pro microplate reader with excitation/emission settings appropriate for each substrate, as specified by the manufacturers. Enzyme activity was quantified as fluorescence intensity and normalized to total protein content.

### Oral Fat Tolerance Test (OFTT)

OFTT was perfumed as described previously^105^. Briefly, overnight fasted KP^−/−^C and non-tumor-bearing control mice were gavaged with olive oil (10 μL/g body weight). Blood was collected at indicated times for measurement of serum triglycerides (TGA) content. TGA in circulation were determined in serum samples using the mentioned kits (Abcam, ab65336) according to the manufacturer’s instruction.

### Protein Synthesis (SUnSET Analysis)

KP^−/−^C or non-tumor-bearing mice were starved for 3 h and then intraperitoneally injected with 0.040 μmol/g whole-body weight of puromycin (Sigma Aldrich, #P8833) in saline 30 minutes prior to euthanasia^79^. After euthanasia, skeletal muscle tissues were immediately snap freeze for total protein extraction or fixed in 10% neural formalin for Western blot or immunofluorescent staining, respectively.

### Histological analysis

Tissues were fixed overnight with neutral-buffered 10% formalin, paraffin-embedded, 5-µm sectioned, and stained with hematoxylin and eosin (H&E).

Tumor-bearing or non-tumor-bearing pancreas were also stained with Masson’s trichrome and Alcian blue methods to indicate tissue morphology using standard protocols. The analysis of histological changes was performed by using a light microscope.

### Histological quantification of adipocyte and myofiber cross-sectional area

Adipocyte and myofiber cross-sectional areas were quantified using ImageJ (NIH). Images were acquired at 20x magnification and calibrated using the embedded scale bar. Histological analyses were performed on tissue sections from four tumor-bearing mice and four control mice. For each animal, the largest tissue section was selected for analysis, with five non-overlapping fields quantified per section. Each dot represents an individual adipocyte or myofiber.

### Immunohistochemistry and immunofluorescence

Tissues were fixed in 10% neutral buffered formalin, paraffin embedded and sectioned at 4-5 µm thickness. Antigen retrieval was performed using Borg Decloaker RTU solution (BD1000G1; Biocare Medical) in a pressurized decloaking chamber (NxGen; Biocare Medical).

For immunohistochemistry (IHC), the following primary antibodies and dilutions were used: Ki67 (Biocare Medical, CRM325C, clone SP6; 1:50), cleaved Caspase-3 (Cell Signaling Technology, 9664, clone 5A1E; 1:800), PRSS2/trypsin (Abclonal, A19275; 1:100), Pancreatic lipase (PNLIP; Proteintech, 11209-1AP; 1:250; and Abclonal, A6396; 1:250), α-Amylase (Santa Cruz Biotechnology, sc46657, clone G10; 1:250), Insulin (Cell Signaling Technology, 4590; 1:200), Glucagon (Cell Signaling Technology, 2760; 1:200), and CK19 (Abcam, ab133496; 1:200). Primary antibodies were incubated overnight at 4 °C. Biotin-conjugated donkey anti-rabbit or anti-rat secondary antibodies were applied (1:500; Jackson ImmunoResearch), followed by signal amplification using the VECTASTAIN Elite ABC immunoperoxidase kit (PK6100; Vector Laboratories) and visualization with the SignalStain DAB substrate kit (8049; Cell Signaling Technology). All antibody dilutions were prepared in common antibody diluent (BioGenex, HK156-5K) supplemented with 10% normal goat serum (Thermo Fisher Scientific, 50062Z).

For immunofluorescence (IF) on paraffin-embedded sections, the following primary antibodies were used: mCherry (Novus Biologicals, NBP1-96752; 1:500), tdTomato (Rockland, 600-401-379; 1:200), α-Smooth muscle actin (Sigma, F3777; 1:250), puromycin (Sigma, MABE343; 1:1,000), Cathepsin B (Cell Signaling Technology, D1C7Y; 1:200), and anti-HA (MilliporeSigma, 11867423001; 1:500). Alexa Fluor-conjugated secondary antibodies (anti-rabbit 488 and anti-rat 568) were used for detection. Sections were mounted using ProLong Gold antifade mounting medium containing DAPI (Invitrogen). Images were acquired using a 40x objective on an Olympus FV1200 laser scanning confocal microscope.

### Histological quantification of insulin-positive pancreatic islets

Insulin-positive pancreatic islets were quantified using ImageJ (NIH). Images were acquired at 20x magnification and calibrated using the embedded scale bar. Histological analyses were performed on insulin-stained IHC sections from four tumor-bearing mice and four control mice. For each animal, the largest pancreatic tissue section was selected for analysis, and all insulin-positive islets within the section were quantified. Each dot represents an individual insulin-positive islet.

### Western blot

Snap-frozen tumor-bearing or control pancreas tissues were homogenized and scraped into RIPA lysis buffer (EMD Millipore, 20-188) containing protease and phosphatase inhibitors (Cell Signaling Technology, 5871), rocked for 15 min at 4 °C, and insoluble material was sedimented by centrifugation at 21,000 x g for 20 min at 4 °C. Homogenized muscle tissues were lysed in the muscle lysing buffer (20 mM Tris-HCl, 5 mM EGTA, 100 mM KCl, 1% TritonX 100, 50 mM NaF, 2 mM Sodium Orthovanadate, 3 mM Benzamide, 1 mM PMSF, 10 μg/mL Aprotinin; pH 7.4). Protein concentration was determined via BCA, and samples were mixed with LDS sample buffer (Thermo Fisher Scientific, NP0008) and 2.5% 2-mercaptoethanol then incubated at 95 °C for 5 min. Proteins were resolved by SDS-PAGE then transferred onto nitrocellulose membranes using a wet tank transfer system (Bio-Rad). Membranes were blocked in 5% non-fat milk in TBS-T and probed overnight at 4 °C with the appropriate antibody diluted in 5% BSA in TBS-T.

Primary antibodies were used including Vinculin (Cell Signaling Technology, 13901, clone E1E9V, 1:1,000), PRSS2 (Trypsin, Abclonal, A19275, 1:1,000), PNLIP (Pancreatic lipase, 11209-1AP-Proteintech, 1:1,000), Elastase (Abcam, ab21590, 1:1,000 dilution), α-Amylase (Santa Cruz, G10 clone-sc46657, 1:1,000), LC3B (Cell Signaling Technology, E7X4S, 1:1,000), phospho-ribosomal protein S6 Ser 235/236 (Cell Signaling Technology, 4858, 1/1,000), ribosomal protein S6 (Cell Signaling Technology, 2217, 1:1,000), phospho-p70 S6 kinase Thr389 (Cell Signaling Technology, 9205, 1:1,000), p70 S6 kinase (Cell Signaling Technology, 9202, 1:1,000), phospho-AMPK (Cell Signaling Technology, 2535, 1:1,000), AMPK (Cell Signaling Technology, 2532, 1:1,000), FOXO1 (Abclonal, A13862, 1:1,000), FOXO3A (Abclonal, A0102, 1:1,000), Anti-HA (Millipore-Sigma, 11867423001, 1:1,000), Cathepsin B (Cell Signaling Technology, D1C7Y, 1:1,000), Cathepsin L (R&D Systems, AF1515-SP, 1:1,000), ULK1 (Cell Signaling Technology, D8H5, 1:1,000), ATG5 (Cell Signaling Technology, D5F5U, 1:1,000), ATG7 (Sigma-Aldrich, A2856, 1:1,000), Puromycin (Sigma, MABE343, 1:10,000), EIF3F (Fortis Life Sciences, A303-005A-T, 1:1,000), Proteasome 20S Alpha7 subunit (ENZO Life sciences, BML-PW8110-0025, 1:1,000), Proteasome 19S Rpt3/S6b subunit (ENZO Life sciences, BML-PW8765-0025, 1:1,000), UBCJ2 (ENZO Life sciences, ENZ-ABS840-0100, 1:5,000), BNIP3 (Abcam, ab109362, 1:1,000), phosho-Acetyl-CoA Carboxylase I (S79) (Cell Signaling Technology, 6571S, 1:1,000), Acetyl-CoA Carboxylase I (Cell Signaling Technology, 3661S, 1:1,000), ATG3 (Cell Signaling Technology, 3415, 1:1,000), Calpain (Thermo Scientific, 9A4H8D3, 1:1,000), Lamp1 (Cell Signaling Technology, C54H11, 1:1,000), Lamp2 (Santa Cruz, H4B4, sc-18822, 1:1,000), Catalase (Cell Signaling Technology, D4P7B, 12980S, 1:1,000), Histone H3 (Cell Signaling Technology, D1H2, 4499T, 1:1,000), Cox IV (Cell Signaling Technology, 3E11, 4850T, 1:1,000), MuRF1 (ECM Biosciences, MP3401, 1:1,000), Atrogin-1 (ECM Biosciences, A2041P, 1:1,000).

Secondary antibodies were used including anti-rabbit IgG HRP-linked secondary antibody (Cell Signaling Technology, 7074, 1:5,000) and anti-mouse IgG HRP-linked secondary antibody (Cell Signaling Technology, 7076, 1:5,000).

Primary antibodies were diluted in 5% BSA in TBS-T. The following day, membranes were washed three times with TBST on a rocker for 10 min. Secondary antibodies were applied for 60 min. Membranes were then washed again three times with TBST for 10 min, and signal was detected with ECL using film. In cases where re-probing of the same membrane was required, membrane was incubated with hydrogen peroxide solution 30% (w/w in water) (Sigma-Aldrich, H1009) for 1 h at 37 °C to inactivate HRP. Western blots were quantified by densitometry using Image J and Fiji; data are normalized to the loading control from the same gel.

### RNA sequencing

20-30 mg homogenized tumor-bearing pancreas tissues (6-week-old KP^−/−^F PDAC or *Atg7* mKO KP^−/−^F PDAC mice) were used in the total RNA extraction. RNA was extracted using Zymogen kit (R1054). RNA was diluted to a concentration of 2 ng/mL and 5 mL of each sample was transferred to a 96-well plate, then poly-A enrichment and cDNA synthesis were performed in parallel, followed by library preparation using the Nextera XT kit and paired-end sequencing on an Illumina NextSeq 500. Reads were mapped to the mm10 genome using STAR (v2.7.8a) and default parameters. Reads that mapped to transcripts were counted using feature Counts (with parameters-M-O-fraction-primary). Normalization and differential gene expression analysis were performed using the edgeR (v4.0.1) package in R. Gene set enrichment analysis was performed using the fgsea (v1.28.0) package in R with gene signatures from the Molecular Signatures Database.

### Statistics and reproducibility

Sample sizes, reproducibility, and statistical tests used for each figure are reported in the figure legends. All graphs were generated in GraphPad Prism 10 (GraphPad Software). Metabolite concentrations from plasma and TIF were subjected to multivariate and univariate statistical analysis using Metaboanalyst 6.0^106^ as described previously^46^. Briefly, metabolite concentrations were log-transformed. Metabolites with >50% missing values across all samples were removed. For metabolites retained in the dataset, missing values were imputed using one-fifth of the lowest detected positive value to approximate the detection limit. For multivariate analysis, hierarchical clustering was performed using Euclidean distance and Ward’s linkage algorithm. Univariate comparisons between groups were performed using heteroscedastic (unequal variance) tests. Metabolite differences were considered significant at a fold change >2 and a raw P-value <0.05. Using these fold-change and adjusted P-value criteria, the number of differentially abundant metabolites was quantified.

## Illustrations

Experimental schema were generated using BioRender (https://biorender.com/).

## Data availability

All data that support the findings of this study are available upon request from the corresponding author.

## Acknowledgements

We thank members of the Vander Heiden laboratory for helpful discussions and the Koch Institute’s Robert A. Swanson (1969) Biotechnology Center for technical support, specifically Dr. Aurora Burds Connor in the Preclinical modeling, Preclinical Imaging & Testing, Barbara K. Ostrom (1978) Integrated Genomics and Bioinformatics, Hope Babette Tang (1983) Histology, and Metabolite Profiling facilities. We also thank the MIT Division of Comparative Medicine staff for help with colony maintenance and animal care. This work was supported in part by the Koch Institute Support (core) Grant P30-CA014051 from the National Cancer Institute. We would also like to thank all members of the Vander Heiden lab for helpful discussions. Y.G. is supported by a postdoctoral fellowship from the Ludwig Center at MIT’s Koch Institute for Integrative Cancer Research. S.S. acknowledges support from the Damon Runyon Cancer Research Foundation (DRG-2367-19). K.M.E. was supported by a Boehringer Ingelheim Fonds MD fellowship. K.L.A. was supported by the National Science Foundation (DGE-1122374) and National Institutes of Health (NIH) (F31CA271787, T32GM007287). B.T.D. was supported by F30HL156404 from NHLBI and T32GM007753 from NIGMS. M.G.V.H. acknowledges current and past support from the Hale Center at Dana-Farber, the Lustgarten Foundation, the MIT Center for Precision Cancer Medicine, the Ludwig Center at MIT, and the NCI (R35CA242379, P30CA14051).

## Author Contributions

Y.G. and M.G.V.H. conceived the project. Y.G. designed and performed experiments, assembled the manuscript, and prepared the figures. S.S. performed implantation of mouse PDAC cells into Nu/J mice, conducted pancreatic injections of LUAD and HCC cells into the pancreas, maintained and shared the KPC PDAC GEMM. K.M.E. carried out the D20 diet and performed ^15^N-yeast protein labeling, helped ^15^N-yeast protein gavage, and assisted with high and low protein-diet studies. A.M.B. contributed to protein and free amino acid diet experiments in KP^−/−^C PDAC GEMMs. K.L.A. analyzed LC-MS metabolite data from plasma and MIF and also maintained C57BL/J6 mice. G.E. and Ö.H.Y. provided inducible colorectal GEMMs and performed implantation of colorectal organoids into the rectum, pancreas, and liver. B.T.D. analyzed RNA-sequencing data. H.S. and Ö.H.Y. provided the LysoTag transgenic mouse line and assisted with optimization of muscle Lyso-IP. V.L.T., S.H., E.Ö., D.A.S., and Y.K.K. contributed to the analysis of KPC and KP^−/−^C PDAC GEMMs. T.K. and M.W. performed LC-MS experiments. E.W. provided *Atg7^fl/fl^* mouse. W.A.F-P., W.M.R., and T.J. generated and provided the *Pdx1-P2A-FlpO* PDAC GEMM model. B.M.W. and J.A.N. acquired funding and provided input. M.G.V.H. supervised the study and acquired funding. Y.G. and M.G.V.H. wrote the manuscript with input from all authors.

## Lead contact

Further information and requests for resources and reagents should be directed to and will be fulfilled by Matthew G. Vander Heiden.

