## Extended Data figures and legends for "Pancreatic cancer-associated organ dysfunction promotes muscle autophagy and contributes to peripheral tissue wasting"

#### Extended Data Figure Legends

**Extended Data Figure 1 (Related to Figure 1): Mouse models of PDAC exhibit tissue wasting that begins with early disease. a-h.** Whole-body weights (a), food intake (b), and tissue weight of pancreas (c), liver (d), perigonadal adipose tissue (AT) (e), quadriceps (f), gastrocnemius (g), and soleus (h) muscle normalized to total body weight in control and $KP^{-/-}C$ mice (*n* = 10 except for *n* = 8 in panel b). **i.** Gross images and H&E tissue image of gastrocnemius muscle from 10-week-old control and $KP^{-/-}C$ mice as indicated. **j.** Survival of KPC mice (*n* = 15 KPC). **k-q.** Whole-body weights (k) (*n* = 11 control, *n* = 15 KPC), blood glucose levels (l) (*n* = 7 control, *n* = 7 KPC), and tissue weights of liver (m), pancreas (n), perigonadal AT (o), subcutaneous ATs (p), quadriceps, gastrocnemius, and soleus muscle (q) normalized to total-body weight in end-stage KPC mice relative to age-matched controls (*n* = 8 control, *n* = 15 KPC for all tissue weight data). Both male and female mice used in experiments. Statistical analysis was performed using unpaired two-sided *t*-tests, data are mean ± S.D and *n* represents the number of mice analyzed. Scale bars: 100 μm.

**Extended Data Figure 2 (Related Figure 1): Pancreatic endocrine dysfunction accompanies PDAC in mice. a.** Schematic showing approach to quantify metabolites in plasma collection from control and PDAC mice. **b.** Partial least squares discriminant analysis comparing metabolite profiles measured in plasma 6-week-old control and $KP^{-/-}C$ mice (*n* = 3 control, *n* = 4 $KP^{-/-}C$). **c.** Hierarchical clustering of metabolites measured in plasma from 6-week-old control and PDAC mice. Columns of the heatmap were z-score normalized (*n* = 3 control, *n* = 4 $KP^{-/-}C$). **d.** Levels of BCAAs measured in plasma of fed and 16h fasted 6-week-old control and $KP^{-/-}C$ mice as indicated. **e.** Blood glucose measured over time in $KP^{-/-}C$ mice and littermate controls (*n* = 10 except *n* = 8 for 7-10-week-old $KP^{-/-}C$ mice). **f.** Hemoglobin A1c levels measured in blood from 6-week-old control and $KP^{-/-}C$ mice (*n* = 5). **g.** Plasma glucagon levels measured in plasma of fed and 16h fasted 6-week-old control and $KP^{-/-}C$ mice as indicated (*n* = 5 control, *n* = 4 $KP^{-/-}C$). **h.** Immunohistochemistry (IHC) staining for CK19, glucagon, and insulin in pancreas tissue from 6-week-old control and $KP^{-/-}C$ (PDAC) mice as indicated. **i.** Quantification of insulin+ islets from the IHC analysis shown in h (*n* = 7 control, *n* = 6 $KP^{-/-}C$ mice). Both male and female mice used. Statistical analysis was performed using unpaired two-sided *t*-tests, data are mean ± S.D and *n* represents the number of mice analyzed. Scale bars: 100 μm.

**Extended Data Figure 3 (Related to Figure 1): Neither high-sugar nor high-protein diets mitigate PDAC-induced muscle wasting. a.** Schematic image for assessing how exposing mice to high glucose in water impacts on survival, tumor growth, and tissue wasting in mice with PDAC. **b.** Survival of $KP^{-/-}C$ mice (PDAC) when exposed to regular drinking water or water with 20% glucose (D20) as in a (*n* = 5 $KP^{-/-}C$ on regular water, *n* = 6 $KP^{-/-}C$ on D20 water). **c-g.** Whole-body weight (c) and tissue weights of liver (d), tumor-bearing pancreas (e), subcutaneous and perigonadal ATs (f), quadriceps, gastrocnemius, and soleus muscle (g) normalized to total body weight in $KP^{-/-}C$ mice at endpoint after exposure to regular drinking water or D20 as in a (*n* = 5 $KP^{-/-}C$ on regular water, *n* = 6 $KP^{-/-}C$ on D20 water). **h.** Schematic image for assessing how exposing mice to high glucose in water together with diets containing normal protein (1X) and increased protein (2X) impacts tumor growth, and tissue wasting in mice with PDAC. **i-l.** Tissue weights of tumor-bearing pancreas (i), liver (j), subcutaneous and perigonadal ATs (k), quadriceps and gastrocnemius as well as soleus muscle (l) normalized to total body weight in $KP^{-/-}C$ mice at endpoint after exposure to regular drinking water or D20 with different protein concentrations as described in h (*n* = 8 $KP^{-/-}C$ on regular water and 1X Protein diet, *n* = 7 $KP^{-/-}C$ on regular water and 1X Protein diet, *n* = 8 $KP^{-/-}C$ on D20 and 1X Protein diet, *n* = 8 $KP^{-/-}C$ on regular water and 2X Protein diet, *n* = 8 $KP^{-/-}C$ on D20 and 2X Protein diet). **m.** Experimental schematic depicting how exposure to modified AIN-93G diets containing 0.5X, 1X, or 2X protein affects survival, tumor growth, and tissue wasting in $KP^{-/-}C$ PDAC mice. **n.** Hierarchical clustering of amino acid concentrations measured in plasma from control and $KP^{-/-}C$ mice exposed to diets with different protein concentrations as described in m. Columns of the heatmap were z-score normalized. (*n* = 7 $KP^{-/-}C$ on 0.5X Protein diet, *n* = 11 $KP^{-/-}C$ on 1X Protein diet, *n* = 7 $KP^{-/-}C$ on 2X Protein diet. *n* = 7 control mice on 0.5X Protein diet, *n* = 10 littermate control mice on 1X Protein diet, and *n* = 6 littermate control mice on 2X Protein diet). **o.** Survival of $KP^{-/-}C$ mice when exposed to different protein diet as in m (*n* = 19 $KP^{-/-}C$ on 0.5X Protein diet, *n* = 13 $KP^{-/-}C$ on 1X Protein diet, *n* = 18 $KP^{-/-}C$ on 2X Protein diet). **p-t.** Whole-body weight (p) and tissue weights of liver (q), pancreas (r), perigonadal AT (s), quadriceps, gastrocnemius, and soleus muscle (t) normalized to total body weight in $KP^{-/-}C$ mice at endpoint after exposure to different protein diets as in m. (*n* = 7 $KP^{-/-}C$ and *n* = 10 control mice on 0.5X Protein diet; *n* = 10 $KP^{-/-}C$ and *n* = 8 control mice on

1X Protein diet; *n* = 7 $KP^{-/-}C$ and *n* = 9 control mice on 2X Protein diet). Both male and female mice used in those experiments. P-values on survival experiments in b and p were calculated with Gehan-Breslow-Wilcoxon test. All other statistical analysis was performed using unpaired two-sided *t*-tests, data are mean ± S.D and *n* represents the number of mice analyzed.

**Extended Data Figure 4 (Related to Figure 2): Early tissue wasting does not occur in mice with flank PDAC tumors and is independent of the adaptive immune system**. **a.** Schematic showing implantation of murine $KP^{-/-}C$ PDAC cancer cells into the flank (subcutaneous) of syngeneic mice to generate tumors. **b.** Trichrome and alcian blue staining of tumor tissues harvested from the mice with PDAC cells implanted in the flank as in a, and in tumor tissue from mice with PDAC cells orthotopically in the pancreas as indicated. Pancreas tissue from mice with PBS-injected into the pancreas is also shown as a control (images are representative of *n* = 3 animals analyzed for each group). **c-e.** Whole-body weight (c), food intake (d), and blood glucose levels (e) measured in mice without (control, PBS injected) and with flank tumors derived from PDAC cells as in a (*n* = 5). **f-i.** Weights of flank PDAC tumors (f), pancreas (g), liver (h) as well as subcutaneous and perigonadal ATs (i) normalized to whole-body weights (*n* = 5). **j.** H&E staining of perigonadal AT from control and flank PDAC mice, and quantification of adipocyte area (*n* = 129 adipocytes analyzed from histology sections of *n* = 3 control, *n* = 3 flank PDAC). **k.** H&E staining of gastrocnemius muscle tissue from control and flank PDAC mice, and quantification of myofibers area (*n* = 264 myofibers analyzed from histology sections of *n* = 3 control and *n* = 3 flank PDAC mice). **l.** Tissue weights of quadriceps, gastrocnemius, and soleus muscle normalized to whole-body weights from control and flank PDAC mice as in a (*n* = 5). **m.** Schematic showing orthotopic implantation of murine KPC PDAC cells into the pancreas tail of 12-week-old male Nu/J mice. **n-r.** Whole-body weight (n) and tissue weights of pancreas (o), subcutaneous (p) and perigonadal ATs (q), as well as quadriceps, gastrocnemius, and soleus muscle (r) normalized to total body weight 4-weeks after injection of PBC (control) or implantation of PDAC cells into the pancreas as in m. (*n* = 8 control, *n* = 4 PDAC). Statistical analysis was performed using unpaired two-sided *t*-tests, data are mean ± S.D and *n* represents the number of mice analyzed. Scale bars: 100 μm.

**Extended Data Figure 5 (Related to Figure 2): The presence of a tumor in the pancreas leads to tissue wasting in mice. a.** Schematic showing approach to implant KPC PDAC cells to form tumors in different tissue locations in syngeneic 12-week-old female C57BL/6J mice. This includes dual injection into both the liver and pancreas to form tumors in each site (Dual injection, O+I). **b.** Representative images of PDAC tumors forming in the pancreas, liver, or pancreas and liver (Dual injection) from experiments outlined in a. **c-g.** Whole-body weights (c). tissue weights of liver (d), pancreas (e), perigonadal AT (f), and quadriceps, gastrocnemius, and soleus muscle (g) normalized to whole-body weights from the indicated mice with tumors injected into different tissue locations as outlined in a at endpoint. (*n* = 3 PBS-injected pancreas control, *n* = 3 orthotopic injection, *n* = 4 hepatic injection, and *n* = 3 hepatic and orthotopic injection). **h.** Schematic showing approach to implant different murine tumor cells obtained from models of hepatocellular carcinoma (HCC), lung adenocarcinoma (LUAD), and KPC PDAC into the pancreas tails of syngeneic 12-week-old female C57BL/6J mice. **i-m.** Whole-body weights, tissue weights of liver (j), pancreas (k), subcutaneous and perigonadal ATs (l), and quadriceps, gastrocnemius, and soleus muscle (m) normalized to whole-body weights from mice with the indicated cancer cell type injected into the pancreas as outlined in h at endpoint. (*n* = 5 control, *n* = 4 HCC-injected mice, *n* = 4 LUAD-injected mice, *n* = 4 PDAC-injected mice). **n.** Schematic showing approach to implant murine rectal tumor organoids into the pancreas tail of syngeneic 12-week-old male C57BL/6J mice. **o-s.** Whole-body weights (o), weight of pancreas (p), liver (q), subcutaneous and perigonadal ATs (r), and quadriceps, gastrocnemius, and soleus muscle (s) normalized to total body weight from control mice or mice with rectal organoid-derived tumors in the pancreas (*n* = 5 control, *n* = 4 Rectal tumor organoid-injection). **t.** Schematic showing approach to implant murine rectal tumor organoids into the rectum (R) or liver (via intrasplenic injection, ISL) of syngeneic 12-week-old male C57BL/6J mice. **u-z.** Whole-body weight after 6-weeks, and weight of colon (v), liver (w), pancreas (x), perigonadal AT (y), and quadriceps, gastrocnemius, and soleus muscle (z) normalized to total body weight from control mice or mice with rectal organoid-derived tumors implanted into the indicated sites as described in t at endpoint. (*n* = 5). Statistical analyses were performed using one-way ANOVA with Tukey's post hoc test for data in panels c-m. For all other comparisons, unpaired two-sided *t*-tests were used. Data are presented as mean ± S.D., and *n* denotes the number of mice analyzed.

**Extended Data Figure 6 (Related to Figure 2): Pancreatic cancer causes pancreatic exocrine dysfunction. a.** Schematic showing experimental strategy to determine if a corn starch diet (CSD) or a high starch diet (HSD) affects blood glucose in mice with PDAC. **b.** Pie chart showing relative nutrient composition of a mouse purified diet (AIN-93G). **c.** % digestible and non-digestible (resistant) starch (w/w) in standard AIN-93G, CSD and HSD diets. **d.** Blood glucose measurements from 6-week-old $KP^{-/-}C$ mice (PDAC) or littermate controls after an overnight fast and 24h of exposure of an AIN-93G or HSD as indicated ($n$ = 3). **e-h.** Following oral gavage of $^{15}N$-*Spirulina* diet to 6-week-old control or $KP^{-/-}C$ (PDAC) mice, assessment of % $^{15}N$-labeling (M+1) in blood over time of tyrosine (e), leucine (f), serine (g), and glycine (h) ($n$ = 4). **i-l.** Following oral gavage of $^{15}N$-labeled protein from yeast to 6-week-old control or $KP^{-/-}C$ (PDAC) mice, assessment of % $^{15}N$-labeling (M+1) over time of tyrosine (i), leucine (j), serine (k), and glycine (l) in plasma proteins ($n$ = 4). Both male and female mice were used. Statistical analyses were performed using one-way ANOVA with Tukey's post hoc test for d. For all other comparisons, unpaired two-sided *t*-tests were used. Data are presented as mean ± S.D., and $n$ denotes the number of mice analyzed.

**Extended Data Figure 7 (Related to Figure 2): Supplementing pancreatic enzymes restores pancreatic exocrine function and mitigates early PDAC-induced tissue wasting. a.** Schematic showing experimental strategy to assess how pancreatic enzyme supplementation (PES) in diet compared to a control AIN-93G diet affects mice with PDAC. **b-f.** Quantification of total protein (b), protease activity (c), lipid amount (d), lipase activity (e), and amylase activity (f) in feces from mice with $KP^{-/-}C$ PDAC tumors implanted in the pancreas tail fed a control or PES diet as in a ($n$ = 6 PDAC on control diet, $n$ = 7 PDAC on PES diet). **g.** Quantification of intestinal protease activity in mice with PDAC tumors implanted in the pancreas tail and fed a control or PES diet for 15-days ($n$ = 6). **h-n.** Whole-body weight (h), daily food intake (i), blood glucose levels (j) and weights of liver (k), pancreas (l), perigonadal AT (m), quadriceps, gastrocnemius, and soleus skeletal muscle (n) normalized to whole-body weight in mice with PDAC tumors implanted in the pancreas tail fed a control or PES diet for 2-weeks ($n$ = 12 PDAC on control diet, $n$ = 10 PDAC on PES diet). **o.** H&E staining of the indicated mouse tissues collected from mice with PDAC tumors implanted in the pancreas tail fed a control or PES diet for 2-weeks. **p.** Schematic showing experimental strategy to assess how PES delivered by daily oral gavage affects mice with PDAC. **q-u.** Whole-body weight (q) and tissue weights of liver (r), pancreas (s), perigonadal AT (t), quadriceps, gastrocnemius, and soleus muscle (u) normalized to total body weight in 4-week-old $KP^{-/-}C$ mice with PDAC tumors that were treated with vehicle (PBS) or PES via oral gavage for 2-weeks as in p (*n* = 13 PBS, *n* = 18 PES ). **v-z.** Intestinal protease activity (v), fecal total protein amount (w), fecal protease activity (x), fecal lipase activity (y), and fecal amylase activity (z) in 4-week-old $KP^{-/-}C$ mice with PDAC tumors that were treated with vehicle (PBS) or PES via oral gavage (*n* = 10 PBS, *n* = 9 PES). Both male and female mice were used. For all comparisons, unpaired two-sided *t*-tests were used. Data are presented as mean ± S.D., and *n* denotes the number of mice analyzed. Scale bars: 100 µm.

**Extended Data Figure 8 (Related to Figure 2) Supplementing pancreatic enzymes does not affect tissue size in non-tumor-bearing mice. a.** Schematic showing experimental strategy to assess effects of pancreatic enzyme supplementation (PES) included in the diet on wildtype C57BL/6J mice. **b-h.** Food intake (b), whole-body weight (c) and weight of liver (d), pancreas (e), perigonadal AT (f), quadriceps, gastrocnemius, and soleus (g) normalized to total body weight of 12-week-old mice fed a control or PES containing diet as in a (*n* = 5 control diet, *n* = 6 PES diet). **h-j.** Quantification of total protein (h) and protease activity (i) in feces of 12-week-old mice fed a control or PES containing diet for 4-weeks as in a (*n* = 5 control diet, *n* = 6 PES diet). **j-l.** Fecal protease activity measured in stool collected from 12-week-old mice fed a control or PES containing diet for 3-weeks (j), 2-weeks (k), and 1-week (l) (*n* = 4 control diet, *n* = 4 PES diet). **m-n.** Quantification of lipase activity (m) and amylase activity (n) in feces of 12-week-old mice fed a control or PES containing diet for 4-weeks as in a (*n* = 5 control diet, *n* = 6 PES diet). **o-q.** Quantification of intestinal protease activity (o), lipase activity (p), and amylase activity (q) in 12-week-old mice fed a control or PES containing diet for 4-weeks (*n* = 5 control diet, *n* = 6 PES diet). Both male and female mice were used. Statistical analysis was performed using unpaired two-sided *t*-tests, data are mean ± S.D. *n* represents the number of mice analyzed.

**Extended Data Figure 9 (Related to Figure 3): Assessment of glucose metabolism in muscle from PDAC-bearing and control mice. a.** Schematic showing experimental approach for use of intravenous (IV) [U-$^{13}$C]-glucose infusion to assess glucose fate in muscle of PDAC bearing and its littermate control mice. **b-c.** Fractional labeling of fully labeled glucose (m+6) (b) and unlabeled glucose (m+0) (c) in the plasma of 6-weeks-old male control and KP$^{-/-}$C (PDAC) mice following infusion with [U-$^{13}$C]-glucose at a rate of 0.4 mg/min for 6 hours (*n* = 3). **d-o.** Fractional labeling of lactate (d), pyruvate (e), serine (f), serine (m+3) (g), alanine (h), aspartate (i), malate (j), glutamine (k), citrate (l), fumarate (m), succinate (n), and α-ketoglutarate (α-KG) (o) in gastrocnemius muscle from 6-weeks-old male control and KP$^{-/-}$C (PDAC) mice following infusion with [U-$^{13}$C]-glucose (*n* = 3). **p-s.** Total ion counts (TIC) measured for aspartate (p), α-KG (q), lactate (r), and the indicated metabolites (s) in gastrocnemius muscle from 6-weeks-old male control and KP$^{-/-}$C (PDAC) mice following infusion with [U-$^{13}$C]-glucose (*n* = 3). Statistical analysis was performed using unpaired two-sided *t*-tests, data are mean ± S.D and *n* represents the number of mice analyzed.

**Extended Data Figure 10 (Related to Figure 3): Assessment of ubiquitin proteosome system and calpain protease activity in muscle from PDAC-bearing mice. a-b.** Proteosome activity measured in muscle lysates from 4-week-old (a) (*n* = 6 KP$^{-/-}$C, *n* = 7 control) or 6-week-old (b) (*n* = 30 KP$^{-/-}$C, *n* = 36 control) control or (PDAC) mice without (DMSO) or with addition of the proteasome inhibitor bortezomib as indicated (b). **c.** Western blot analysis of the ubiquitin proteosome system (UPS)-related proteins ubiquitin-conjugating ligase E2 (Ubcj2), the 19S proteasome protein Rpt3, the 20S proteasome protein Alpha-7, and the muscle specific ubiquitin E3 ligases MuRF-1 and Atrogin-1 in muscle lysates from control and KP$^{-/-}$C mice as indicated. Vinculin was also assessed as a loading control **(***n* = 10 KP$^{-/-}$C, *n* = 5 control). **d-f.** Quantification of Ubcj2 (d), Alpha-7 (e), and Rpt3 (f) protein levels from western blots shown in c. (*n* = 10 KP$^{-/-}$C, *n* = 5 control). **g-h.** Western blot analysis for ubiquitin in muscle lysates from control and KP$^{-/-}$C mice as indicated (g), and quantifications of polyubiquitin proteins (h) from the western blot shown in g (*n* = 9 KP$^{-/-}$C, *n* = 7 control). **i.** Calpain protease activity measured in muscle lysate from 6-week-old control or KP$^{-/-}$C (PDAC) mice without (control) or with addition of the calpain inhibitor J61766.LB0 as indicated (*n* = 17 KP$^{-/-}$C, *n* = 22 control). **j-k.** Western blot analysis for calpain in muscle lysates from control and KP$^{-/-}$C mice as indicated (k), and quantifications of calpain (k) from the western blot shown in j (*n* = 10 KP$^{-/-}$C, *n* = 5 control). Both male and female mice were used, and gastrocnemius muscle was the source of muscle lysates. Both proteasome and calpain activity assays were performed with fresh lysates. For all comparisons, unpaired two-sided *t*-tests were used. Data are presented as mean ± S.D., and *n* denotes the number of mice analyzed.

**Extended Data Figure 11 (Related to Figure 3): Assessment of lysosomal amino acids muscle of mice with PDAC. a.** Schematic showing experimental approach to use immunoprecipitation (IP) to assess lysosomal material from *Ckm-Cre; TMEM192-HA* (muscle Lyso-Tag) mice. **b.** Immunofluorescence (IF) staining for TMEM192-HA protein in quadriceps muscle tissue from 12-week-old fed or overnight fasted mice as indicated (*n* = 2). **c.** Western blot analysis to assess the indicated markers of different subcellular compartments in whole-tissue lysates (WTL) and IP fractions of quadriceps muscle from 12-week-old fed or overnight fasted mice, with or without TMEM192-HA expression as indicated. **d.** Schematic showing experimental approach for use of lyso-IP of skeletal muscle to assess lysosomal metabolites. **e.** IF staining for TMEM192-HA and cathepsin B in quadriceps muscle tissue from mice without (control) or with KPC PDAC cells implanted in the pancreas tail (*n* = 4). **f.** Amino acid levels measured in lyso-IP material collected from quadriceps muscles as in d from PDAC-bearing or control mice (*n* = 4). Statistical analysis was performed using unpaired two-sided *t*-tests, data are mean ± S.D and *n* represents the number of mice analyzed. Scale bars: 200 µm.

**Extended Data Figure 12 (Related to Figure 3): Single amino acid-deficiency in the diet can cause tissue wasting. a.** Schematic showing experimental approach to assess the effects of amino acid deficient diets on mouse tissue weights. **b.** % whole-body weight change of 12-week-old C57BL/6J mice after exposure to the indicated diet for 4-weeks. (*n* = 12 control diet, *n* = 7 -tryptophan (-Trp), *n* = 8 on -branched-chain amino acids (-BCAA), *n* = 13 -methionine (-Met), *n* = 8 -lysine (-Lys), *n* = 10 -serine and glycine (-Ser-Gly)). **c-i.** Tissue weights of pancreas (c), liver (d), subcutaneous (e) and perigonadal AT (f), quadriceps (g), gastrocnemius (h), and soleus (i) muscle normalized to body weights from 12-week-old C57BL/6J mice after exposure to the indicated diet for 4-weeks. (*n* = 12 control diet, *n* = 7 -Trp, *n* = 8 -BCAA, *n* = 13 -Met, *n* = 8 -Lys, *n* = 10 -Ser-Gly). Malle and female mice were used. Statistical analyses were performed using one-way ANOVA with Tukey's post hoc test.

**Extended Data Figure 13 (Related to Figure 3): Supplementing free amino acids in the diet improves skeletal muscle mass in mice with PDAC. a.** Schematic showing experimental approach to use diets with different levels of free amino acids (FAA) to assess effects on tissue weights and survival. **b-f.** Whole-body weight (b) and tissue weights of pancreas (c), liver (d), perigonadal and subcutaneous ATs (e), quadriceps, gastrocnemius, and soleus muscle (f) normalized to total body weight of control or $KP^{-/-}C$ (PDAC) mice at end point that were fed a diet with FAA amino acids that match levels in a AIN-93G purified diet (1x), or a diet with low (0.5X) or high (2X) levels of FAA, beginning at 4-weeks of age (*n* = 24 0.5X FAA, *n* = 16 1X FAA, *n* = 14 2X FAA, *n* = 18 $KP^{-/-}C$ PDAC 0.5X FAA, *n* = 21 $KP^{-/-}C$ PDAC 1X FAA, *n* = 16 $KP^{-/-}C$ PDAC 2X FAA). **g.** Survival of $KP^{-/-}C$ mice exposed to the indicated diets (*n* = 31 $KP^{-/-}C$ 0.5X FAA, *n* = 29 $KP^{-/-}C$ 1X FAA, *n* = 32 $KP^{-/-}C$ on 2X FAA). Both male and female mice used. Statistical analyses were performed using one-way ANOVA with Tukey's post hoc test, data are mean ± S.D and *n* represents the number of mice analyzed. P-values on survival experiments were calculated with Gehan-Breslow-Wilcoxon test.

**Extended Data Figure 14 (Related to Figure 4): Loss of *Atg7* in muscle limits PDAC tumor growth and tumor-associated tissue wasting. a.** Schematic showing experimental design for implantation of murine KPC PDAC cells into the pancreas tail of mice without and with loss of *Atg7* in muscle. **b-f.** Whole-body weight (b) and tissue weights of liver (c), pancreas (d), perigonadal AT (e), quadriceps, gastrocnemius, and soleus muscle (f) normalized to total body weight of 8-week-old male *Ckm-Cre* (control) and $Atg7^{fl/fl}$*; Ckm-Cre* (*Atg7* mKO) mice 4-weeks after tumor implantation in the pancreas as described in a (*n* = 7). **g.** Schematic showing experimental design for implantation of murine KPC PDAC cells into the subcutaneous flank of mice without and with loss of *Atg7* in muscle. **h-m.** Whole-body (h) and tissue weights of liver (i), pancreas (j), flank tumors (k), perigonadal AT (l), quadriceps, gastrocnemius, and soleus muscle (m) normalized to total body weight of 8-week-old male *Ckm-Cre* (control) and $Atg7^{fl/fl}$*; Ckm-Cre* (*Atg7* mKO) mice 4-weeks after tumor implantation in the flank as described in g (*n* = 6 control and *n* = 5 *Atg7* mKO mice). Statistical analysis was performed using unpaired two-sided *t*-tests, data are mean ± S.D and *n* represents the number of mice analyzed.

**Extended Data Figure 15 (Related to Figure 4): Impaired muscle Atrogin-1 modestly affects PDAC progression and tissue wasting**. **a.** Schematic showing experimental design for generation of mice with a conditional loss of function *Atrogin-1* allele. We genotyped animals by PCR to obtain wild type mice and mice homozygous for the *Atrogin-1* allele *(Atrogin-1$^{fl/fl}$)*. **b.** PCR genotyping to assess the *Atrogin-1* allele in different tissues of mice without or with a *Ckm-Cre* allele as indicated. **c.** Schematic showing experimental design for implantation of murine KPC PDAC cells into the pancreas tail of mice without and with loss of functional *Atrogin-1* in muscle. **d-g.** Whole-body weight (d) and tissue weights of pancreas (e), perigonadal AT (f), quadriceps, gastrocnemius, and soleus muscle (g) normalized to total body weight of 8-week-old male *Ckm-Cre* (control) and *Atrogin-1$^{fl/fl}$; Ckm-Cre* (*Atrogin-1* mLOF) 4-weeks after tumor implantation in the pancreas as described in c. (*n* = 4 *Ckm-Cre*, *n* = 3 *Atrogin-1* mLOF) Statistical analysis was performed using unpaired two-sided *t*-tests, data are mean ± S.D and *n* represents the number of mice that were analyzed.

**Extended Data Figure 16 (Related to Figure 4): Assessment of how muscle autophagy influence plasma metabolites in mice with PDAC**. **a.** Schematic showing experimental design for determining how muscle autophagy affects plasma metabolite levels in mice with KP$^{-/-}$F PDAC. **b.** Hierarchical clustering of plasma metabolites measured in control mice (*Pdx-1-P2A-FlpO; Trp53$^{Frt/Frt}$ ; Ckm-Cre), Atg7* muscle knock-out mice (*Pdx-1-P2A-FlpO; Trp53$^{Frt/Frt}$ ; Ckm-Cre*; *Atg7$^{fl/fl}$* (*Atg7* mKO)), KP$^{-/-}$F PDAC mice (*Pdx-1-P2A-FlpO; Kras$^{FSF-G12D/+}$; Trp53$^{Frt/Frt}$*), and *Atg7* mKO KP$^{-/-}$F PDAC mice (*Pdx-1-P2A-FlpO; Kras$^{FSF-G12D/+}$; Trp53$^{Frt/Frt}$; Ckm-Cre*; *Atg7$^{fl/fl}$*), as indicated. Columns of the heatmap were z-score normalized. (*n* = 11 control, *n* = 13 KP$^{-/-}$F PDAC, *n* = 9 *Atg7* mKO, and *n* = 11 *Atg7* mKO KP$^{-/-}$F PDAC mice, biologically independent samples.).

**Extended Data Figure 17: (Related to the Figure 5) Muscle-derived amino acids are redistributed to both tumor and host tissues in mice with PDAC. a-e.** Fraction of free amino acids that are $^{15}$N-labeled measured in the non-tumor head of pancreas (a), liver (b), small intestine (c), quadriceps (d), and soleus muscle (e) of 8-week-old male mice with KPC PDAC cells implanted in to the pancreas tail (PDAC), without or with deletion of *Atg7* in muscle (*Atg7* mKO) who had been fed an $^{15}$N-labeled *Spirulina* diet for 2-weeks with a chow diet for 4-weeks as described in Figure 5 (*n* = 4). **f-j.** Fraction of free amino acids that are $^{15}$N-labeled measured in the non-tumor head of pancreas (f), liver (g), small intestine (h), quadriceps (i) and soleus muscle (j) of 8-week-old male mice with pancreatic cancer cells implanted in to the pancreas tail (PDAC), without or with deletion of *Atg7* in muscle (*Atg7* mKO) who had been fed an $^{15}$N-labeled *Spirulina* diet for 2-weeks with a chow diet for 4-weeks as described in Figure 5 (*n* = 4). Statistical analysis was performed using unpaired two-sided *t*-tests, data are mean ± S.D and *n* represents the number of mice analyzed.

**Extended Data Figure 18 (Related to Figure 6): RNA-sequencing of PDAC tumors from mice without or with *Atg7* loss in the muscle**. **a-b.** Volcano plot showing normalized enrichment score (NES) versus $-\log_{10}$ adjusted P value from gene set enrichment analysis (GSEA) of PDAC tumors isolated from 6-week-old mice. Tumors were obtained from $KP^{-/-}F$ PDAC-bearing mice *(Pdx-1-P2A-FlpO; $Kras^{FSF\text{-}G12D/+}$; $Trp53^{Frt/Frt}$)* with or without muscle-specific loss of *Atg7* (*Pdx-1-P2A-FlpO; $Kras^{FSF\text{-}G12D/+}$; $Trp53^{Frt/Frt}$; Ckm-Cre; $Atg7^{fl/fl}$*; *Atg7* mKO $KP^{-/-}F$ PDAC). Gene set enrichment was performed using MSigDB Hallmark and immune gene signatures. (*n* = 5). **c-h.** Volcano plots showing $\log_2$ gene expression changes in PDAC tumors from *Atg7* mKO $KP^{-/-}F$ PDAC mice relative to $KP^{-/-}F$ PDAC tumors. Genes are grouped by functional categories, including extracellular matrix remodeling (c), focal adhesion (d), phagocytosis (e), and immune response pathways comprising response to chemokines (f), T cell activation (g), and Foxp3 target genes (h). (*n* = 5).

**Extended Data Figure 19: (Related to the Figure 6) Protein intake does not impact tumor growth or tissue size in mice with PDAC and loss of *Atg7* in muscle. a.** Schematic showing experimental design for determining whether diets with different amounts of intact protein impact organ size in mice with $KP^{-/-}F$ PDAC (*Pdx-1-P2A-FlpO; $Kras^{FSF\text{-}G12D/+}$; $Trp53^{Frt/Frt}$*) and loss of *Atg7* in muscle (*Pdx-1-P2A-FlpO; $Kras^{FSF\text{-}G12D/+}$; $Trp53^{Frt/Frt}$; Ckm-Cre; $Atg7^{fl/fl}$*; *Atg7* mKO $KP^{-/-}F$ PDAC). **b-f.** Whole-body weight (b), and weight of liver (c), perigonadal and subcutaneous ATs (d), pancreas (e), quadriceps, gastrocnemius, and soleus muscle (f) normalized to total body weight of 4-week-old mice who were placed on diets with a protein amount found in a AIN-

93G purified diet (1x) or a diet with 0.5X protein or 2X protein as indicated and followed until endpoint (*n* = 7 *Atg7* mKO KP$^{-/-}$F PDAC on 0.5X Protein, *n* = 11 *Atg7* mKO KP$^{-/-}$F PDAC on 1X Protein, and *n* = 15 *Atg7* mKO KP$^{-/-}$F PDAC on 2X Protein). Both male and female mice were used. Statistical analyses were performed using one-way ANOVA with Tukey's post hoc test, data are mean ± S.D and *n* represents the number of mice analyzed.

#### Extended Data Fig.1

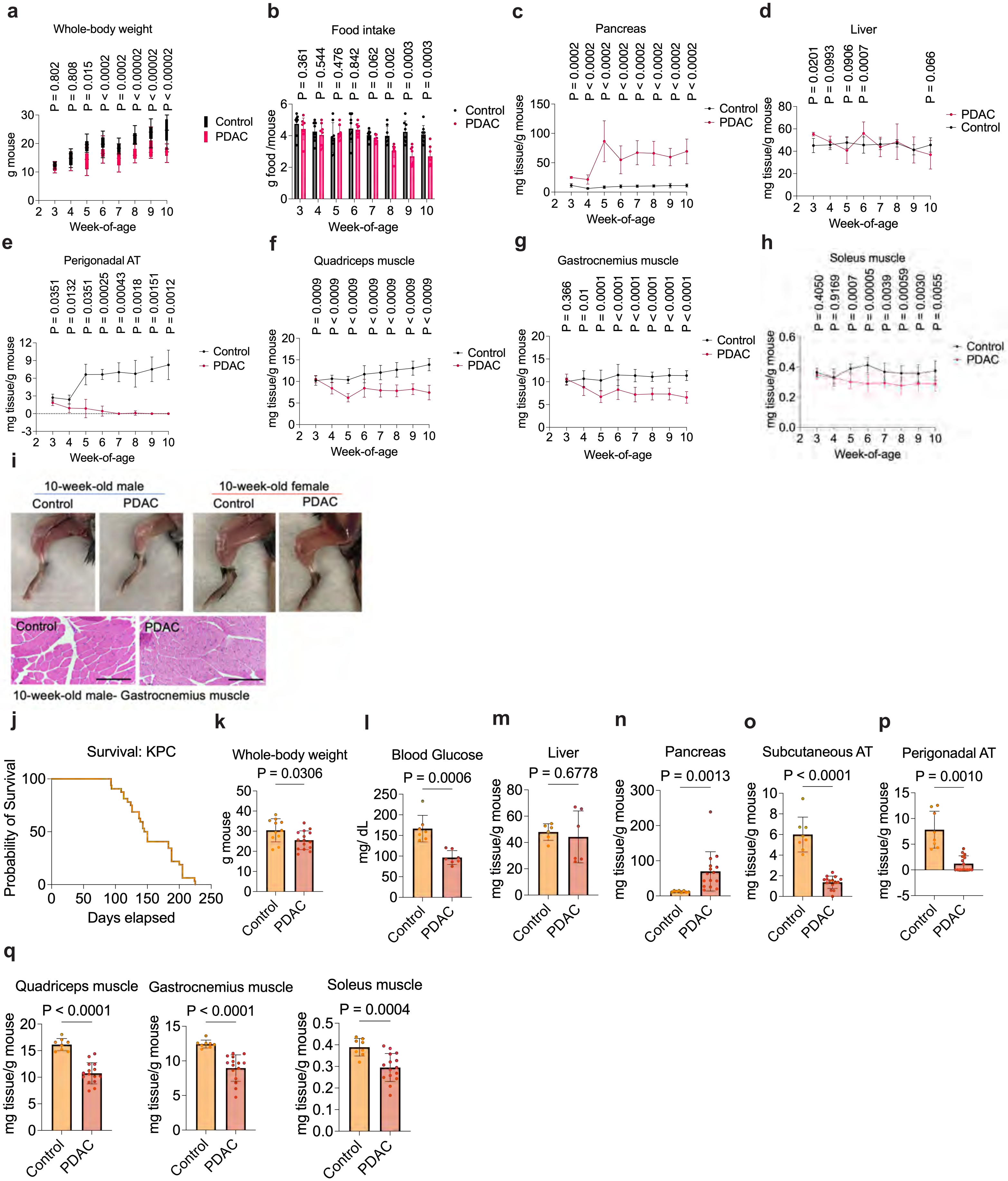

#### Extended Data Fig. 2

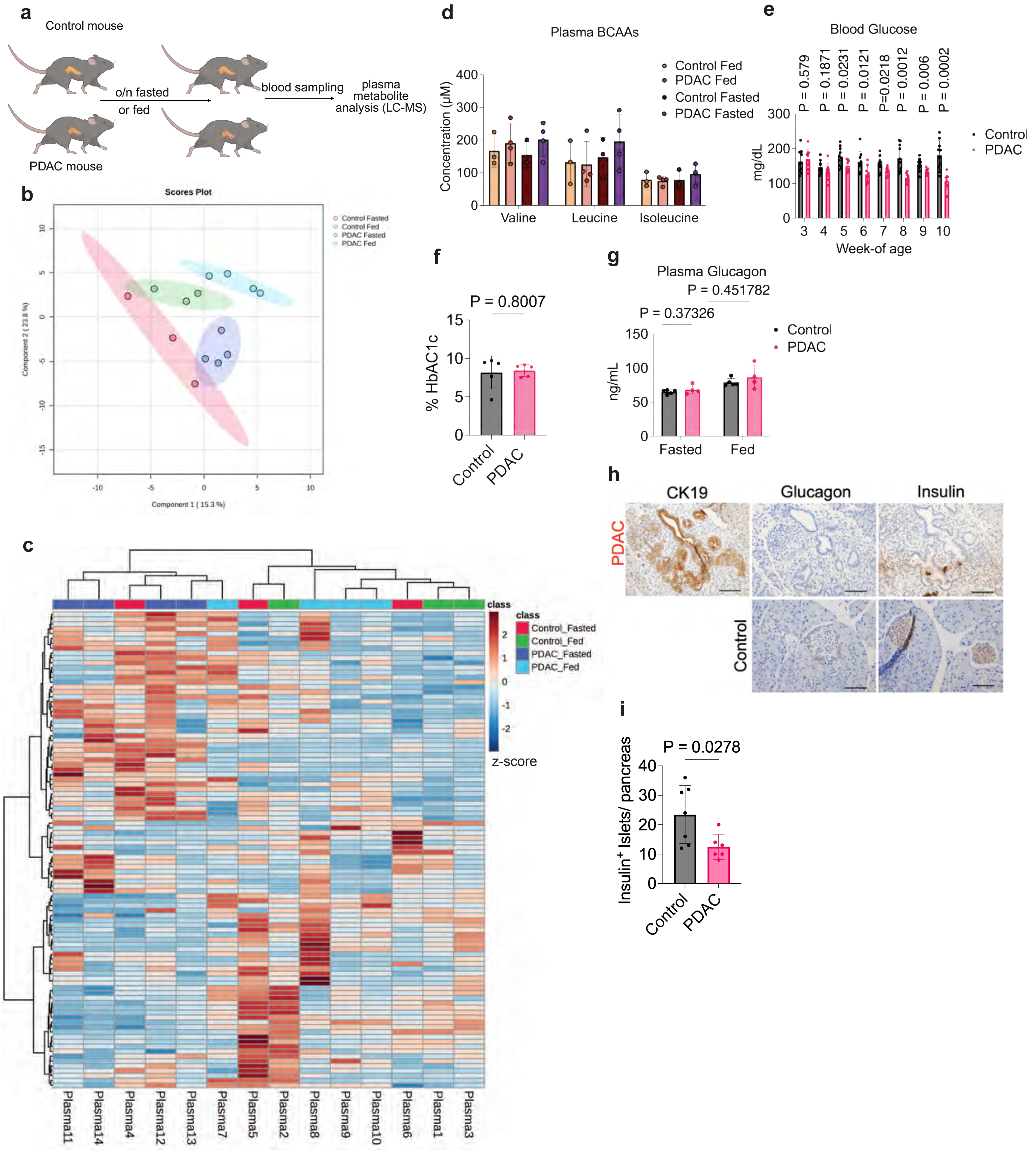

#### Extended Data Fig.3

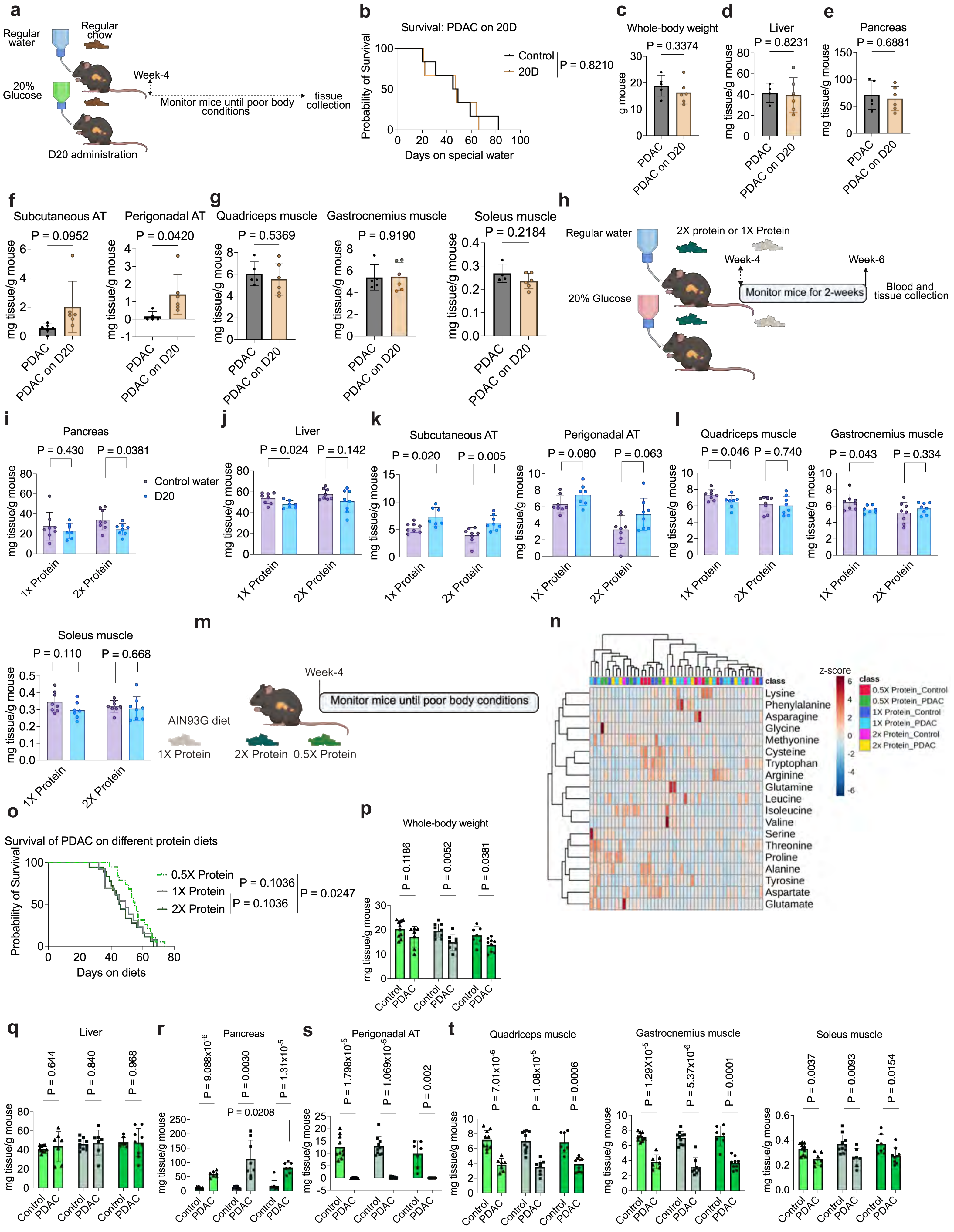

#### Extended Data Fig. 4

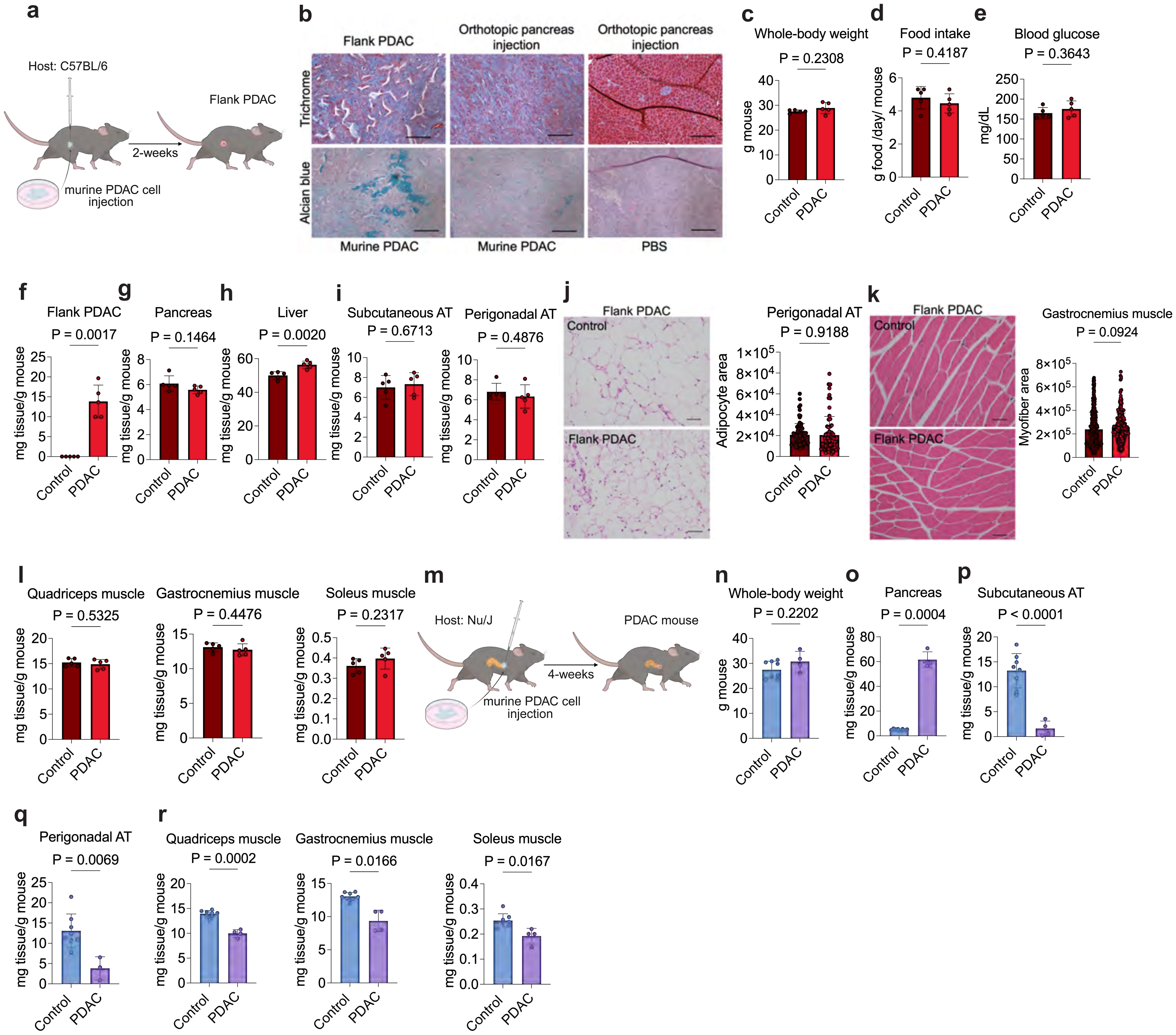

#### Extended Data Fig. 5

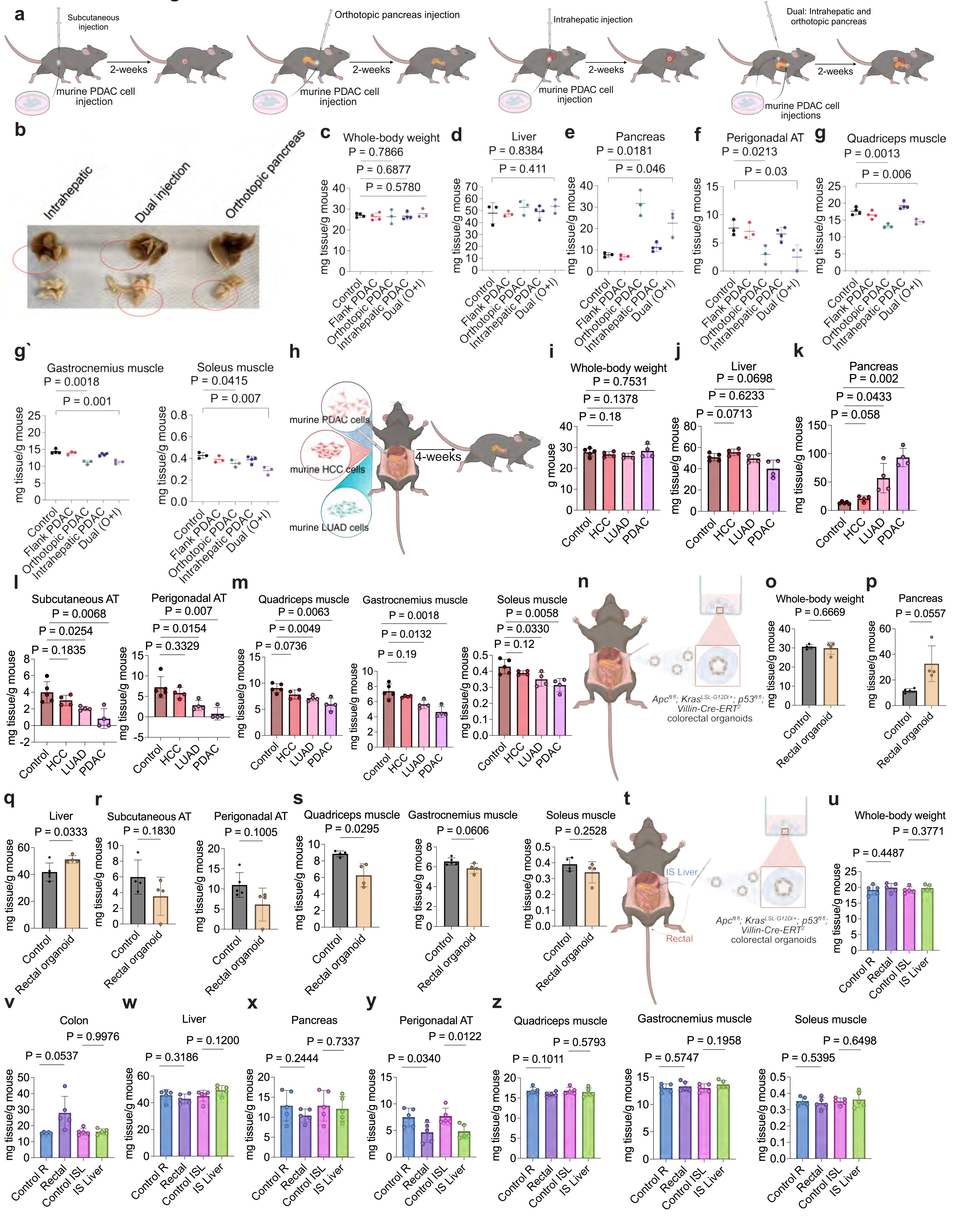

### Extended Data Fig. 6

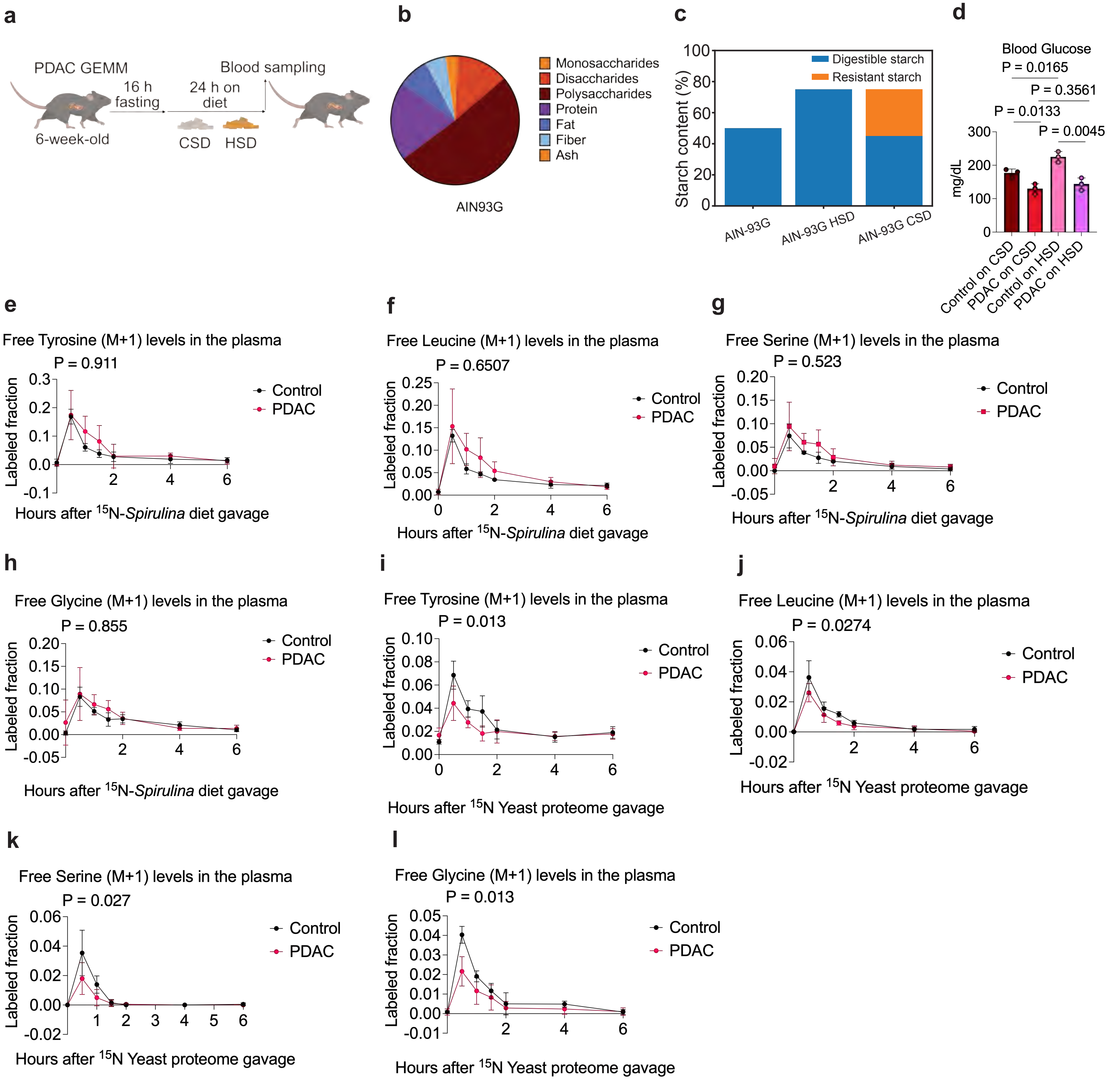

### Extended Data Fig. 7

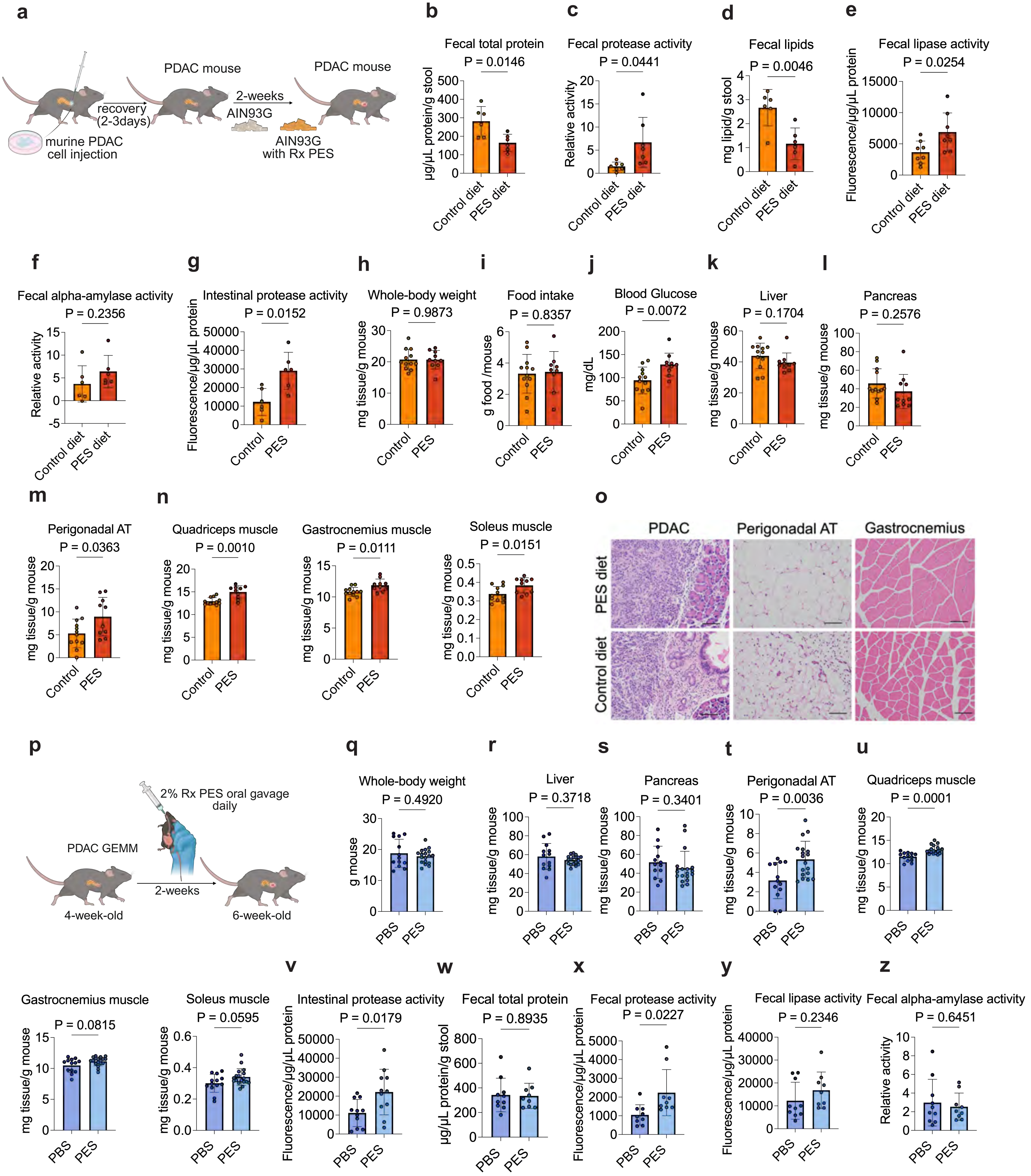

#### Extended Data Fig. 8

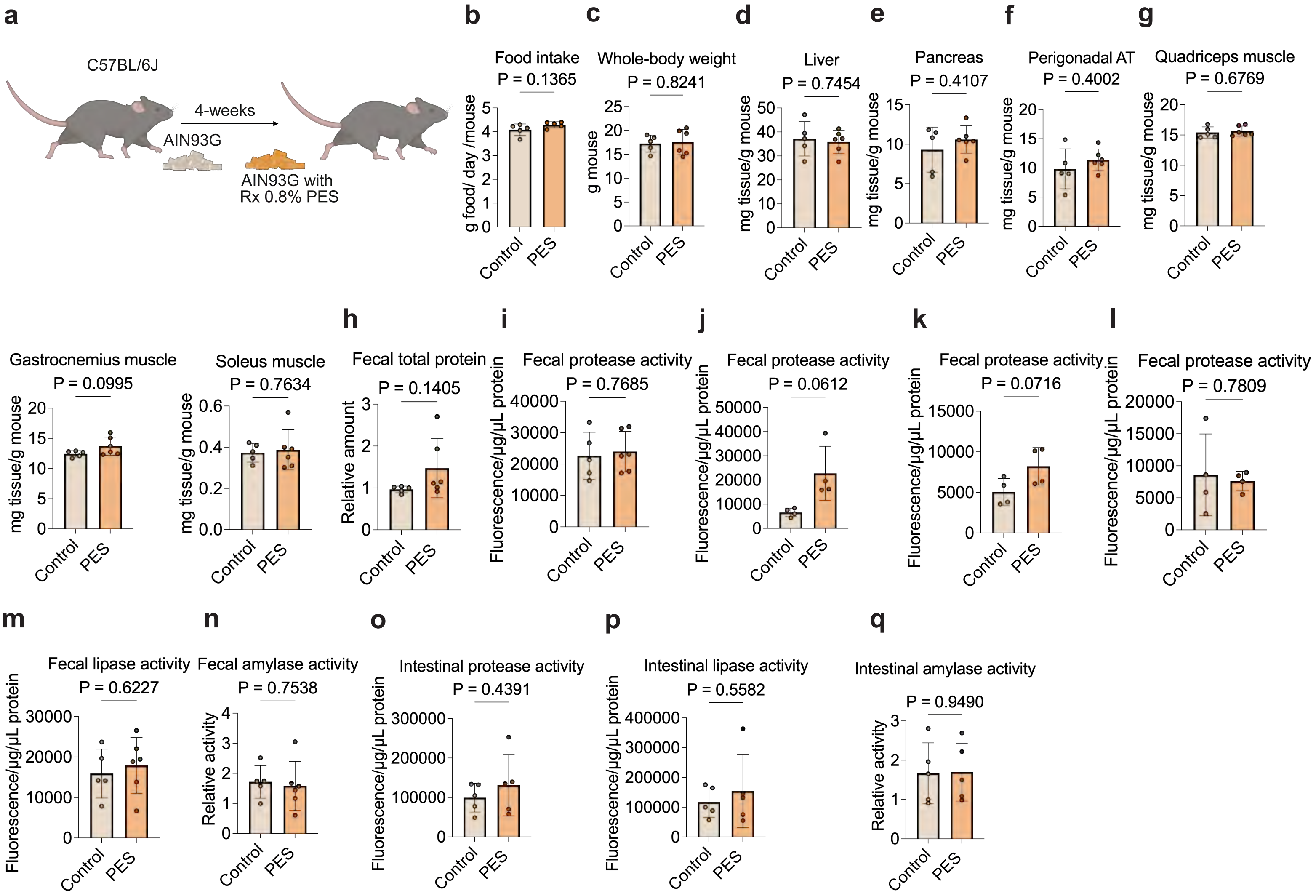

#### Extended Data Fig. 9

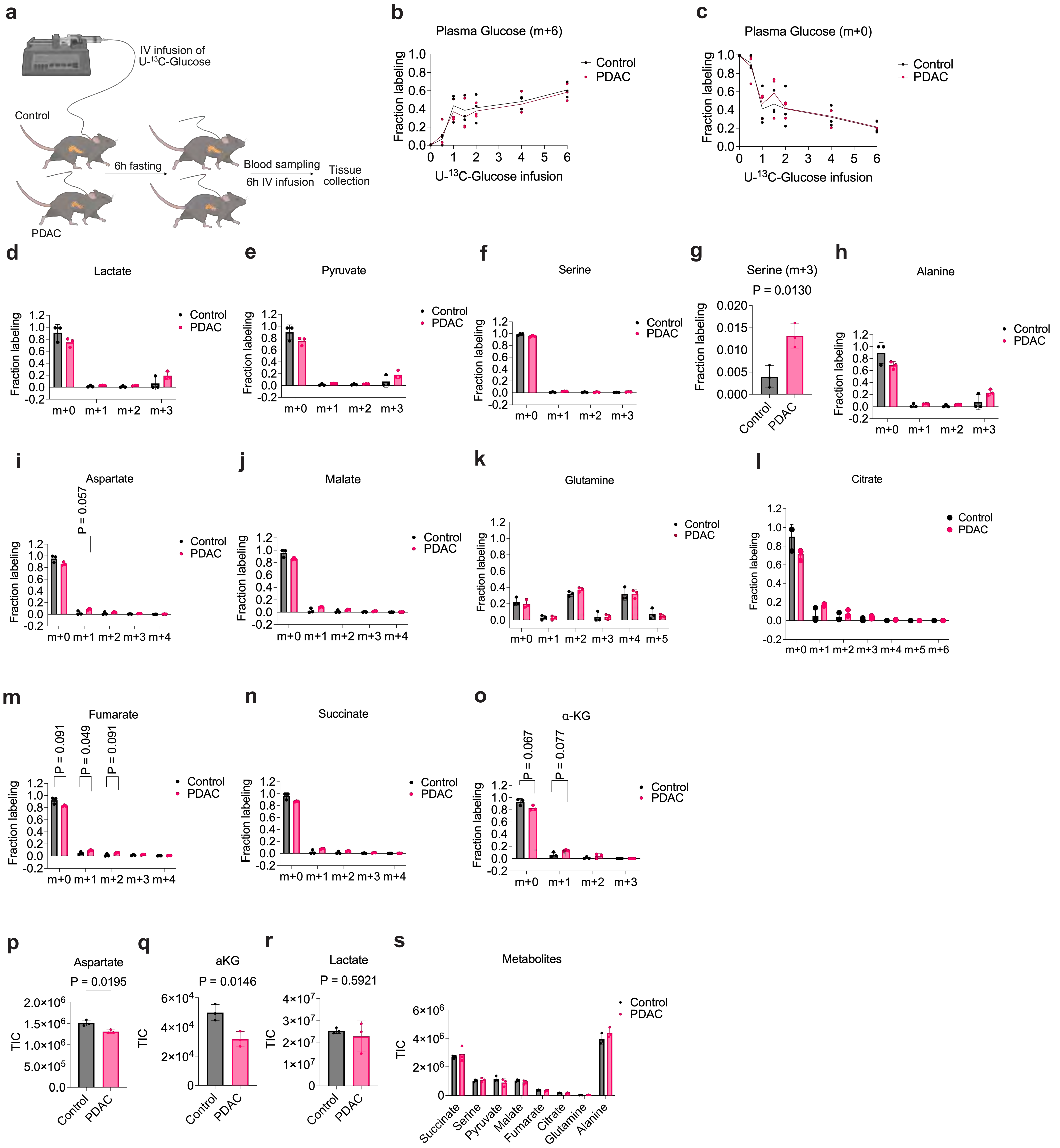

#### Extended Data Fig. 10

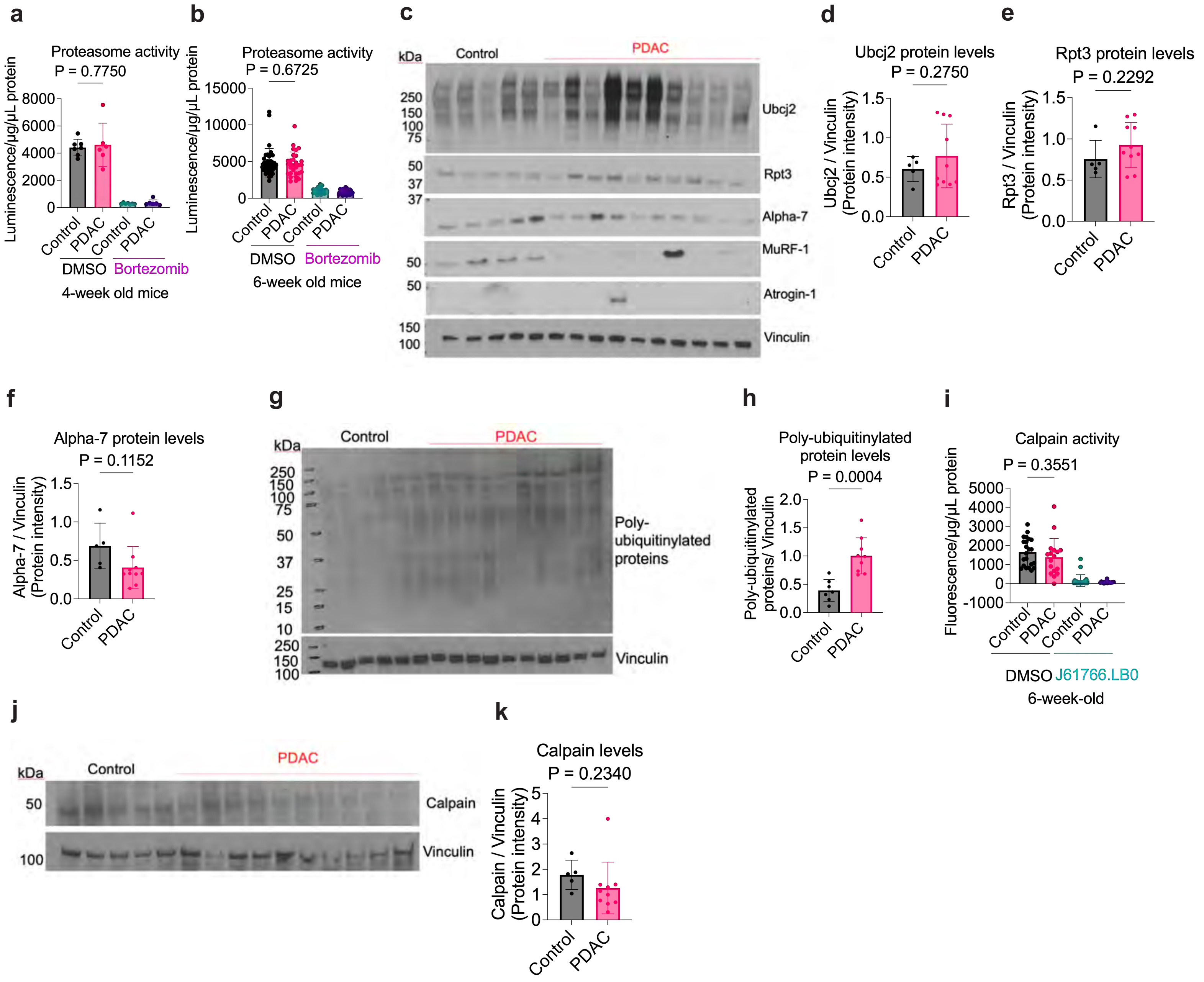

#### Extended Data Fig. 11

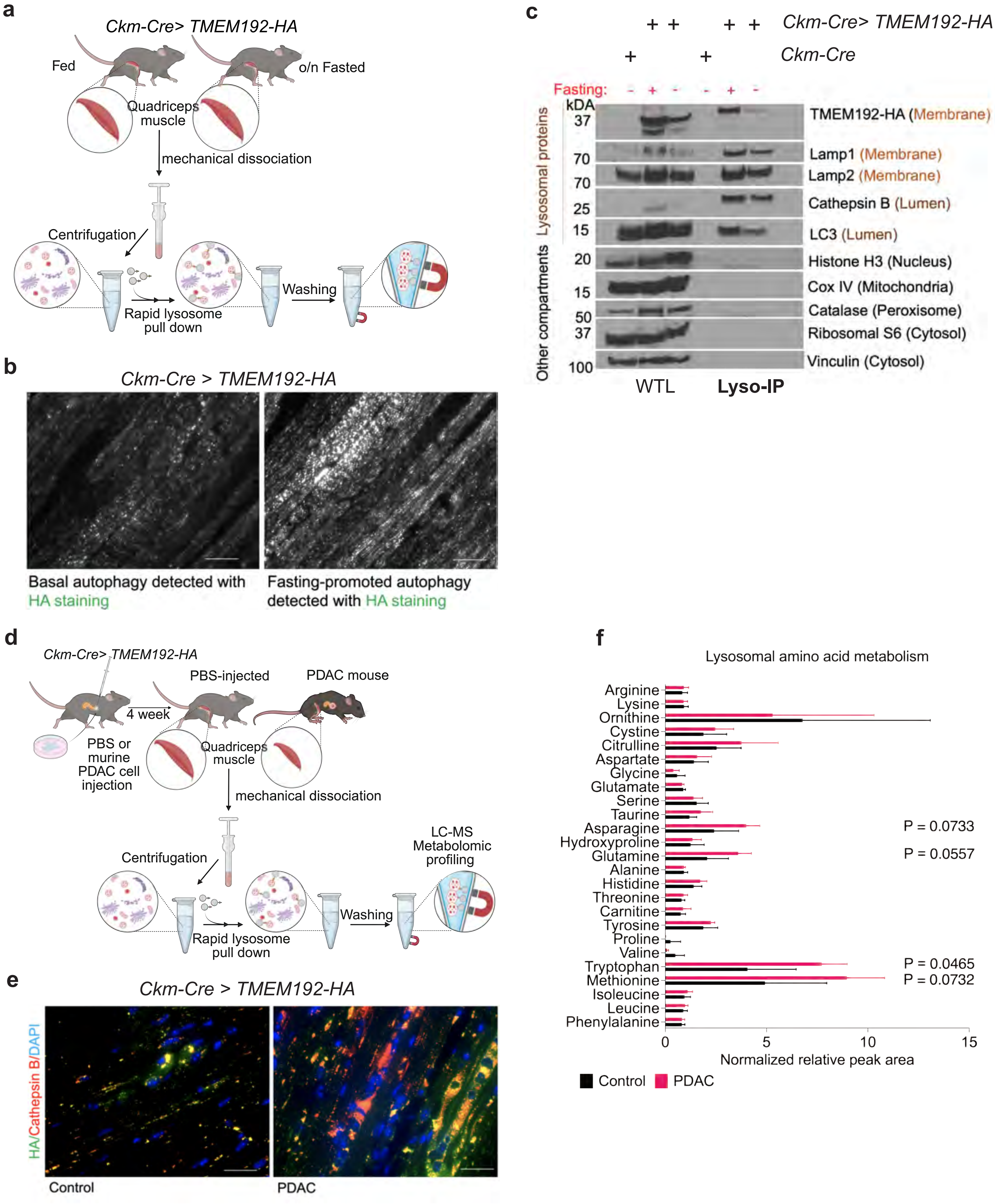

#### Extended Data Fig. 12

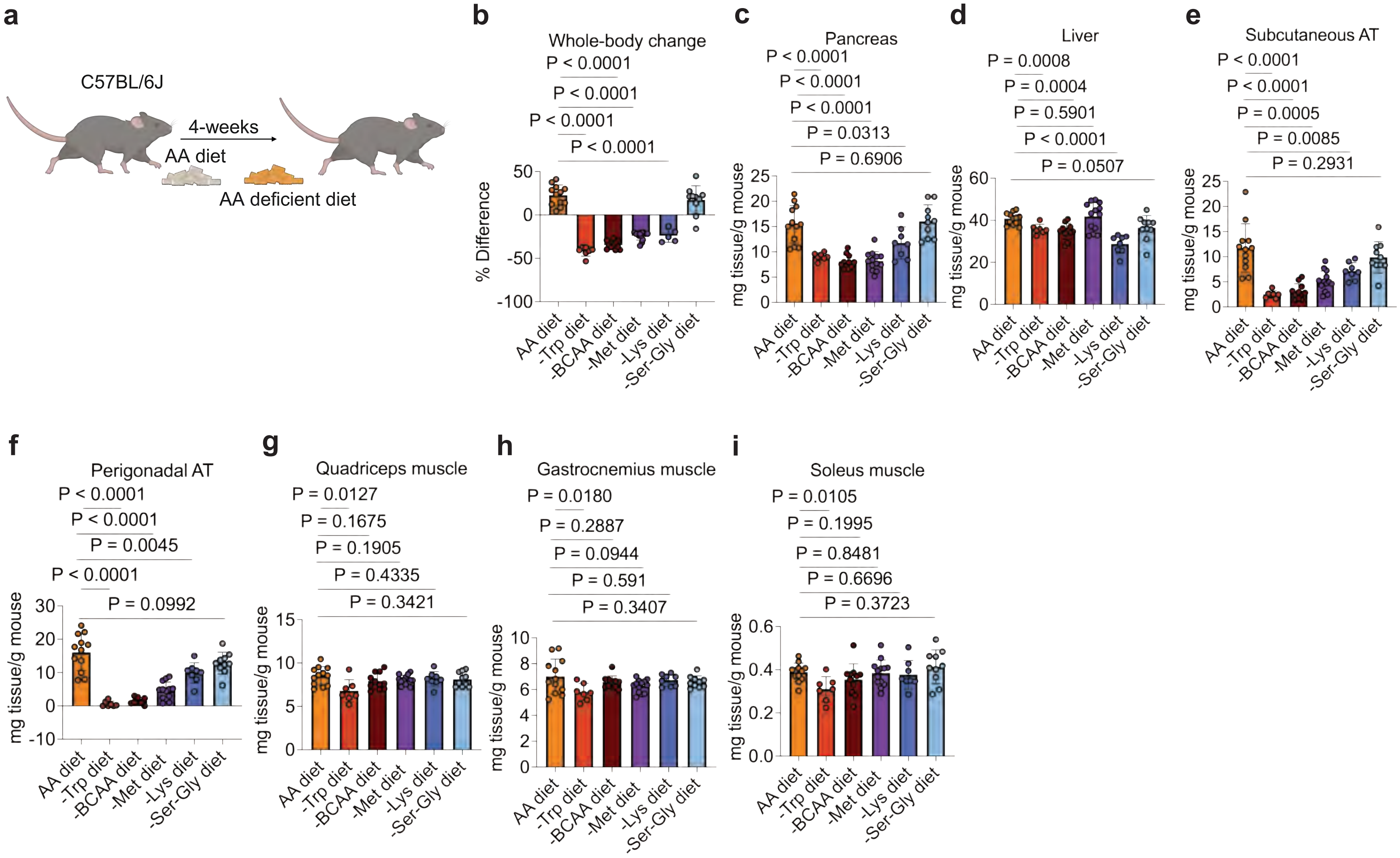

#### Extended Data Fig. 13

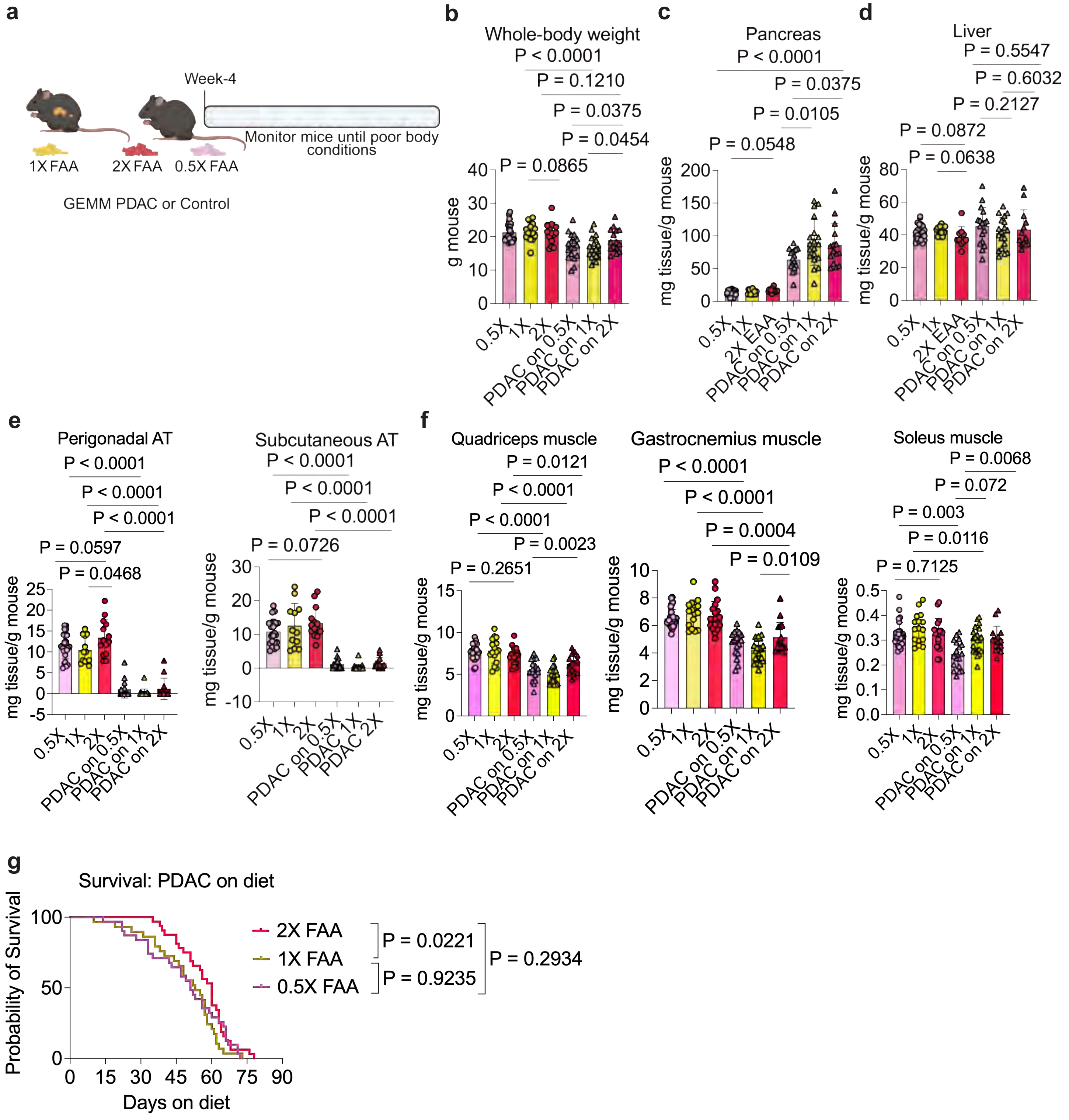

#### Extended Data Fig. 14

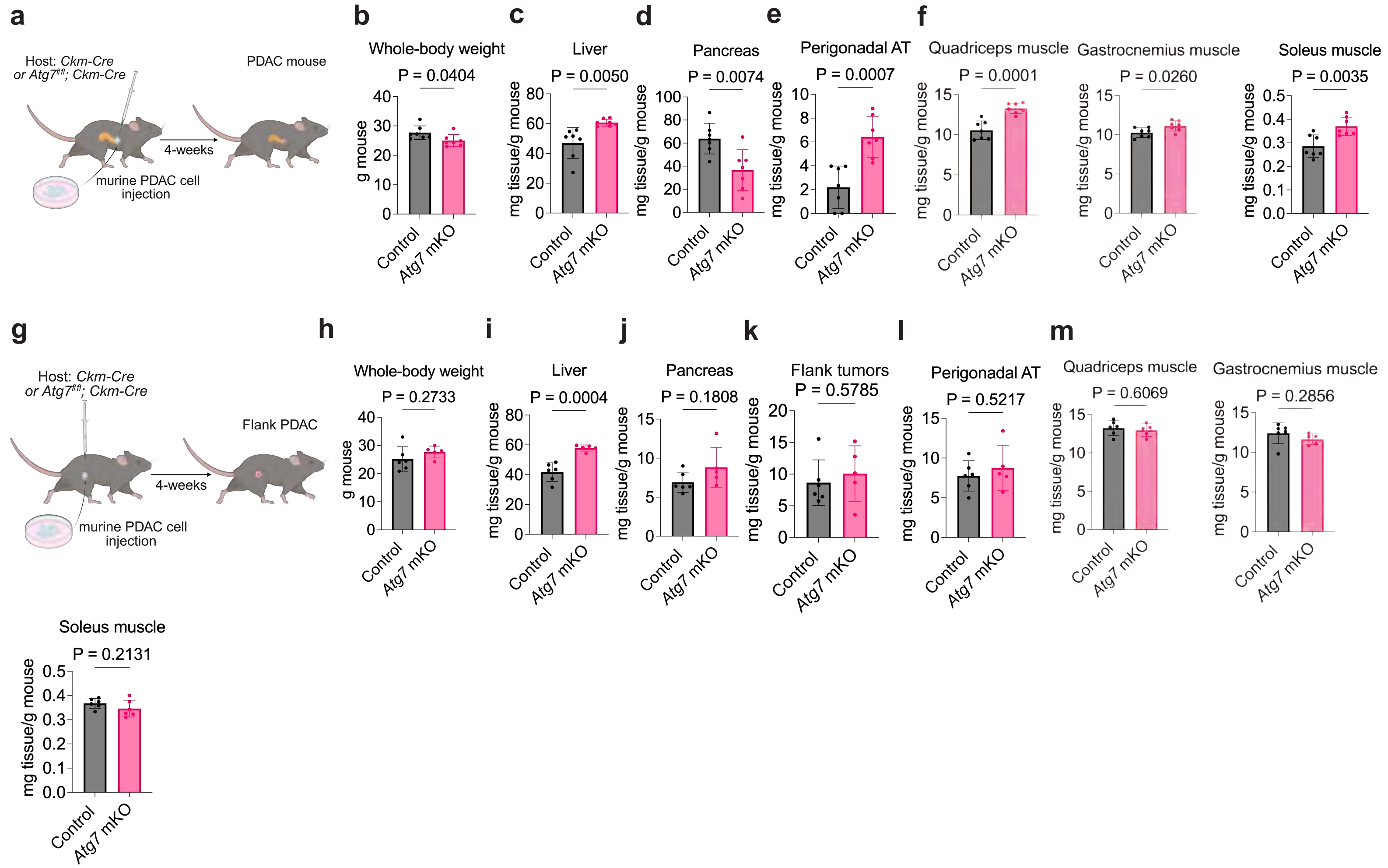

#### Extended Data Fig. 15

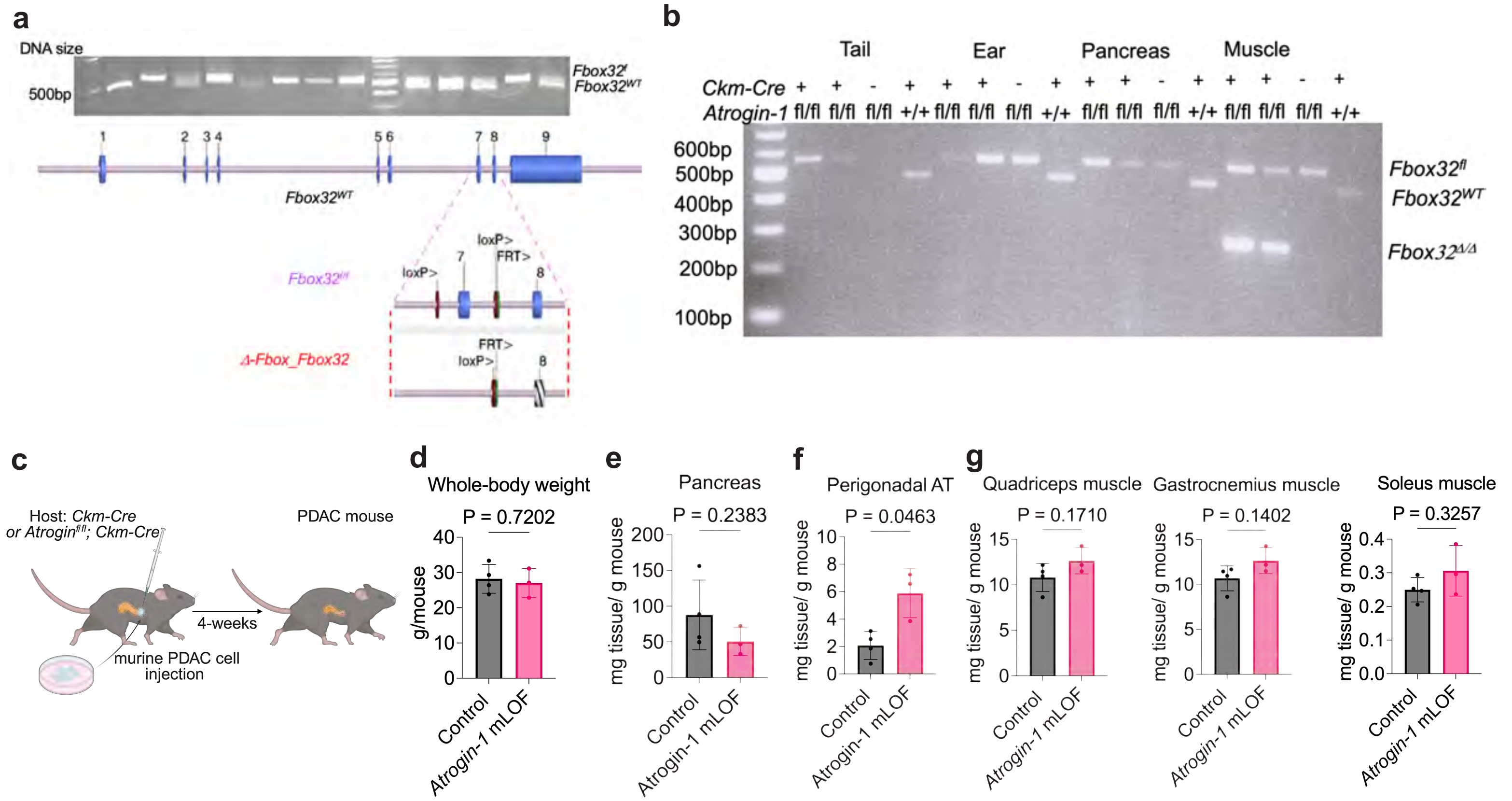

#### Extended Data Fig. 16

**a**

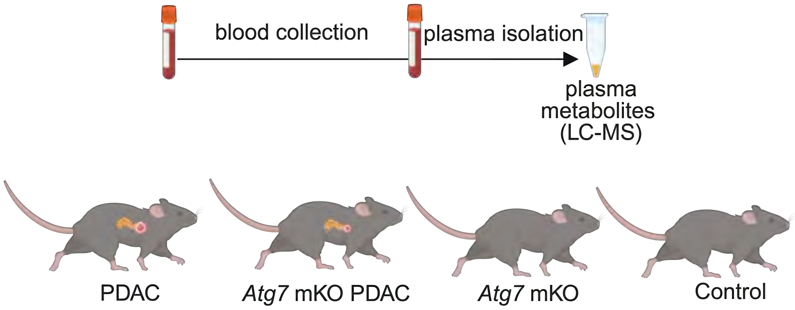


**b**

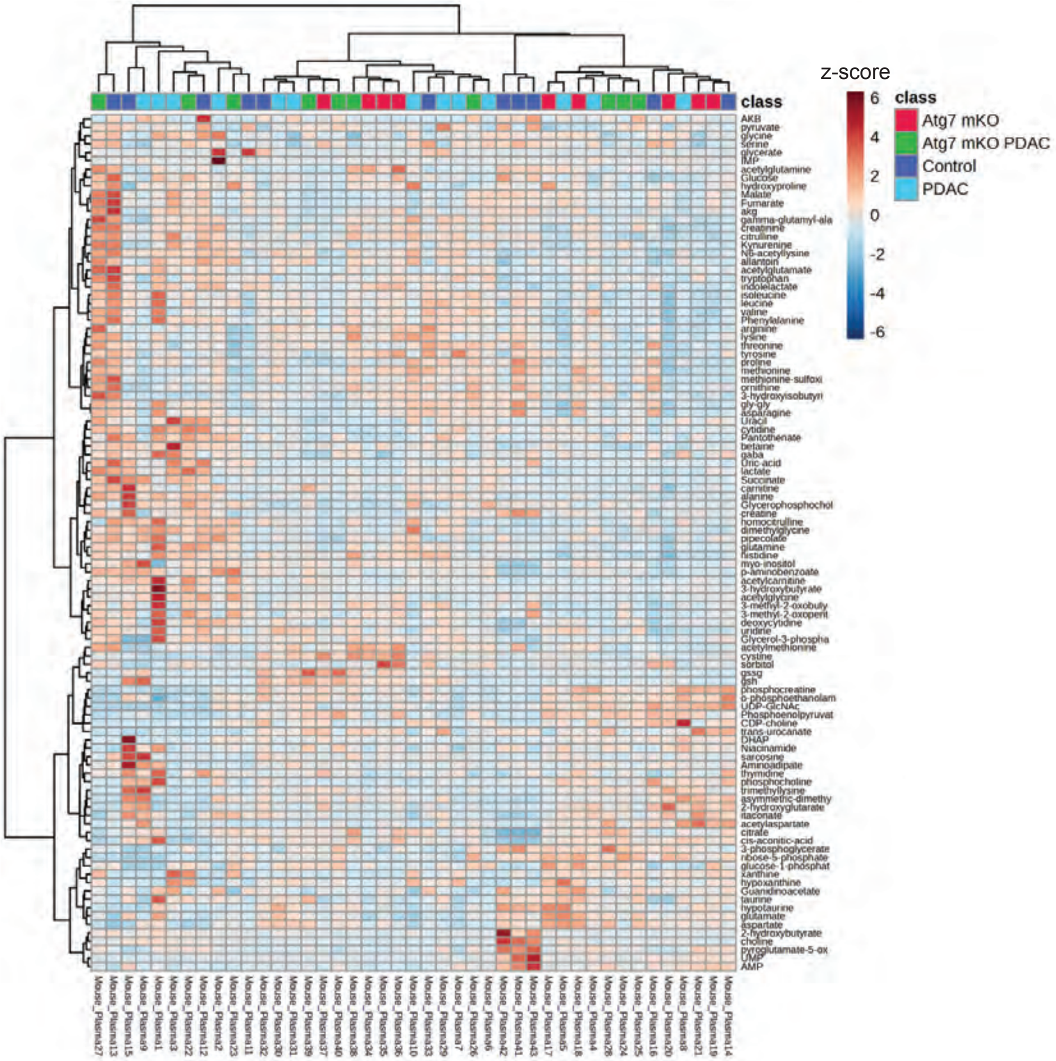

#### Extended Data Fig. 17

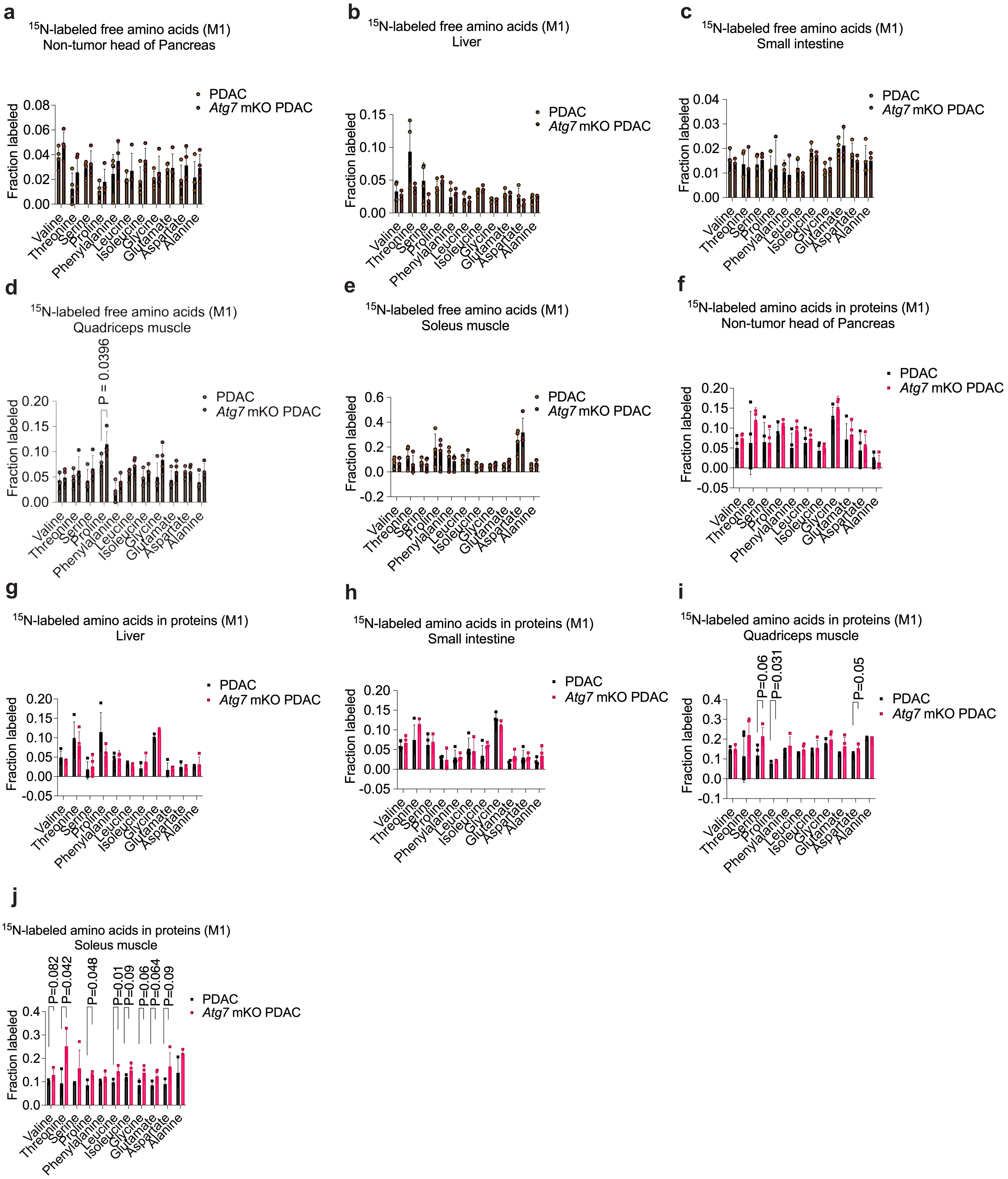

#### Extended Data Fig. 18

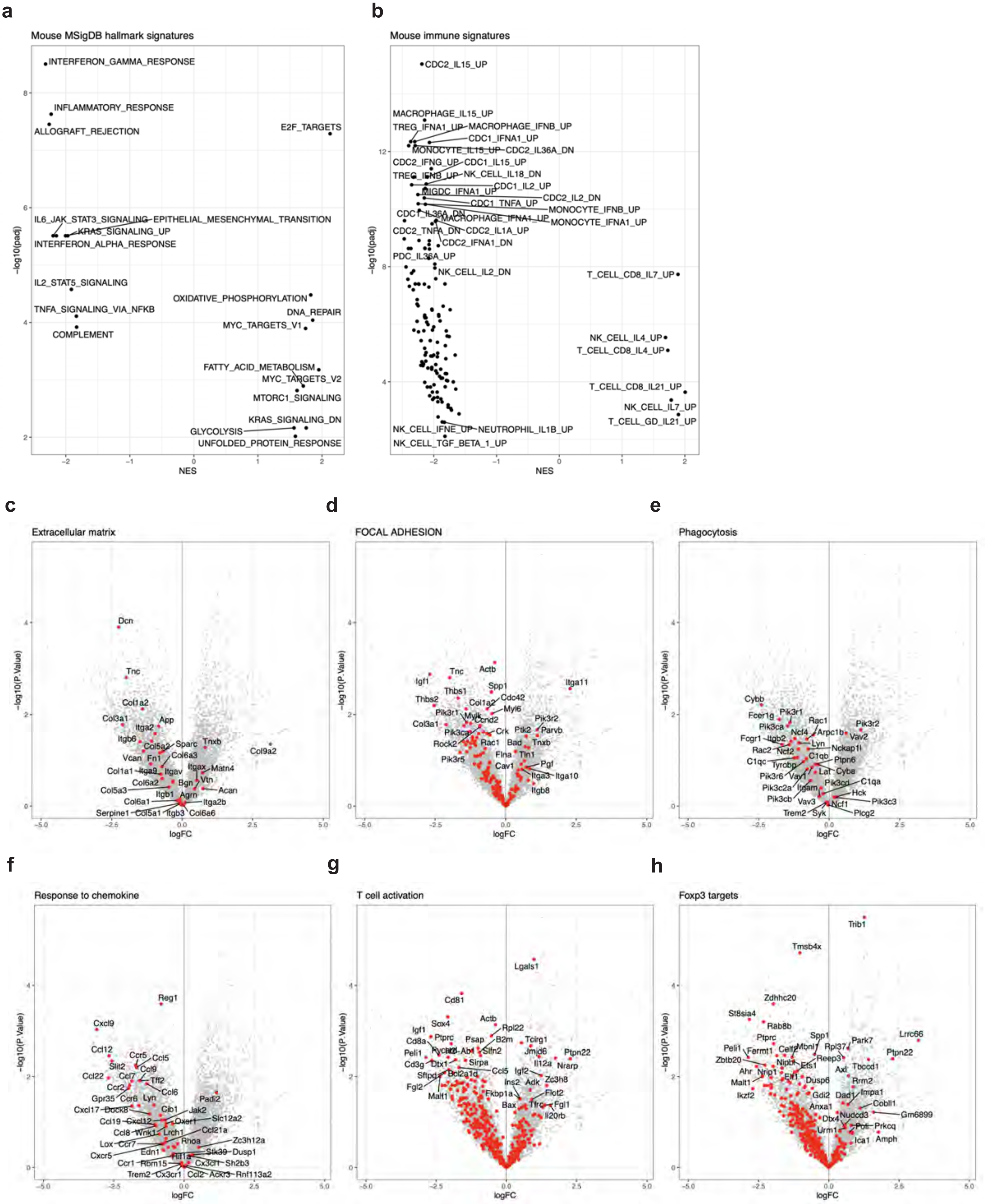

#### Extended Data Fig. 19

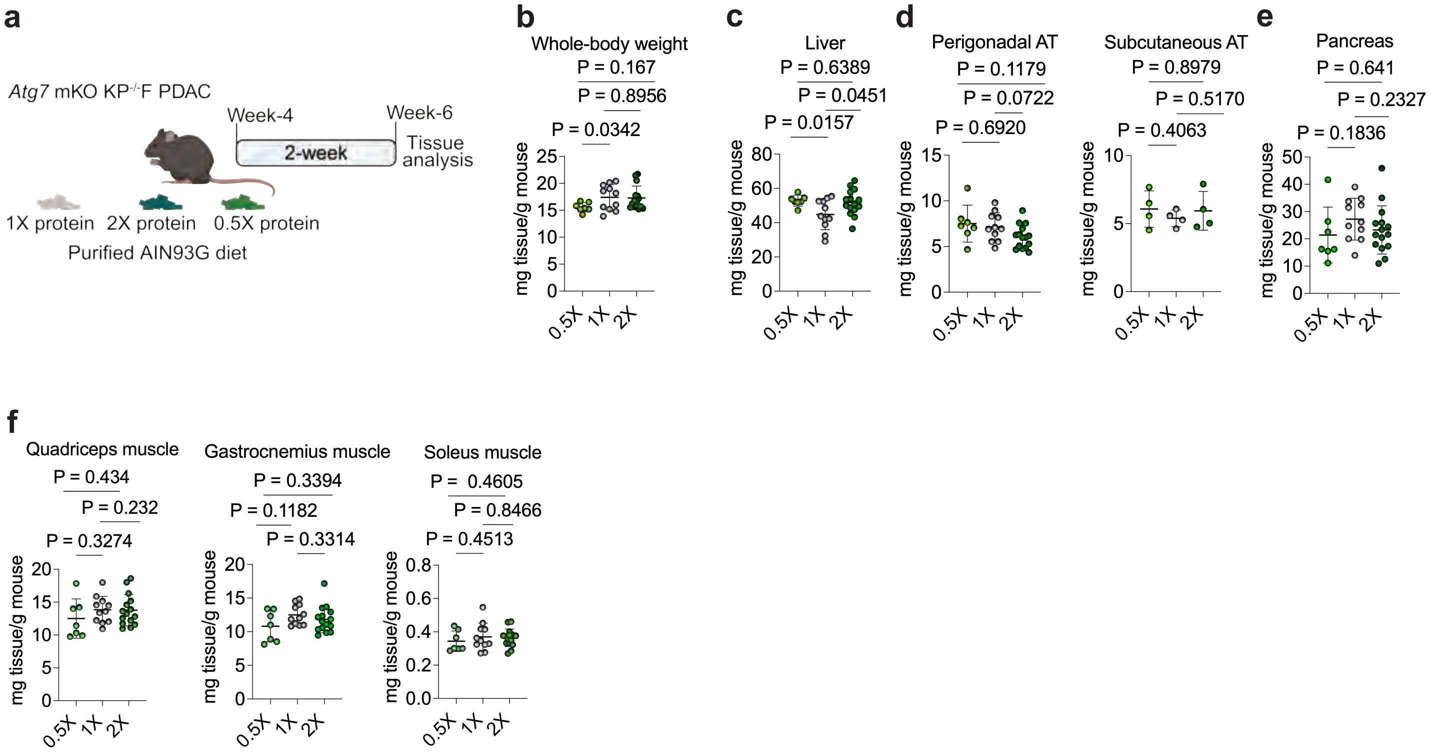
